# Programming human cell type-specific gene expression via an atlas of AI-designed enhancers

**DOI:** 10.1101/2025.09.30.679565

**Authors:** Sebastian M. Castillo-Hair, Christopher H. Yin, Leah VandenBosch, Timothy J. Cherry, Wouter Meuleman, Georg Seelig

## Abstract

Differentially active enhancers are key drivers of cell type specific gene expression. Active enhancers are found in open chromatin, which can be mapped at genome scale across tissue and cell types. Though incompletely understood, the relationship between chromatin accessibility and enhancer activity has been exploited to identify, model, and even design functional enhancers for selected cell types, but to what extent this design strategy can generalize across human cell and tissue types remains unclear. Here, we trained deep neural networks on a large corpus of chromatin accessibility data from hundreds of human biosamples. We used these models to generate an atlas of tens of thousands of synthetic enhancers, targeting hundreds of cell lines, tissues, and differentiation states, aiming to maximize accessibility in target samples and minimize it in all off-target ones. Experimental testing of thousands of designs in a representative subset of ten human cell types and in mouse retina demonstrated their function as specific enhancers, not only in the case of one-versus-all objectives but also when targeting two or three cell types. An explainable AI analysis, enabled by our large-scale enhancer measurements, allowed us to identify similarities and differences between the sequence grammar underlying accessibility and enhancer activity. Our results show that model-guided design of enhancers can help us decipher the cis-regulatory code governing cell type specificity and generate novel tools for selective targeting of human cell states.

## Main

Across tissues, developmental stages, and disease conditions, cells adopt distinct states characterized by unique molecular profiles^1^. Engineered molecular systems that interface with these states hold transformative potential for biotechnology, with applications that include targeted gene therapies^2^ and programmed differentiation for regenerative medicine^3–5^. However, while engineered DNA-^6–10^ and RNA-encoded^11–14^ programs with specificity towards individual cell types have been demonstrated in select contexts, strategies to generalize these to any human cell type or state of interest remain underdeveloped.

A prevalent mechanism for establishing cell type identity during development is the sequential activation of transcription factors (TFs) and their matching enhancers – non-coding DNA elements that recruit TFs and direct RNA polymerase II to initiate transcription at cognate promoters^15^. Their role as drivers of cell type identity makes enhancers attractive as cell state sensors and as tools for targeting transgene expression to cell types of interest. Active enhancers adopt specific epigenomic configurations, such as chromatin accessibility and histone modifications^16^, which can be profiled genome-wide across diverse samples^17,18^. These datasets provide an opportunity for selecting DNA fragments with desired accessibility profiles in the hope that they will act as enhancers driving gene expression to a target cell type. However, such genomic fragments have an uncertain success rate when tested for enhancer activity in isolation^19^, necessitating extensive pre-screening and validation^20,21^. Underlying key limitations include our incomplete understanding of the “grammatical” rules by which enhancer sequence determines activity, further complicated by cooperative, sometimes counterintuitive TF interactions^22^, and uncharacterized differences between the sequence grammar of genomic accessibility compared to enhancer function, due for example to pioneer factors and other chromatin remodelers that may not directly drive transcription^23,24^. Massively parallel reporter assays (MPRAs) allow for large scale screening of functional enhancers^25,26^, but they have limited power to resolve cell type-specific regulation because few cell types or multi-cellular systems are compatible with delivery of large DNA libraries. Single-cell MPRAs^27,28^ can cover a larger variety of cell types in heterogeneous samples, but are still limited by library delivery and are restricted to library sizes of a few hundred sequences^29^.

Recently, Artificial Intelligence (AI) models have been trained on large scale genomic^30–41^ or MPRA-derived^42–45^ datasets and used to make inferences about cis-regulatory function^46,47^. AI model-guided sequence design can generate cis-regulatory elements with superior properties than those obtained from screening natural sequences, because such models can interpolate between measurements and potentially generalize to sequences and conditions not seen during training^6–10,14,42,43,48,49^. In particular, neural network (NN) predictors of bulk^6^ or single-cell^7,9,10^ accessibility have been recently used for enhancer design, with the expectation that learned accessibility grammar extrapolates to enhancer activity. This approach has been so far successfully demonstrated in limited contexts, including two cell types in the developing fly brain^7^, three endothelial and muscle-related cells in zebrafish embryos^10^, and in human melanoma^7^, liver, and immune cell lines^6^. However, its broader potential—leveraging the vast diversity of accessibility datasets to design enhancers for any human tissue of interest—remains largely unexplored.

Here, we show that NN predictors of genomic accessibility can be used to design functional enhancers for human cell and tissue types from a wide range of organ systems and developmental origins (**Figure 1A**). As training data, we use the ENCODE DHS Index^50^, a genomic atlas containing accessibility measurements from 3.59 million genomic DNase I Hypersensitive Sites (DHSs) across 733 human biosamples that include tissues, primary cells, and cell lines. We initially train models and design sequences with cell type-specific accessibility targeted to 64 distinct samples. We experimentally validate thousands of enhancer candidates in 10 cell lines of diverse tissue origin, which were included in the 64 target samples and modeled on an equal footing with primary cells and tissue samples – their most relevant distinguishing feature being that they are amenable to MPRA experiments. To further validate performance in a multicellular biological system, we characterize synthetic enhancers in mouse retina. Furthermore, we design functional synthetic enhancers with tunable target expression and with multiple cell types as targets, demonstrating the ability to generalize to design targets not found in nature. Interpretation of accessibility and enhancer activity models, enabled by the scale of our MPRA validation experiments, reveals key differences in the sequence grammar of these processes. Finally, we extend our modeling approach to all 733 samples in the DHS Index, resulting in a foundational model that can be used to design enhancers for a large variety of human cell states. We use this model to generate an atlas of 52,200 AI-designed human enhancers targeting every unique sample in the DHS Index, and highlight a set of enhancers with differentiation stage-specific activity during stem cell differentiation into cardiomyocytes. Our work demonstrates the potential of AI to design cell type and state-sensing elements, provides the largest repository of validated enhancers to date, and sheds light on generalizable regulatory rules to design enhancers from accessibility predictors.

**Figure 1.**
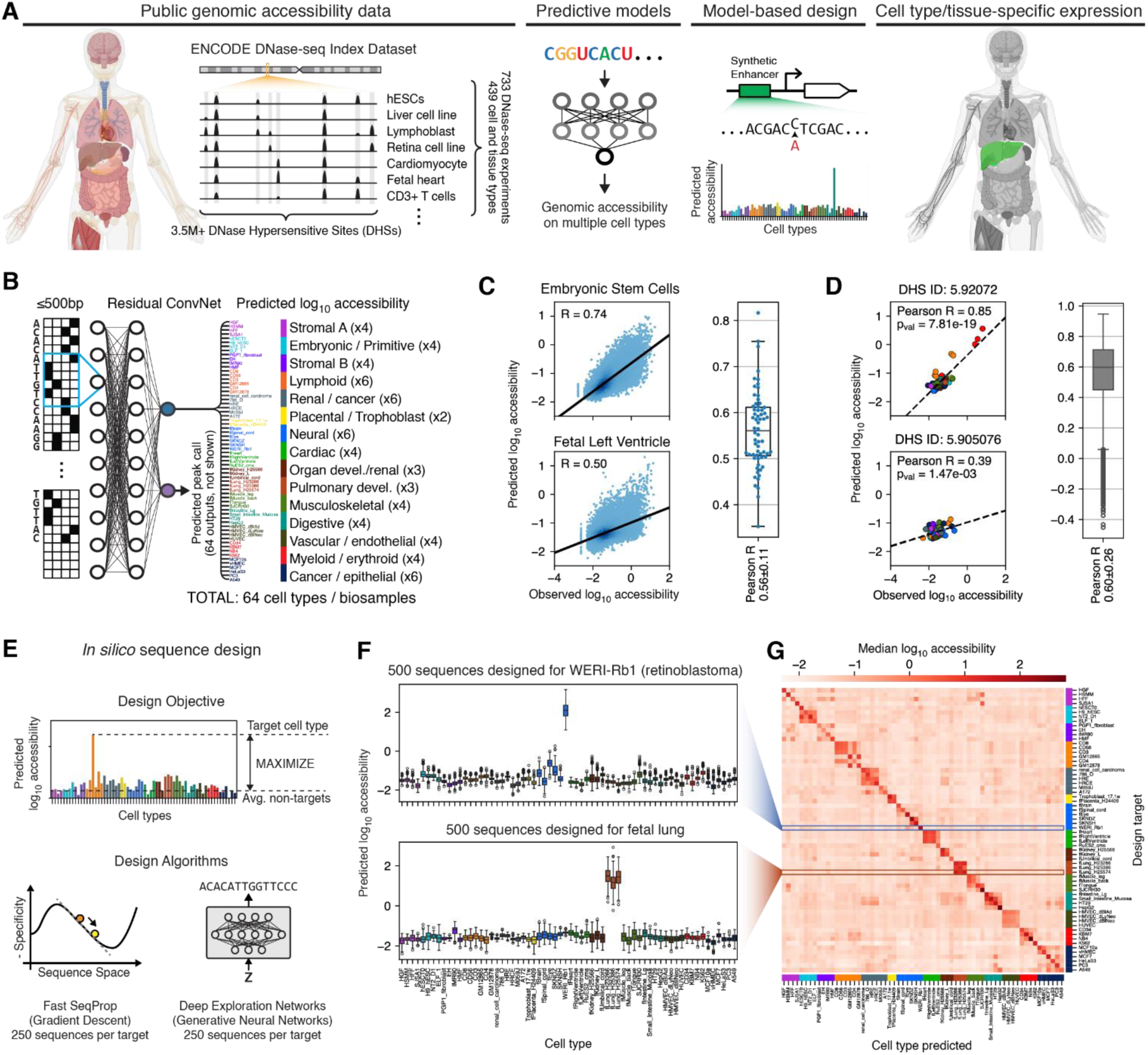
Neural network models of genomic accessibility enable programming cell type-specific gene expression. **(A)** Outline of the sequence design method. NN models trained on public genomic accessibility data are used to design enhancers with cell and tissue type-specific activity. **(B)** High-level schematic of the DHS64 model. Predicted output cell types are colored and categorized by their associated “DHS component”^50^, roughly corresponding to distinct originating tissue types within the DHS Index (**Methods**). See **Supplementary Figure 1** for a more detailed depiction of the model architecture, and **Supplementary Table 1** for the list of modeled biosamples (i.e. cell types). **(C)** Per-biosample model performance. Left: examples of regression performance for two biosamples. Right: Distribution of regression (Pearson R) performance across all 64 biosamples. Numbers in the x axis indicate median ± interquartile range. **(D)** Per-DHS model performance. Left: performance on two example DHSs, where each dot corresponds to a biosample. Right: Pearson R distribution across all DHSs in the test set. Numbers in the x axis labels indicate median ± interquartile range. In **(C)** and **(D)**, we evaluate the model against a test set of held-out DHSs active in ≤10 biosamples and with enhancer-like genomic and chromatin annotations (**Methods**). **(E)** Sequence design procedure, illustrating the specificity objective function and the two NN-based sequence generation methods. **(F)** Predicted accessibility signals for 500 sequences (250 generated with Fast SeqProp + 250 with DEN) designed to be specific to WERI-Rb-1 (a retinoblastoma cell line, top) and a fetal lung tissue (bottom). Predictions were obtained using a separately trained model from the ones used during sequence design (**Methods**). **(G)** Predicted accessibility of sequences targeted to each modeled biosample, with rows showing the median of 500 sequences per target.

### NN models predict genomic accessibility across multiple cell types

We first trained NN models to predict genomic accessibility across cell and tissue types. Previously, the DHS Index data was found to decompose into 16 biological “components”: 15 strongly associated with samples of different tissue origins, and one component corresponding to broadly accessible (i.e. non-specific) DHSs. We initially selected 64 biosamples spanning all 15 tissue-specific components and with high DNase-seq quality metrics. Selected samples included 18 tissues (e.g. fetal heart, fetal lung), 17 primary cells (e.g. CD3 and CD34 cells), 28 cell lines, and one *in vitro* differentiated cell sample according to ENCODE’s classification (**Supplementary Table 1**, **Methods**).

State-of-the-art large transformer models can predict multiple epigenomic features such as accessibility from >100kb sequences^32,37^, but their performance on cell type-specific regulatory elements is limited^51^ and their size make them difficult to use with sequence design methods that require repeated model and gradient evaluations^52,53^. Thus, our initial model, “DHS64” used an ensemble of deep residual neural networks to predict continuous accessibility signals (i.e. normalized log_10_ DNase-seq read density) and peak call probabilities for all 64 biosamples, given an input sequence of up to 500bp (**Figure 1B**, **Supplementary Figure 1A, Methods**). To better assess the ability of our model to capture cell type-specific activity, we evaluated performance on a filtered set of highly specific DHSs with enhancer-like chromatin and genomic features, and found that filtering the training data for DHSs active in 10 or fewer biosamples resulted in best performance (**Supplementary Figure 1B-G, Methods**). For each biosample, DHS64 predicted accessibility signals with a Pearson R of 0.56±0.11 (median ± interquartile range) and peak call probabilities across all accessible regions in the test set with an area under the precision-recall curve (AUPRC) of 0.49±0.13 (**Figure 1C**, **Supplementary Figure 1H-I, Supplementary Table 3**). Example predictions are shown for embryonic stem cells and fetal left ventricle. Furthermore, DHS64 captured differences in accessibility across biosamples for any given DHS with a Pearson R of 0.60±0.26 (**Figure 1D**, **Supplementary Figure 1J)**. Finally, we confirmed that DHS64 accurately predicted the accessibility levels of the most cell type-specific DHSs in the test set (**Supplementary Figure 1K-L**), further confirming that the model makes good predictions on the types of sequences we intend to design.

### Generative sequence design of putative cell type-specific enhancers

We next used DHS64 to generate synthetic sequences with cell type-specific accessibility. Specifically, we sought to maximize the difference between predicted log_10_ accessibility in a target biosample and the average log_10_ accessibility across all others (**Figure 1E**). We used Fast SeqProp^52^, which optimizes sequences via gradient descent, to design 250 sequences with a length of 145 nt targeted to each of the 64 modeled biosamples. Similarly, we used Deep Exploration Networks (DENs)^53^ – generative NNs that produce sequences with high predicted performance and low pairwise similarity – and generated 250 additional sequences per target. To prevent overfitting to a single model, sequence optimization was performed using a mini-ensemble of two independently trained DHS64 models, whereas a third model was used to recalculate performance predictions for validation analysis (**Methods**). We successfully generated sequences predicted to be specific to almost every individual biosample (**Figure 1F-G**). For example, the validation model predicted sequences targeted to the retinoblastoma cell line WERI-Rb-1 to have strong on-target signal and weak off-target signal in all other cell types, including in related biosamples from the “neural” component (**Figure 1F**, top). However, cross-over within very closely related samples was unavoidable, for example within three fetal lung tissue biosamples that differed only in the originating subject (**Figure 1F**, bottom), as well as between the B-lymphocyte-derived GM12878 and GM12865 cell lines (**Figure 1G**). Sequence features of synthetic designs, such as GC content and sequence diversity, were similar to those from DHSs (**Supplementary Figure 2A-C**). No major performance differences were predicted between Fast SeqProp and DEN-generated sequences (**Supplementary Figure 2D**). Predicted performance for synthetic sequences was substantially higher than the most specific DHSs (**Supplementary Figure 2E**), even though our sequences were shorter than the median DHS length (196 nt)^50^.

### Sequences optimized for accessibility show cell type-specific enhancer activity

To test whether sequences designed for differential accessibility would drive cell type specific gene expression, we selected 10 cell lines from the 64 modeled biosamples in which to perform MPRAs. These cell lines span a representative range of tissue origins (components) covered by the model (**Figure 2A**). Beyond diversity of origin, the distinguishing feature of the 10 selected biosamples is that they are compatible with DNA library delivery in contrast to many others (e.g. fetal heart/kidney etc.). In terms of model prediction accuracy and data quality they are otherwise indistinguishable from the remaining 54 biosamples predicted by DHS64, suggesting that insights gained from experimental testing can generalize to all modeled samples.

**Figure 2.**
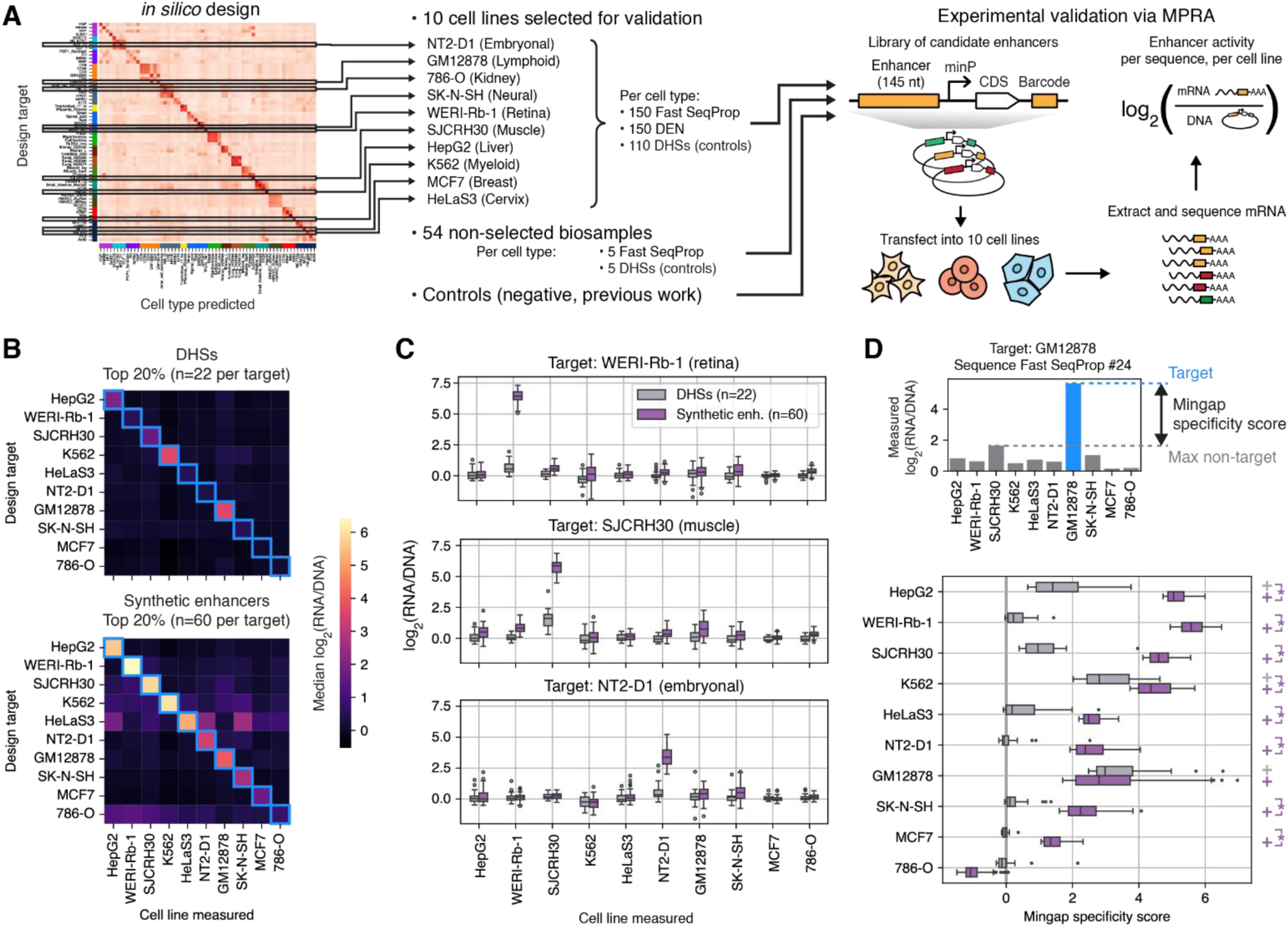
Synthetic sequences optimized for accessibility function as cell type-specific enhancers. **(A)** Experimental validation via MPRAs in a panel of 10 cell lines. **(B)** Measured enhancer activity of DHS-sourced (top) and NN-designed (bottom) sequences targeted to each assayed cell line. Rows represent the median of sequences targeted to each cell line. Light blue squares indicate the intended targets. **(C)** Distribution of measured enhancer activities for sequences targeting three representative cell lines. **(D)** Mingap specificity scores^8^ of DHS-sourced and NN-designed sequences. Top: illustration of mingap score calculation from enhancer activity measurements of an example sequence. Bottom: score distributions by target cell line and sequence source. Plus signs indicate significantly positive median scores (Wilcoxon test, one-sided, Bonferroni-corrected p-value < 0.05). Brackets with asterisks denote whether NN-generated sequences have significantly higher medians than DHS-derived sequences (Mann-Whitney U test, one sided, Bonferroni-corrected p-value < 0.05). In **(B)**, **(C)**, and **(D)**, only the top 20% of sequences by mingap score are used. See **Supplementary Figure 5** and **Supplementary Figure 6A** for distributions of all sequences and controls.

For each of these 10 biosamples we then chose 300 synthetic sequences designed as described above to be active in only the target cell line and inactive in the remaining 63 biosamples. As controls, we also tested 110 DHSs with the highest biosample-specific accessibility and enhancer-like chromatin annotations. DHSs were truncated to their central 145 nt to accommodate oligo pool synthesis, which had a small effect on predicted accessibility and specificity (**Supplementary Figure 3**). We cloned candidate enhancers into a reporter plasmid, transfected the resulting library into all cell lines, and extracted and sequenced mRNA, yielding good data quality and replicate correlation (**Supplementary Figure 4, Methods**). Enhancer activity was quantified in each cell line as log_2_(mRNA counts from cell line/DNA counts in plasmid library) (hereafter log_2_FC), and overall enhancer performance was assessed via the stringent mingap score^8^, defined as the difference between log_2_FC in the target cell line and the maximum log_2_FC across all non-targets (**Methods**).

Designed sequences exhibited the desired expression patterns across all but one cell line, and showed superior performance than corresponding DHSs, which were largely weak or inactive (**Figure 2B-D, Supplementary Figure 5**). Median measured mingap scores for synthetic sequences were significantly positive for 9/10 target cell lines (Wilcoxon test, one-sided, Bonferroni-corrected p-value < 0.05), and significantly higher than DHS-derived sequences for 8/10 (Mann-Whitney U test, one sided, Bonferroni-corrected p-value < 0.05, **Supplementary Figure 6A**). These differences were particularly pronounced among the top performing sequences (**Figure 2D**). In some cases, such as HepG2 and WERI-Rb-1, most synthetic sequences showed high specificity and strong on-target expression (**Supplementary Figure 5, Supplementary Figure 6A**), while in others, performance varied depending on the design method (**Supplementary Figure 5, Supplementary Figure 6A**). For the kidney-derived 786-O, all design methods failed to produce enhancers with positive specificity. DHS-derived sequences achieved significantly positive median scores only for GM12878 (**Supplementary Figure 6A**) and, among the top 20% performers, also for K562 and HepG2 (**Figure 2C**). Notably, GM12878 was the only target cell line where DHSs had equivalent performance to synthetic sequences. These conclusions remained consistent when using alternate specificity metrics, including one based on average non-target activities as well as the tissue specificity index^54^ (**Supplementary Figure 6B-C**). Finally, sequences designed for HepG2 and K562 showed similar specificity across these two cell lines compared to synthetic enhancers we previously designed via enhancer activity predictors trained on MPRA data^6^, and retained specificity across a larger variety of cell types (**Supplementary Figure 7**). These results demonstrate that synthetic enhancers optimized with DHS64 for accessibility can function as effective cell type-specific enhancers across diverse cell types, outperforming endogenous DHS-derived sequences, and with similar performance to those designed via predictors of enhancer activity.

Finally, to provide evidence of specificity at a larger scale, we assayed, across the same 10 cell lines, five DHS-sourced and five Fast SeqProp-designed sequences targeted to each of the 54 remaining biosamples. Since these sequences were not tested in their intended targets, DHS64 predicted them to show low activity, with some expression in cell lines corresponding to the same biological component as their target (**Supplementary Figure 8A**). Our results were consistent with these expectations (**Supplementary Figure 8B**). Specifically, synthetic enhancers had low differential (max - min log_2_FC < 2) activity across cell lines for 38/54 targets, whereas DHSs were overall inactive (**Supplementary Figure 8C-D**). Among the 16 targets with differential expression, enhancers designed for 11 of these showed activity in a cell line related to the target biosample, which was well-captured by DHS64 predictions (**Supplementary Figure 8E-H**). Only for a few target biosamples (e.g. the prostatic adenocarcinoma cell line PC3, the glioblastoma cell line A-172, and the variant mammary epithelial cell line vHMEC) did designed enhancers show unanticipated off-target activity (**Supplementary Figure 8F, I)**. Overall, sequences designed for accessibility specific to non-assayed cell types typically exhibited low enhancer activity, occasionally showed activity in a cell line related to the target cell type, and, only in rare cases, displayed unexpected broad expression.

### Comparing the sequence composition of DHSs and synthetic enhancers

We next investigated regulatory grammar features that explained the superior activity of synthetic enhancers compared to DHSs. We first scanned thousands of known TF binding sites, grouped into motif clusters by binding site similarity, across all selected DHSs and designed sequences (**Methods**). Thus, we identified 249 motif clusters corresponding to 730 TFs, present in sequences from at least one biosample. As observed in our previous work^6^, on average synthetic sequences contained more motifs per base pair than their DHS counterparts (**Figure 3A, Supplementary Figure 9A-B**). Furthermore, motifs present in DHSs accessible in fewer cell types or corresponding to TFs expressed with greater cell type or tissue specificity were generally used more in the synthetic sequences, whereas motifs that were ubiquitous in DHSs or corresponded to broadly expressed TFs were de-emphasized (**Supplementary Figure 9C-D, Supplementary Figure 10**).

**Figure 3.**
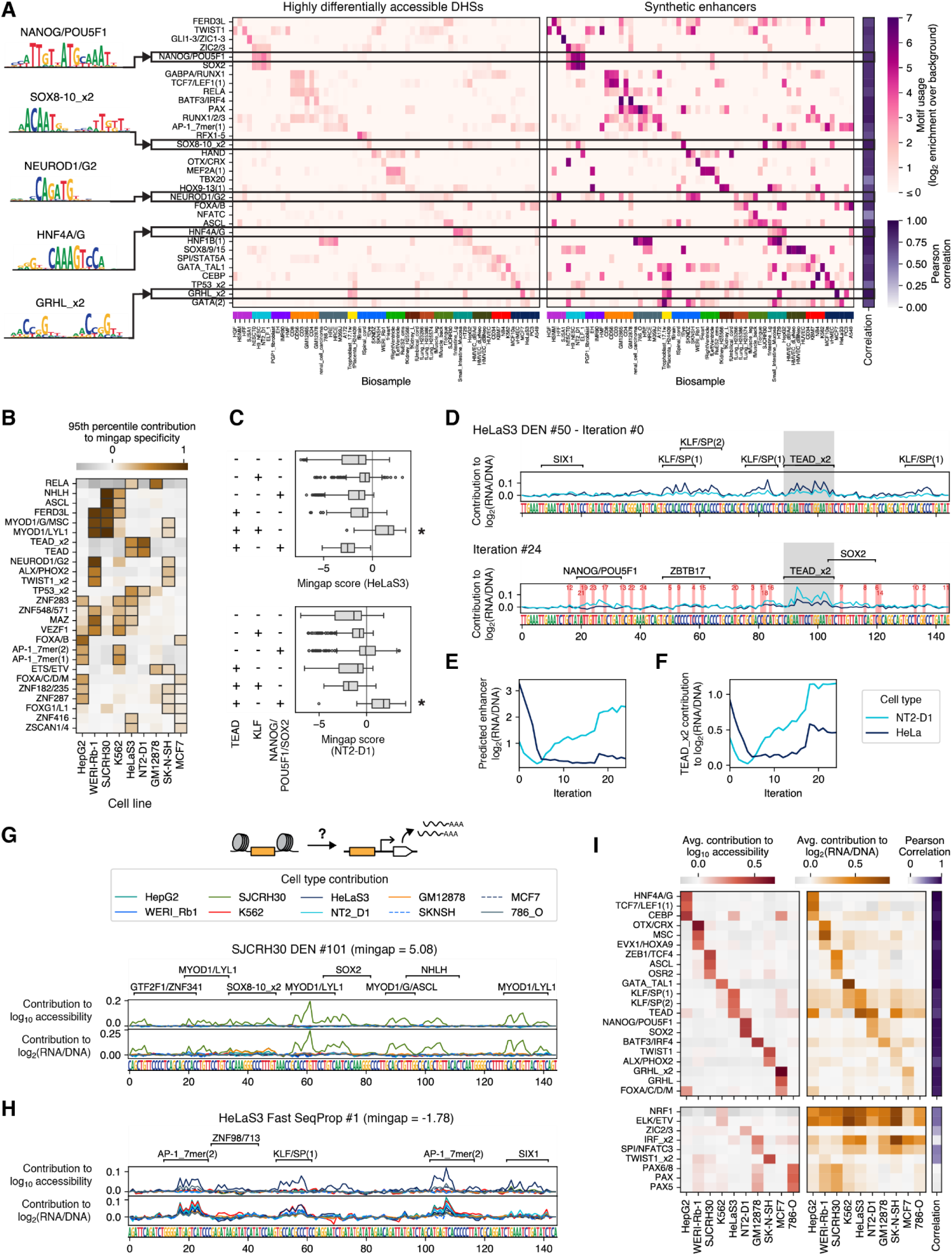
Transcription factor motif grammar of DHS accessibility is captured and amplified in synthetic enhancers. **(A)** Utilization of the most biosample-specific motifs in DHSs (top 1,000 per cell type by specificity, left) and synthetic enhancers (500 per cell type, right). For each target biosample, motif utilization was calculated as the log_2_ enrichment in motif counts per sequence relative to a background of DHSs specific to all non-target biosamples. Then, the motif with the highest enrichment on each biosample was selected and duplicates were removed. See also **Supplementary Figure 11. (B)** Motifs that contribute to specificity in multiple cell lines depending on sequence context. We used SHAP^55^ and DHS64-MPRA, a predictor of MPRA-measured enhancer activity (i.e. log_2_(RNA/DNA)), to calculate contributions to mingap specificity of every motif occurrence across every assayed sequence towards every cell line (**Methods**). Then, for each motif and cell line, the 95^th^ percentile of contributions was selected and plotted. This indicates a motif’s potential to contribute to cell line-specific activity under the appropriate context. Finally, motifs were retained if they showed substantial contribution to more than one cell line. See also **Supplementary Figure 14. (C)** Mingap score towards HeLaS3 (top) and NT2-D1 (bottom) of assayed sequences targeting individual cell types containing motif combinations that include TEAD. Asterisks indicate median activity above zero (Wilcoxon one-sided, Bonferroni-corrected p < 5e-2). TEAD can drive specificity towards either cell line depending on co-occurring motifs, whereas none of these motifs can drive specificity on their own. **(D-F)** *In-silico* evolution of an enhancer from HeLaS3 to NT2-D1 specificity. Starting from an MPRA-validated HeLaS3-specific synthetic enhancer, a greedy *in silico* mutagenesis approach was used to iteratively select mutations that increase the difference in NT2-D1 minus HeLaS3 contributions of a TEAD motif. Mutations inside TEAD were not allowed. Throughout this process, TEAD contributions change from HeLaS3-to NT2-D1-specific without changes in its own sequence. **(D)** Initial (top) and final (bottom) sequences, motifs identified via FIMO, and enhancer activity contributions. The TEAD motif region is shaded in grey. Mutations introduced are shaded in red. **(E)** Evolution of predicted enhancer activity. **(F)** Evolution of TEAD motif contributions. **(G and H)** Concordance between accessibility and enhancer regulatory features of two example sequences. From top to bottom: motifs, contributions to predicted accessibility in all 10 cell lines, contributions to predicted enhancer activity, and sequence. Contributions were calculated using SHAP with DHS64 for accessibility and DHS64-MPRA for enhancer activity (**Methods**). For the well-performing sequence in **(G)**, SJCRH30-specific contributions to accessibility and activity match closely. For the poorly performing sequence in **(H)**, motifs contribute specifically to HeLaS3 accessibility, but broadly to enhancer activity. **(I)** Motifs expected to be the most important drivers of cell type specificity, and their average contributions towards cell type accessibility (left) and enhancer activity (right). For each cell line, we selected the top 3 motifs with the highest contribution to a mingap score calculated on accessibility contributions (target minus max non-target contributions). We then calculated Pearson correlations between cell line contributions to accessibility and enhancer activity, as a measure of how well the regulatory grammar of accessibility transferred to enhancer activity. Finally, we separated motifs based on whether they correlated well (R > 0.6, top) or poorly (R < 0.6, bottom). See also **Supplementary Figure 17**.

Among the motifs emphasized, usage patterns across cell types correlated between DHSs and synthetic sequences (**Figure 3A, Supplementary Figure 11**), indicating that synthetic sequences recapitulated and/or exaggerated the motif grammar present in the most specific DHSs. Of the motifs most enriched in a specific biosample, many were not uniquely associated with that biosample or even the corresponding biological component. For instance, the SOX8-10 motif, which was most enriched in the fetal eye sample, was also present in sequences from the muscle-derived cell line SJCRH30 and liver-derived cell line HepG2 among others. Overall, these findings point to a complex and rich TFBS grammar within DHSs – captured and amplified by our model-based design approach – where specificity is not simply conferred by individual motifs.

### A combinatorial TF motif code explains cell type specificity

Variation in MPRA-measured enhancer performance suggests that not all TF motifs, present due to their predicted contribution to accessibility, were successful at driving enhancer activity. To identify specific motif grammar features that enhanced or hindered specificity, we first compared motif usage between the top and bottom 25% of all tested single-target synthetic enhancers ranked by mingap score (**Supplementary Figure 12, Methods**). For many target cell types – most prominently WERI-Rb1 (retina), SJCRH30 (muscle), and HeLaS3 (cervix) – certain motifs were differentially enriched between the top and bottom performing sequences, suggesting an explanation for the observed performance gap. A particularly dramatic example was HeLaS3, where KLF motifs were associated with high-performing and AP-1 motifs with low-performing sequences. For other cell lines, however, the most used motifs were not statistically enriched in either set, suggesting that the mere presence of individual motifs could not sufficiently explain performance differences.

We next hypothesized that motif effects could change depending on sequence context and investigated this hypothesis using explainable AI. We developed DHS64-MPRA, a NN predictor of MPRA-measured enhancer activity, by finetuning DHS64 on our MPRA data (**Supplementary Figure 13, Methods**). We used SHAP^55^ to compute DHS64-MPRA-derived nucleotide-level contributions towards enhancer activity on every tested sequence and cell line. We then calculated contributions of every motif occurrence in their full sequence context by adding up contributions of all nucleotides within the motif matching region (**Methods**). Some motifs contributed to activity in only one cell line across occurrences, such as the liver associated HNF4A/G on the liver cell line HepG2 (**Supplementary Figure 14A**). However, many motifs switched cell type-specific contributions depending on context, such as the muscle-associated MYOD1/G (**Supplementary Figure 14B-C**).

We next calculated motif contributions to mingap specificity scores, and, for each cell line, used the 95h percentile across motif occurrences as a measure of a motif’s ability to drive cell type-specific activity in a favorable context. We thus found 40 motifs that could meaningfully drive cell type-specific enhancer activity in multiple cell lines (**Figure 3B**, **Supplementary Figure 14D**). In some cases, enhancer specificity was achieved via motif combinations that included these multi-specific motifs (**Figure 3C**, **Supplementary Figure 15**). For example, the TEAD motif drove cell type-specific activity in HeLaS3 cells when combined with KLF/SP but drove activity in NT2-D1 cells when paired with NANOG/POU5F1/SOX2. The same motifs without their companions were not sufficient to drive expression (**Figure 3C**). Similarly, FOXA, in combination with HNF4A/G drove activity specifically in HepG2 and with GRHL drove specificity to MCF7. SOX8-10 combined with HNF4A/G or MYOD1/G drove activity in HepG2 or SJCRH30, but none of these motifs were associated with strong performance without their partners (**Supplementary Figure 15**).

To test causality, we assayed sequences with select motifs embedded into inert background sequences (**Supplementary Figure 16A**). These experiments recapitulated several of the observed effects, such as insufficiency of GRHL2, MYOD1, and HNF4A to drive MCF7, SJCRH30, and HepG2 activity (**Supplementary Figure 16G-I**), and the ability of FOXD3 to rescue HNF4A activity in HepG2 (**Supplementary Figure 16J-M**).

We further illustrate how a motif can contribute to specificity in multiple cell types via an *in silico* evolution experiment. We first identified an MPRA-characterized HeLaS3-specific sequence containing a TEAD motif with high predicted HeLaS3 contribution. Keeping the TEAD motif constant, we iteratively mutated the remaining sequence to increase the contribution of the TEAD site towards NT2-D1 while decreasing HeLaS3 contributions until convergence. (**Figure 3D-F**, **Methods**). The initial ∼7 mutations disrupted four distinct KLF/SP motifs and largely destroyed predicted HeLaS3 enhancer activity, whereas the following 17 mutations created NANOG/POU5F1 and SOX2 motifs and progressively increased enhancer specificity towards NT2-D1. In this process, TEAD motif contributions switched from HeLaS3-to NT2-D1-specific despite there being no change in its sequence. Overall, these results highlight the varied and complex modes of cell type-specific combinatorial motif effects captured and exploited by our enhancer design method.

### Explainable AI helps distinguish between TFs that contribute to accessibility and enhancer activity

Finally, we used explainable AI to investigate what aspects of the regulatory grammar learned from genomic accessibility translated into enhancer function. To this end, we compared nucleotide contributions to enhancer activity calculated via DHS64-MPRA with contributions to accessibility predicted by DHS64 in all assayed cell lines (**Figure 3G-I, Methods**). In high-performing sequences, we observed strong concordance between contributions from the accessibility and MPRA models (**Figure 3G**), indicating that the regulatory grammar learned from DHSs translated effectively into enhancer function. In contrast, poorly performing sequences showed noticeable divergence between the two models, highlighting regulatory features that failed to drive expression or drove expression too broadly (**Figure 3H**).

We then calculated the average contribution of every motif towards accessibility and enhancer activity in each cell type (**Figure 3F**, **Supplementary Figure 17A-B, Methods**), and measured agreement between both via Pearson correlations. While many motifs featured strong agreement (e.g. MYOD1/G for SJCRH30 (muscle), HNF4A/G for HepG2 (liver), and NANOG/POU5F1 for NT2-D1 (embryonal), other motifs – such as those from the IRF and AP-1 families – exhibited broad contributions to enhancer activity even when their accessibility contributions were more cell type-specific. Conversely, motifs from the PAX family contributed strongly to accessibility in 786-O, but had negligible impact on enhancer activity, helping explain the lack of functional enhancers in that cell line (**Figure 3E** bottom, **Supplementary Figure 17B**). Despite these discrepancies, accessibility contributions of motifs were generally predictive of their cell type-specific effects on enhancer activity (**Supplementary Figure 17C**). Overall, our analysis indicates that our design approach effectively captures and amplifies much of the regulatory grammar underlying cell type-specific accessibility into functional enhancer activity, though certain motifs remain challenging to translate effectively.

### Accessibility predictors enable programming complex functions into synthetic enhancers

We next explored whether our approach could program enhancers with complex functions, including responding to multiple cell types and predictably tuning output activity. First, we designed enhancers with specificity towards two (**Figure 4A-C**) and three (**Figure 4D-F**) unrelated cell types and validated them via MPRAs as above. We randomly selected eight pairs and eight triplets from the 10 assayed cell lines, ensuring that each target group contained cell types from different biological components. Using Fast SeqProp, we generated 117 sequences optimized for high accessibility in all target cell types within each group (**Supplementary Figure 18A** and E, bottom, **Methods**). We additionally sought to include, for each target group, 50 DHSs with enhancer-like chromatin marks and positive peak calls in all target cell types (**Supplementary Figure 18A and E**, top, **Methods**). However, such DHSs were scarce, and insufficient numbers were found for 1/8 pairs and 5/8 triplets (**Supplementary Figure 18B and F**). Notably, no DHSs met our criteria for one triplet. We scored these designs via an extended mingap score obtained by subtracting the minimum log_2_FC across targets from the maximum nontarget log_2_FC (**Figure 4C**, top). When considering the top 20% performing sequences from each target group, synthetic enhancers frequently exhibited the expected expression patterns whereas DHSs were almost always inactive (**Figure 4A-F, Supplementary Figure 18C and G**). Synthetic sequences had significantly positive median mingap scores in 5/8 target pairs and 4/8 target triplets (Wilcoxon test, one-sided, Bonferroni-corrected p-value < 0.05 **Figure 4B** and **E**). Enhancers generally achieved specificity by combining regulatory elements from enhancers designed for individual targets (**Figure 4G-I**, **Supplementary Figure 19**). For example, sequences targeting the liver-associated HepG2 and retina-associated WERI-Rb-1 cell lines used HNF4A/G and FOXA/C/D/M motifs, specific to HepG2, along with NEUROD1/G2 and OTX/CRX, specific to WERI-Rb-1. However, the total number of TF motifs per enhancer remained similar (**Supplementary Figure 19**), with each motif generally present at a lower copy number compared to their single-target counterparts (**Figure 4H-I**). This may explain the observed decrease in target predicted accessibility and measured log_2_FC compared to single-target designs: as the number of targets increases, sequence length becomes insufficient to accommodate all regulatory elements required for strong activation in all target cell types. Supporting this hypothesis, longer sequences designed for multiple targets achieved higher predicted activities (**Supplementary Figure 20**). Thus, longer enhancers may be necessary to maintain specificity as the number of targets increase.

**Figure 4.**
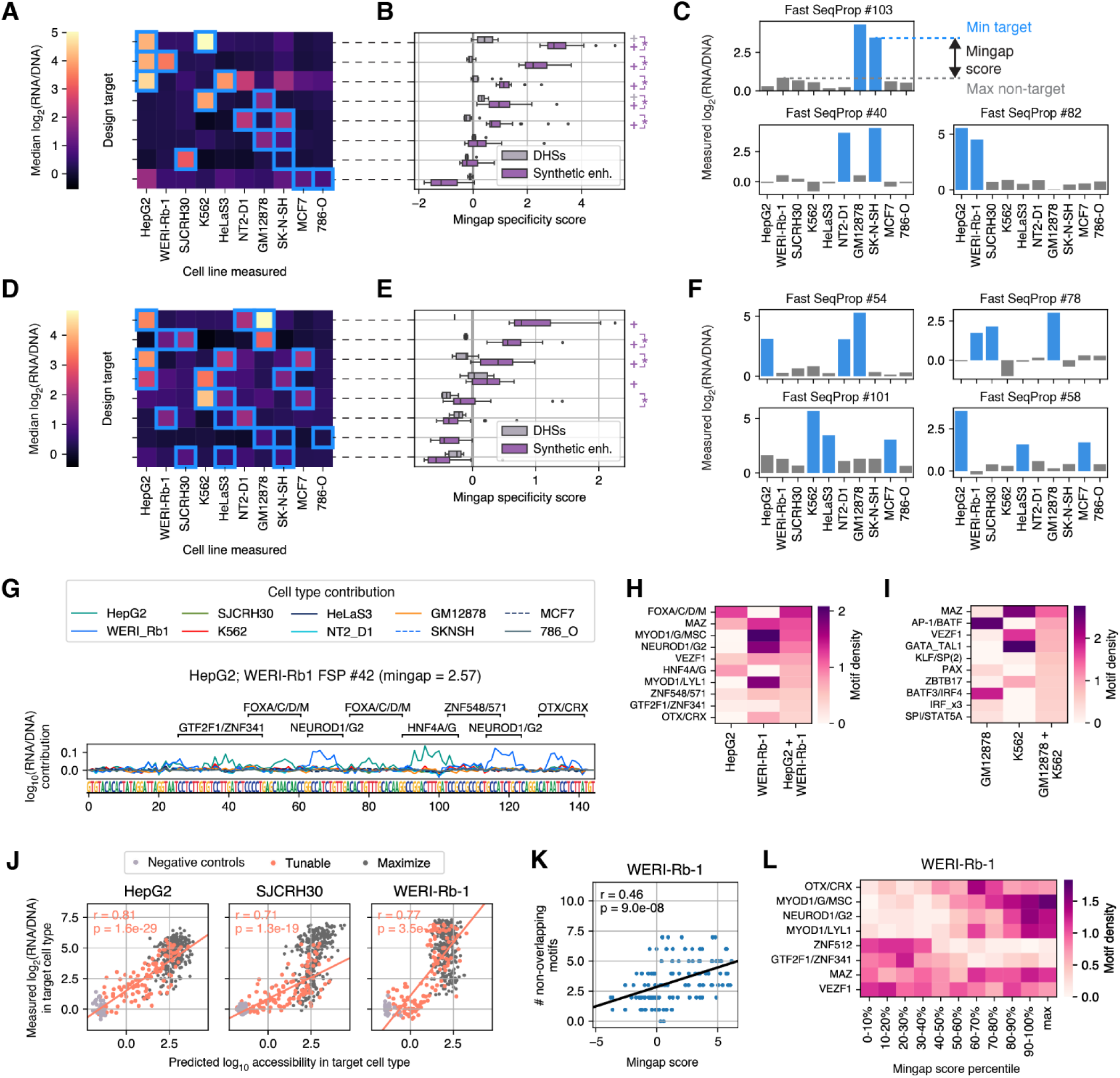
Synthetic enhancers achieve complex design objectives such as responding to multiple cell types and predictably tuning on-target activity. (A-C) Median expression **(A)**, mingap specificity **(B)**, and activity of top-performing synthetic enhancers **(C)** targeting the indicated cell line pairs. In **(A)** and **(B)**, only the top 20% by mingap score are considered. In **(B)**, “+” denotes significantly positive mingap scores, and brackets with “*” denote significantly greater performance in synthetic compared to DHS-sourced enhancers, as in Figure 2. In **(C)**, blue bars indicate target cell types. **(D-F)** Median expression **(D)**, mingap specificity **(E)**, and activity of top-performing synthetic enhancers **(F)** targeting the indicated cell line triplets. Median expression and top-performing individual examples from control DHSs can be found in **Supplementary Figure 18. (G)** FIMO-aligned TF motifs and nucleotide contributions to enhancer activity predictions, for an example enhancer targeting cell lines HepG2 and WERI-Rb-1. **(H-I)** TF motif density (avg. # motifs per sequence) in two double-target enhancer sets, compared to their corresponding single-target designs. Only enhancers within the top 20% by mingap score in each set were used. For each target pair, the top 10 motifs by density are shown. See also **Supplementary Figure 19**. **(J)** Relationship between predicted target accessibility and measured target enhancer activity for sequences designed for maximal (i.e. Figure 2) and tunable target activity, alongside negative controls, for three target cell lines. Linear regression fits, Pearson r coefficients, and p-values were calculated using tunable enhancers only (salmon markers). See **Supplementary Figure 21** for data from all other cell lines. **(K)** Total number of motifs in tunable WERI-Rb-1 enhancers as a function of their experimentally measured mingap specificity. **(L)** Motif density of individual TF motifs for tunable enhancers binned by mingap specificity. Enhancers designed for maximal specificity (Figure 2) are shown next for comparison with the label “max”.

Next, we sought to design cell type-specific enhancers with tunable intermediate target activities (**Figure 4J-L**). Using Fast SeqProp, we designed 120 sequences per cell line, with target accessibility setpoints ranging from inactive to maximally active, and validated them experimentally via MPRAs (**Methods**). For 7/10 cell lines, target enhancer activity was tunable, specific, and significantly positively correlated with predicted target accessibility across the entire range (**Figure 4J, Supplementary Figure 21**). For SK-N-SH, while specificity was tunable, target activity increased but later decreased with increasing setpoints. Finally, for HeLaS3 and 786-O, we observed tunable on-target activity but lack of specificity. These aligned with previous failures in the context of designing maximally specific enhancers (**Supplementary Figure 6A**. Tunable activity was generally achieved by increasing the number of cell type-specific TF motifs along with increasing setpoints (**Supplementary Figure 22**). However, in some cases distinct TFs were associated with different activity levels. For example, WERI-Rb-1 enhancers switched from ZNF512 and GTF2F1/ZNF341 to NEUROD1/G2 and OTX/CRX as target activity increased (**Figure 4K-L**). These results show that optimizing for submaximal predicted accessibility is a feasible strategy to design cell type-specific enhancers with tunable activity. In summary, DHS64 enabled the predictable design of enhancers with complex functions, including tunable target activities and specificity for multiple cell types.

### Enhancers designed for retinal targets are active in mouse retinas

To evaluate whether our enhancers function in a multicellular biological system, we performed a retinal tissue MPRA by transfecting our enhancer library in P0 mouse retina (**Figure 5**, **Methods**). Although we did not design enhancers specifically for mouse retina, two of our modeled biosamples – the retinoblastoma cell line WERI-Rb-1 and fetal eye tissue – originate from related sources, leading us to hypothesize that sequences targeting these biosamples would be functional in the retina. Consistent with this expectation, WERI-Rb-1 and fetal eye-targeted synthetic enhancers displayed the highest activity, outperforming synthetic enhancers for all other targets (median log_2_FC = 4.88 and 3.93, respectively, **Figure 5B**), as well as to DHS-sourced controls which were largely inactive (**Supplementary Figure 23A**). Enhancer activity in mouse retina and in WERI-Rb-1 were highly correlated (r = 0.74, **Supplementary Figure 23B**): high-performing WERI-Rb-1 enhancers generally showed high retinal activity (**Figure 5D**, top) while enhancers designed for tunable submaximal WERI-Rb-1 activity displayed graded log_2_FC levels (**Figure 5C**). On the other hand, while all 5 enhancers targeted to the fetal eye biosample drove gene expression in the retina, two of them showed low activity in all our MPRA-assayed cell lines, including WERI-Rb-1 (**Figure 5D**, bottom, **Supplementary Figure 23C**). TF motif alignment and model interpretation suggested that WERI-Rb-1 and fetal eye-targeted sequences leverage distinct regulatory grammar (**Figure 5E**): whereas WERI-Rb-1 designs use NEUROD1/G2, ATOH1/MSC, and a motif from the retinal cone TF CRX, fetal eye-targeted sequences use PAX6, associated with eye development and gene expression in the retinal ganglion and other eye structures^56^, as well as SOX9, associated with expression in the retinal pigment epithelium^57^, which our models predict will not contribute to WERI-Rb-1 activity. In conclusion, our accessibility predictor-based approach enables the design of functional, cell type-specific enhancers that outperform their genomically derived counterparts, even in tissues.

**Figure 5.**
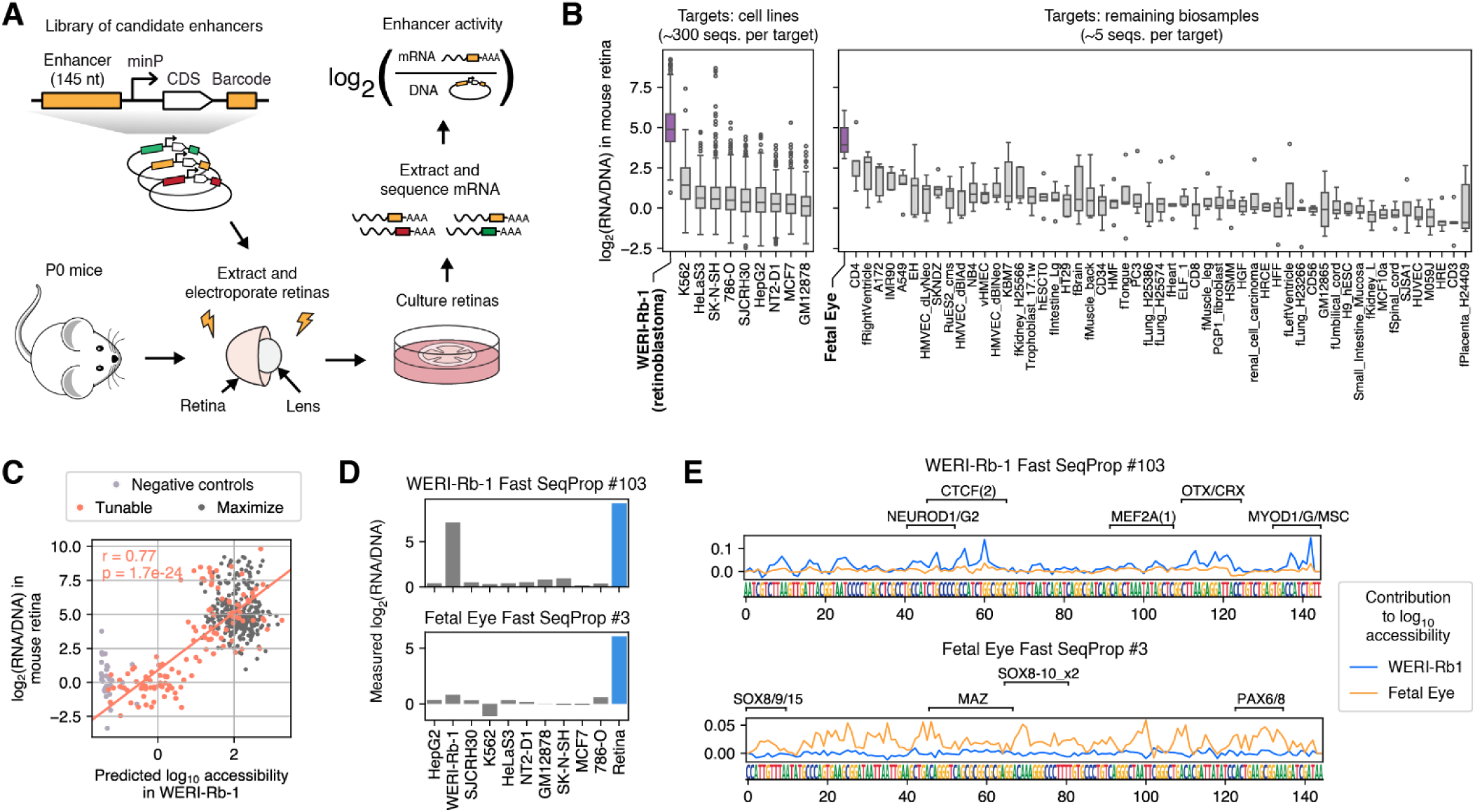
Synthetic enhancers designed for eye-related biosamples function in mouse retinas. **(A)** Schematic of the mouse retina MPRA. **(B)** Distributions of measured log_2_(RNA/DNA) from synthetic enhancers targeted to all DHS64 modeled biosamples. Targets are divided in two groups: main target cell lines (Figure 2), for which we designed and assayed ∼300 sequences, and all 54 remaining biosamples (**Supplementary Figure 8**), for which we included ∼5 sequences per target. In each panel, targets were sorted by median log_2_(RNA/DNA). Eye-related biosamples have been highlighted. See **Supplementary Figure 23** for matched DHS-sourced controls. **(C)** Tunable WERI-Rb-1 enhancers: comparison between predicted WERI-Rb-1 accessibility and measured mouse retina log_2_(RNA/DNA). Each marker represents an individual sequence. **(D)** log_2_(RNA/DNA), measured across all cell lines and in the mouse retina, of two example sequences: one originally designed for WERI-Rb-1, and one designed for the fetal eye biosample. **(E)** Sequences, TF motifs, and contributions towards predicted accessibility of the example enhancers in **(D)**.

### An atlas of synthetic human enhancers generated by models trained on the full DNase I Index

To enable targeting the full range of sample types in the DNase I Index, we trained an extended model, DHS733, to predict chromatin accessibility across all 733 biosamples (**Figure 6A**, **Supplementary Figure 24**). In contrast to DHS64, where only the most cell type-specific DHSs were used for training, DHS733 was trained on all DHSs by explicitly maximizing the prediction-measurement correlation across biosamples for each DHS^30,36^. This formulation enabled us to utilize the complete dataset while still capturing cell type-specific regulation as effectively as with our previous filtering-based approach (**Supplementary Figure 24, Methods**). Next, we constructed an “Atlas of Synthetic Human Enhancers,” comprising enhancers targeted to every one of the 261 non-redundant biosamples (i.e. with unique identifying names). We used Fast SeqProp to optimize 200 sequences with predicted accessibility specific to each target with respect to all other biosamples, for a total of 52.2k designs (**Figure 6B**, **Supplementary Figure 25A-C, Supplementary Table 12**, **Methods**). To validate their function, we performed an *in silico* version of the MPRA validation workflow in **Figure 2**: starting from enhancers from the Atlas targeted to each of our 10 previously assayed cell lines, we estimated cell line enhancer activities via DHS64-MPRA, and calculated mingap scores as before. We found that enhancers from the DHS733-generated Atlas performed comparably to those designed with DHS64 (**Supplementary Figure 25C-E**), demonstrating that our expanded model can successfully support cell type-specific enhancer design. Although validation is limited to sample types for which we have MPRA data and could therefore train DHS64-MPRA, the high success rate in those cases suggests that designs targeting other samples also have a high likelihood of being functional enhancers. Moreover, the agreement between DHS733 and DHS64 designs implies that the sequence grammar used to distinguish a particular sample from 63 broadly selected samples are the same that distinguish it from all other samples in the DNase Index.

**Figure 6.**
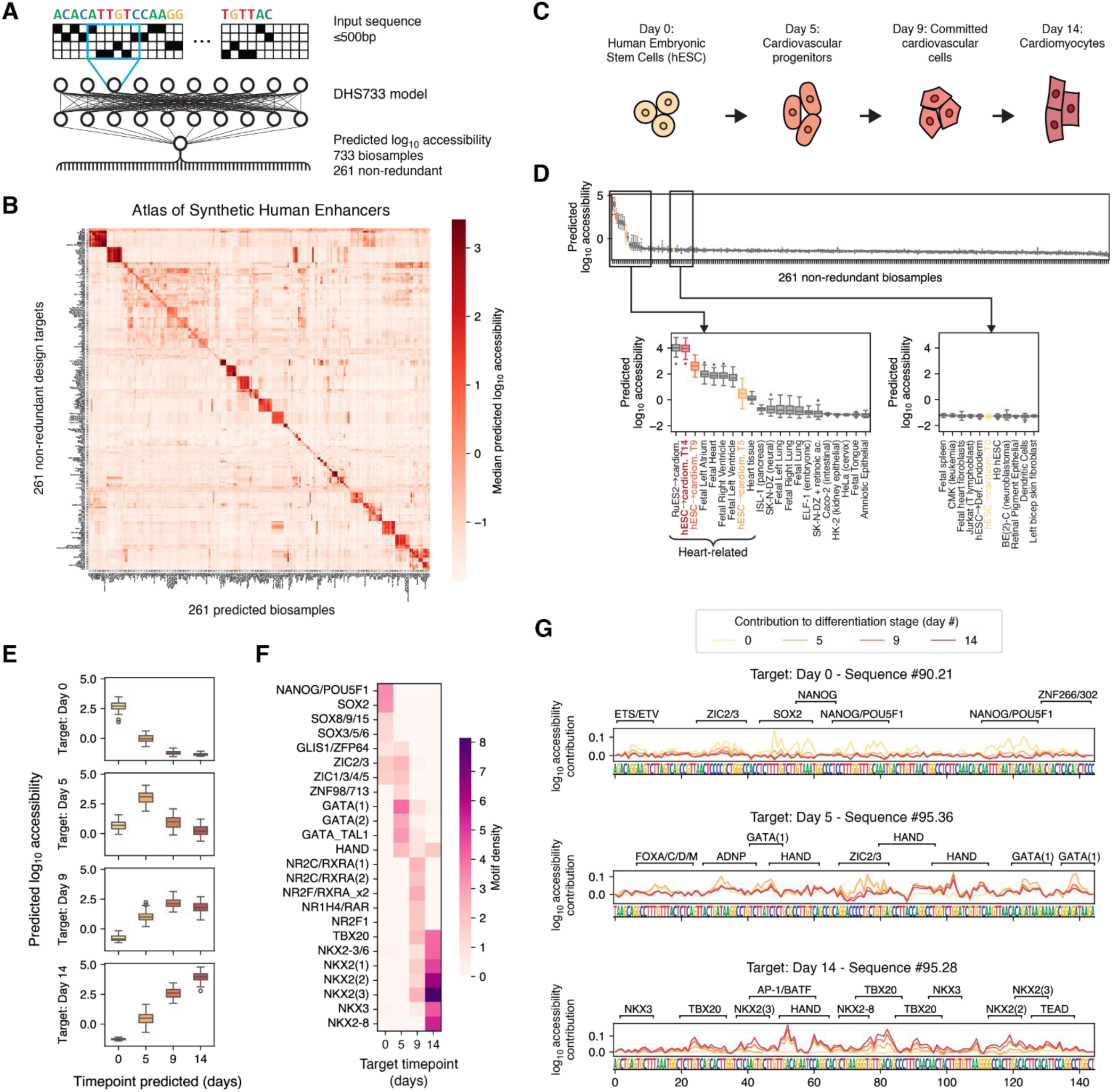
An Atlas of Synthetic Human Enhancers targeted to every biosample in the DNase-I Index includes differentiation stage-specific sequences. **(A)** DHS733, trained on the entire DNase I Index, predicts chromatin accessibility across all 733 biosamples, 261 of which are non-redundant. **(B)** The Atlas of Synthetic Human Enhancers: 200 enhancers designed for each of the 261 non-redundant biosamples predicted by the DHS733 model. The plot shows median accessibility for each target over each biosample, predicted by an independent model not used during sequence design (**Methods**). **(C)** The DNase I Index includes a time-course experiment measuring accessibility during differentiation of the H7 human embryonic stem cell (hESC) line into cardiomyocytes, sampled at four timepoints. **(D)** Predicted accessibility of synthetic enhancers in the Atlas targeted to the final differentiation timepoint (day 14). Biosamples from the time-course experiment are colored as in **(C)**. Top: predictions across all 261 non-redundant biosamples, sorted by median. Bottom left: 21 biosamples with the highest predictions. Bottom right: initial state (hESC, day 0) and unrelated biosamples with similar predicted accessibility. **(E)** Predicted accessibility across differentiation time of enhancers designed to target each timepoint. Data in the “day 14” panel is the same as in **(D)**. **(F)** Top 7 motifs with the highest density (# motifs / sequence) in enhancers designed towards each timepoint. Duplicates appearing in multiple targets were removed randomly. **(G)** Example designs targeting the 0-, 5-, and 14-day timepoints, along with TF motifs identified via FIMO and predicted contributions (**Methods**).

### The Atlas of Synthetic Human Enhancers includes differentiation state-specific sequences

Beyond static cell types, the epigenetic and gene expression state of a cell can change dynamically in response to external stimuli or during processes such as development and differentiation. The DNase I Index includes biosamples that capture such dynamic states, including cells stimulated with hormones and morphogens, as well as stem cells undergoing directed differentiation into endodermal, neural, hematopoietic, and cardiac cells. To evaluate whether enhancers in the Atlas could discriminate between closely related cell states, we focused on biosamples from a time course experiment where human embryonic stem cells (hESCs) were induced to differentiate into cardiomyocytes, with chromatin accessibility profiled at 0, 5, 9, and 14 days of differentiation (**Figure 6C**). Synthetic enhancers targeted to these biosamples showed high predicted on-target and low off-target activity in all but a few biologically related samples. For example, sequences targeted to the most differentiated cardiomyocyte state (day 14) also showed activity in other heart-related samples, but very low predicted activity in undifferentiated hESCs (**Figure 6D**). Sequences designed for every timepoint showed the desired predicted accessibility profile throughout the differentiation time course (**Figure 6E**): sequences targeted to the initial pluripotent state had high predicted accessibility at day 0, which sharply declined by day 5 and was nearly absent by day 9; day 5-targeted sequences showed a peak of predicted activity at day 5 and lower activity at surrounding timepoints; and sequences targeted to the final differentiated states exhibited low accessibility early in differentiation that increased over time. Analysis of the designed sequences revealed features consistent with known regulatory grammar (**Figure 6F, G**): sequences targeting the initial pluripotent state contained motifs for pluripotency-associated TFs such as SOX2, NANOG, and POU5F1^58^; day 5-targeted sequences contained motifs for TFs expressed during the early postcardiac mesoderm stage such as those from the HAND (e.g. HAND2) and GATA (e.g. GATA4) families; and day 14-targeted sequences were enriched for cardiac development motifs such as those from the TBX (e.g. TBX5, 20) and NKX2 (e.g. NKX2-5) families^59,60^. Together, these results suggest that enhancers in the Atlas can be responsive not only fully differentiated cell types but also to transient cell states during differentiation.

## Discussion

Here, we showed that models trained on an accessibility atlas of hundreds of human biosamples enable designing enhancers with high specificity towards almost any sample type, against tens and even hundreds of others (**Figure 1**, **Figure 2**, **Figure 6**). Furthermore, we provide two repositories of AI-designed enhancers, one with 32k sequences targeting each of the DHS64-modeled biosamples (**Supplementary Table 5**), and a larger Atlas of 52.2k sequences targeting every non-redundant biosample in the DNase I Index (**Supplementary Table 12**), the largest of such repositories to date. Where such comparisons are available, the performance of these enhancers is similar to those designed with models directly trained on enhancer activity data (**Supplementary Figure 7**). By measuring enhancer activities of thousands of candidate sequences in multiple cell lines and comparing their sequence features with expectations from models trained on accessibility data, our work fills a long-standing gap in understanding the relationship between drivers of accessibility and activity. Importantly, the motif grammar identified by DHS64 was sufficient to provide synthetic enhancers with specificity towards 9 out of 10 tested cell lines (**Figure 2B-D**), suggesting extensive overlap between cis-regulatory elements responsible for accessibility and activity. We also found that most individual TF motifs contribute similarly to accessibility and activity across cell types (**Figure 3I, Supplementary Figure 17**).

However, we also observed a non-negligible number of motifs whose accessibility contributions did not transfer properly to enhancer activity (**Figure 3F**, **Supplementary Figure 17**). These “failures” took many different forms, such as PAX motifs whose predicted effect in 786-O failed to materialize, TP53 which showed broad enhancer effects despite limited predicted accessibility influence, and AP-1 which contributed to GM12878 specificity but showed significant off-target activity in HeLaS3. Since AP-1 has been observed to cooperate with other motifs to drive specificity in many cellular processes^61–63^, our results raise the question of whether our models or training data were insufficient to capture AP-1 grammar nuances, or whether the observed effects are a true mismatch between accessibility and enhancer grammar. A future study where accessibility and enhancer activity are simultaneously measured for the same reporter would be well suited to answering these questions.

Our enhancers can be programmed with complex responses including tunable target expression and simultaneous activity in multiple cell types, even when corresponding natural enhancers are nonexistent (**Figure 4**). In enhancers with tunable target activities, increases in TF density were almost always monotonically correlated with target activity (**Figure 4K-L, Supplementary Figure 22**), whereas enhancers designed for multiple cell types combined motif grammar from each target (**Figure 4A-I, Supplementary Figure 19**). Motifs were generally evenly distributed across the sequence length, with no obvious clustering corresponding to individual cell types (**Figure 4G**). This flexible motif positioning more closely resembles the “billboard” enhancer model, as opposed to the “enhanceosome” model where motifs are located within rigid position and distance constrains^15^.

When tested in mice retina, synthetic enhancers targeted to eye-related biosamples showed the highest activity among all tested sequences (**Figure 5**), showing that they retain their specificity in tissues. These experiments take advantage of the observation that designs targeting very closely related biosamples types are predicted to show crosscutting activity. Future work using single cell RNA sequencing will be necessary to determine which specific cell types within the retina are targeted by each of the synthetic enhancers, given that the enhancers targeting WERI-Rb-1 and fetal eye tissue, respectively, achieved high activity in mouse retina using different motif grammar and thus potentially targeting different cell types.

Model-guided design of biological sequences requires deciding, explicitly or otherwise, to what extent to deviate from natural examples where model accuracy is the highest but which may have submaximal activity. Consistent with previous results^6,8^, we found that high-performing synthetic enhancers do not just borrow but also exaggerate sequence features from their natural counterparts, including TF motifs that mediate specificity towards one or multiple cell types depending on context (**Figure 3A-F**). These results suggest that, if the goal is to design enhancers with maximal performance, an excessive focus on retaining natural features via performance caps^7,10^ or generative models that maintain motif grammar features^64,65^ may be ultimately detrimental.

In our experiments, genome-sourced enhancers were weaker and less specific than synthetic sequences (**Figure 2**), raising the question of why nature would leave such unrealized potential for regulatory activity. A confounding factor is the assayed enhancer length: DHSs have a median length of 196 nt^50^, and enhancers in the literature are sometimes defined to include up to 2,000 nt^15,66,67^, but here we truncate DHSs to their central 145 nt. Bases removed during truncation may be important for activity, and indeed DHS64 predicts a corresponding decrease in accessibility (**Supplementary Figure 3**). However, this effect is small compared to the difference with synthetic enhancers (**Supplementary Figure 2E**). Furthermore, even when compared with full-length DHSs, our enhancers contain a higher motif density per base pair and an enrichment for the most cell type-specific motifs (**Supplementary Figure 9**), suggesting that they would maintain their strength advantage against full DHSs. While individual genomic enhancers might be weaker, multiple enhancers often act together on the same promoter^67^, with the number of TF motifs necessary for activation likely split among them. Why evolution would favor this “distributed” regulatory model is an open question^67^, but our results suggest that an alternative mode based on stronger compact enhancers is at least possible. In addition, enhancers are subject to rapid evolution^15,68^, with new enhancers arising via a small number of mutations on neutral DNA^69^ or by repurposing existing enhancers to alternative cell states^70^. Thus, maintaining enhancer plasticity – likely a key facilitator of mammalian evolution^15,70^ – may be at odds with regulatory elements consisting of densely packed TF motifs. Still, we also observe that the specificity of synthetic enhancers can be switched from one cell type to another with a limited number of mutations and while maintaining a subset of the motif content (**Figure 3D-F**). These results suggest that even enhancers with high information density can maintain a high degree of evolvability. Finally, strong activation may not be desirable in all situations. For example, gain-of-function mutations that increase enhancer-TF affinity can result in defects during limb^71^ and heart^72^ development.

Finally, while synthetic enhancers targeting fully differentiated cell types are now within reach, targeting general cell states such as disease and transient differentiation states could enable a wider array of biotechnological applications. Our Atlas of Synthetic Enhancers contains enhancers targeting stem cells differentiating into various lineages, including cardiomyocytes at various intermediate stages (**Figure 6**). An exciting future application is using these to express differentiation factors with temporal precision to guide differentiation of stem cells^5^, thereby improving differentiation efficiency and possibly even reaching currently unattainable cell states. Similarly, specifically targeting diseased states could enable gene therapies with increased specificity. In conclusion, our work shows that functional enhancers specific to a variety of human cell types can be designed from neural network models of DNA accessibility, and provides a blueprint for generalizing such an approach to other systems with available accessibility data, including to transient cell states in development, differentiation, and disease.

## Funding

This work was supported by NIH awards R33CA286947, R01GM149631 and R56HG013312 to G.S., NIH award R35HG011317 to W.M., NIH awards R01EY028584 and R01EY033364 to T.J.C., and a BrightFocus Postdoctoral Fellowship in Macular Degeneration to L.V.B.

## Author contributions

S.C-H conceptualized the study, performed neural network modeling, enhancer design, experimental design, cell culture experiments, data analysis, and wrote the manuscript. C.Y. conceptualized the study, performed modeling, cell culture experiments, data analysis, and wrote the manuscript. L.V. performed mouse experiments. T.J.C designed mouse experiments and provided feedback for the manuscript. W.M. conceptualized the study, assisted with analysis of DNase I Index data, designed experiments, and provided feedback for the manuscript. G.S. conceptualized the study, designed experiments, and wrote the manuscript.

## Competing interests

G.S. is a co-founder of Parse Biosciences, a single cell RNA sequencing company.

## Code availability

Code used for modeling and data analysis is available at https://github.com/castillohair/enhancer-design.

## Data availability

Processed cell line and mouse MPRA results can be found in **Supplementary Table 9**. Sequences of all designed enhancers along with model predictions can be found in **Supplementary Table 5** and **Supplementary Table 12**.

## References

1. Regev, A. et al. The Human Cell Atlas. eLife 6, e27041 (2017).

2. Dunbar, C. E. et al. Gene therapy comes of age. Science 359, eaan4672 (2018).

3. Santorelli, M., Lam, C. & Morsut, L. Synthetic development: building mammalian multicellular structures with artificial genetic programs. Curr. Opin. Biotechnol. 59, 130–140 (2019).

4. Toda, S., Brunger, J. M. & Lim, W. A. Synthetic development: learning to program multicellular self-organization. Curr. Opin. Syst. Biol. 14, 41–49 (2019).

5. Wang, L. et al. Sensing and guiding cell-state transitions by using genetically encoded endoribonuclease-mediated microRNA sensors. *Nat*. Biomed. Eng. 8, 1730–1743 (2024).

6. Yin, C. et al. Iterative deep learning design of human enhancers exploits condensed sequence grammar to achieve cell-type specificity. Cell Syst. 16, (2025).

7. Taskiran, I. I. et al. Cell-type-directed design of synthetic enhancers. Nature 626, 212–220 (2024).

8. Gosai, S. J. et al. Machine-guided design of cell-type-targeting cis-regulatory elements. Nature 634, 1211– 1220 (2024).

9. de Almeida, B. P. et al. Targeted design of synthetic enhancers for selected tissues in the Drosophila embryo. Nature 626, 207–211 (2024).

10. Kempynck, N. et al. CREsted: modeling genomic and synthetic cell type-specific enhancers across tissues and species. 2025.04.02.646812 Preprint at 10.1101/2025.04.02.646812 (2025).

11. Xie, Z., Wroblewska, L., Prochazka, L., Weiss, R. & Benenson, Y. Multi-Input RNAi-Based Logic Circuit for Identification of Specific Cancer Cells. Science 333, 1307–1311 (2011).

12. Jain, R. et al. MicroRNAs Enable mRNA Therapeutics to Selectively Program Cancer Cells to Self-Destruct. Nucleic Acid Ther. 28, 285–296 (2018).

13. Oesinghaus, L., Castillo-Hair, S., Ludwig, N., Keller, A. & Seelig, G. Quantitative design of cell type-specific mRNA stability from microRNA expression data. 2024.10.28.620728 Preprint at 10.1101/2024.10.28.620728 (2024).

14. Khoroshkin, M. et al. A generative framework for enhanced cell-type specificity in rationally designed mRNAs. 2024.12.31.630783 Preprint at 10.1101/2024.12.31.630783 (2024).

15. Long, H. K., Prescott, S. L. & Wysocka, J. Ever-Changing Landscapes: Transcriptional Enhancers in Development and Evolution. Cell 167, 1170–1187 (2016).

16. Heinz, S., Romanoski, C. E., Benner, C. & Glass, C. K. The selection and function of cell type-specific enhancers. Nat. Rev. Mol. Cell Biol. 16, 144–154 (2015).

17. Song, L. & Crawford, G. E. DNase-seq: A High-Resolution Technique for Mapping Active Gene Regulatory Elements across the Genome from Mammalian Cells. Cold Spring Harb. Protoc. 2010, pdb.prot5384 (2010).

18. Grandi, F. C., Modi, H., Kampman, L. & Corces, M. R. Chromatin accessibility profiling by ATAC-seq. Nat. Protoc. 17, 1518–1552 (2022).

19. Lambert, J. T. et al. Parallel functional testing identifies enhancers active in early postnatal mouse brain. eLife 10, e69479 (2021).

20. Mich, J. K. et al. Functional enhancer elements drive subclass-selective expression from mouse to primate neocortex. Cell Rep. 34, (2021).

21. Ben-Simon, Y. et al. A suite of enhancer AAVs and transgenic mouse lines for genetic access to cortical cell types. 2024.06.10.597244 Preprint at 10.1101/2024.06.10.597244 (2024).

22. Friedman, R. Z. et al. Active learning of enhancer and silencer regulatory grammar in a developing neural tissue. 2023.08.21.554146 Preprint at 10.1101/2023.08.21.554146 (2024).

23. Balsalobre, A. & Drouin, J. Pioneer factors as master regulators of the epigenome and cell fate. Nat. Rev. Mol. Cell Biol. 23, 449–464 (2022).

24. Barral, A. & Zaret, K. S. Pioneer factors: roles and their regulation in development. Trends Genet. 40, 134–148 (2024).

25. Melnikov, A. et al. Systematic dissection and optimization of inducible enhancers in human cells using a massively parallel reporter assay. Nat. Biotechnol. 30, 271–277 (2012).

26. Arnold, C. D. et al. Genome-Wide Quantitative Enhancer Activity Maps Identified by STARR-seq. Science 339, 1074–1077 (2013).

27. Zhao, S. et al. A single-cell massively parallel reporter assay detects cell-type-specific gene regulation. Nat. Genet. 55, 346–354 (2023).

28. Lalanne, J.-B. et al. Multiplex profiling of developmental cis-regulatory elements with quantitative single-cell expression reporters. Nat. Methods 21, 983–993 (2024).

29. Zhao, J. et al. MPRAbase: A Massively Parallel Reporter Assay Database. 2023.11.19.567742 Preprint at 10.1101/2023.11.19.567742 (2023).

30. Maslova, A., et al. Deep learning of immune cell differentiation. Proc. Natl. Acad. Sci. 117, 25655– 25666 (2020).

31. Atak, Z. K. et al. Interpretation of allele-specific chromatin accessibility using cell state–aware deep learning. Genome Res. 31, 1082–1096 (2021).

32. Avsec, Ž., et al. Effective gene expression prediction from sequence by integrating long-range interactions. Nat. Methods 18, 1196–1203 (2021).

33. Avsec, Ž., et al. Base-resolution models of transcription-factor binding reveal soft motif syntax. Nat. Genet. 53, 354–366 (2021).

34. Janssens, J. et al. Decoding gene regulation in the fly brain. Nature 601, 630–636 (2022).

35. Agarwal, V. & Kelley, D. R. The genetic and biochemical determinants of mRNA degradation rates in mammals. Genome Biol. 23, 245 (2022).

36. Liu, J. et al. Dissecting the regulatory logic of specification and differentiation during vertebrate embryogenesis. 2024.08.27.609971 Preprint at 10.1101/2024.08.27.609971 (2024).

37. Linder, J., Srivastava, D., Yuan, H., Agarwal, V. & Kelley, D. R. Predicting RNA-seq coverage from DNA sequence as a unifying model of gene regulation. Nat. Genet. 57, 949–961 (2025).

38. Lal, A. et al. Decoding sequence determinants of gene expression in diverse cellular and disease states. 2024.10.09.617507 Preprint at 10.1101/2024.10.09.617507 (2025).

39. Hingerl, J. C. et al. scooby: Modeling multi-modal genomic profiles from DNA sequence at single-cell resolution. 2024.09.19.613754 Preprint at 10.1101/2024.09.19.613754 (2025).

40. Pampari, A. et al. ChromBPNet: bias factorized, base-resolution deep learning models of chromatin accessibility reveal cis-regulatory sequence syntax, transcription factor footprints and regulatory variants. 2024.12.25.630221 Preprint at 10.1101/2024.12.25.630221 (2025).

41. Zheng, D. et al. Predicting the translation efficiency of messenger RNA in mammalian cells. Nat. Biotechnol. 1–14 (2025) doi:10.1038/s41587-025-02712-x.

42. Bogard, N., Linder, J., Rosenberg, A. B. & Seelig, G. A Deep Neural Network for Predicting and Engineering Alternative Polyadenylation. Cell 178, 91–106.e23 (2019).

43. Sample, P. J. et al. Human 5′ UTR design and variant effect prediction from a massively parallel translation assay. Nat. Biotechnol. 37, 803–809 (2019).

44. Karollus, A., Avsec, Ž. & Gagneur, J. Predicting mean ribosome load for 5’UTR of any length using deep learning. PLOS Comput. Biol. 17, e1008982 (2021).

45. Linder, J., Koplik, S. E., Kundaje, A. & Seelig, G. Deciphering the impact of genetic variation on human polyadenylation using APARENT2. Genome Biol. 23, 232 (2022).

46. La Fleur, A., Shi, Y. & Seelig, G. Decoding biology with massively parallel reporter assays and machine learning. Genes Dev. 38, 843–865 (2024).

47. Barbadilla-Martínez, L., Klaassen, N., van Steensel, B. & de Ridder, J. Predicting gene expression from DNA sequence using deep learning models. Nat. Rev. Genet. 1–15 (2025) doi:10.1038/s41576-025-00841-2.

48. Castillo-Hair, S. et al. Optimizing 5’UTRs for mRNA-delivered gene editing using deep learning. Nat. Commun. 15, 5284 (2024).

49. De Winter, S., Konstantakos, V. & Aerts, S. Modelling and design of transcriptional enhancers. Nat. Rev. Bioeng. 3, 374–389 (2025).

50. Meuleman, W. et al. Index and biological spectrum of human DNase I hypersensitive sites. Nature 584, 244–251 (2020).

51. Kathail, P. et al. Current genomic deep learning models display decreased performance in cell type-specific accessible regions. Genome Biol. 25, 1–22 (2024).

52. Linder, J. & Seelig, G. Fast activation maximization for molecular sequence design. BMC Bioinformatics 22, 510 (2021).

53. Linder, J., Bogard, N., Rosenberg, A. B. & Seelig, G. A Generative Neural Network for Maximizing Fitness and Diversity of Synthetic DNA and Protein Sequences. Cell Syst. 11, 49–62.e16 (2020).

54. Yanai, I. et al. Genome-wide midrange transcription profiles reveal expression level relationships in human tissue specification. Bioinformatics 21, 650–659 (2005).

55. Lundberg, S. M. & Lee, S.-I. A Unified Approach to Interpreting Model Predictions. in Advances in Neural Information Processing Systems vol. 30 (Curran Associates, Inc., 2017).

56. Lima Cunha, D., Arno, G., Corton, M. & Moosajee, M. The Spectrum of PAX6 Mutations and Genotype-Phenotype Correlations in the Eye. Genes 10, 1050 (2019).

57. Masuda, T. et al. Transcription Factor SOX9 Plays a Key Role in the Regulation of Visual Cycle Gene Expression in the Retinal Pigment Epithelium *. J. Biol. Chem. 289, 12908–12921 (2014).

58. Chew, J.-L. et al. Reciprocal Transcriptional Regulation of Pou5f1 and Sox2 via the Oct4/Sox2 Complex in Embryonic Stem Cells. Mol. Cell. Biol. 25, 6031–6046 (2005).

59. Paige, S. L. et al. A Temporal Chromatin Signature in Human Embryonic Stem Cells Identifies Regulators of Cardiac Development. Cell 151, 221–232 (2012).

60. Liu, Q. et al. Genome-Wide Temporal Profiling of Transcriptome and Open Chromatin of Early Cardiomyocyte Differentiation Derived From hiPSCs and hESCs. Circ. Res. 121, 376–391 (2017).

61. Vierbuchen, T. et al. AP-1 Transcription Factors and the BAF Complex Mediate Signal-Dependent Enhancer Selection. Mol. Cell 68, 1067–1082.e12 (2017).

62. Madrigal, P. & Alasoo, K. AP-1 Takes Centre Stage in Enhancer Chromatin Dynamics. Trends Cell Biol. 28, 509–511 (2018).

63. Maytum, A., Obier, N., Cauchy, P. & Bonifer, C. Regulation of developmentally controlled enhancer activity by extrinsic signals in normal and malignant cells: AP-1 at the centre. Front. Epigenetics Epigenomics 2, (2024).

64. Lal, A., Garfield, D., Biancalani, T. & Eraslan, G. Designing realistic regulatory DNA with autoregressive language models. Genome Res. 34, 1411–1420 (2024).

65. DaSilva, L. F. et al. DNA-Diffusion: Leveraging Generative Models for Controlling Chromatin Accessibility and Gene Expression via Synthetic Regulatory Elements. 2024.02.01.578352 Preprint at 10.1101/2024.02.01.578352 (2024).

66. Li, L. & Wunderlich, Z. An Enhancer’s Length and Composition Are Shaped by Its Regulatory Task. Front. Genet. 8, (2017).

67. Kvon, E. Z., Waymack, R., Gad, M. & Wunderlich, Z. Enhancer redundancy in development and disease. Nat. Rev. Genet. 22, 324–336 (2021).

68. Villar, D. et al. Enhancer Evolution across 20 Mammalian Species. Cell 160, 554–566 (2015).

69. Li, S., Hannenhalli, S. & Ovcharenko, I. De novo human brain enhancers created by single-nucleotide mutations. Sci. Adv. 9, eadd2911 (2023).

70. Vierstra, J. et al. Mouse regulatory DNA landscapes reveal global principles of cis-regulatory evolution. Science 346, 1007–1012 (2014).

71. Lim, F. et al. Affinity-optimizing enhancer variants disrupt development. Nature 626, 151–159 (2024).

72. Jindal, G. A. et al. Single-nucleotide variants within heart enhancers increase binding affinity and disrupt heart development. Dev. Cell 58, 2206–2216.e5 (2023).

