## Supplementary Information for "Programming human cell type-specific gene expression via an atlas of AI-designed enhancers"

### Methods

#### Data preprocessing for DHS64

ENCODE DHS Index data was downloaded from Zenodo (https://doi.org/10.5281/zenodo.3838751) as indicated in the original publication[1]. These data comprise four main parts: 1) a matrix of normalized DNase-seq read density values (i.e. continuous accessibility signals) with 3,591,898 rows corresponding to DHSs and 733 columns corresponding to DNase-seq experiments (biosamples); 2) a matrix with the same dimensions containing binary values corresponding to DHS peak calls, 3) annotations for all 3,591,898 DHSs including their genomic coordinates (e.g. chromosome, summit, start, and end positions); and 4) annotations for all 733 biosamples including germ layer, system, and organ of origin, ENCODE experiment and protocol IDs, and sequencing quality metrics such as number of reads and DNase-seq Signal Portion of Tags (SPOT) scores.

A total of 64 biosamples were selected for modeling, as indicated in the main text, representing each of the 15 tissue-selective biological components and having high read depth and DCC SPOT scores (**Supplementary Table 1**). The continuous accessibility matrix was processed by filtering columns to retain those corresponding to the 64 selected biosamples, performing quantile normalization across biosamples, and log-transforming values via $\log_{10}(x + p)$, where the pseudocount $p$ was chosen as the smallest non-zero value in each biosample. The binary accessibility matrix was similarly filtered to retain only the 64 relevant columns. DHS sequences were obtained by mapping the annotated coordinates to the GRCh38 genome assembly sequence. If the annotated DHS was longer than 500 bases, and if the start (end) position was closer than 250 bases from the summit, the end (start) position was adjusted to reach a total of 500 bases, otherwise 500 bases centered at the summit were used. DHS annotations were further augmented with the following: 1) Distance to the closest transcription start site (TSS) extracted from the Gencode v42 basic annotations, where the TSS positions were obtained by filtering annotations with "feature" = "transcript" and "gene_type" = "protein_coding" or "lncRNA", and extracting the start or end positions depending on the strand orientation. 2) Distance to the closest TSS annotated in refTSS v3.3[2]. 3) Chromatin state annotations from the "full stack" ChromHMM model[3], trained on >1000 chromatin annotation tracks across >100 cell and tissue types, and thus expected to be representative of a variety of cell types. Finally, the following DHSs were removed: 1) 1,453,085 DHSs not active in any of the 64 selected biosamples (i.e. all-zero row in the binary matrix), 2) 126,840 DHSs not in the autosomes, and 3) 31 DHSs with sequences containing non-canonical bases. The remaining 2,011,942 DHSs were used to train and evaluate our DHS64 models.

#### DHS64 model training and evaluation

DHS64 is a deep residual neural network that accepts a zero-padded, one hot-encoded input sequence of up to 500nt and predicts both the $\log_{10}$-transformed continuous accessibility signal as well as the peak call probability across all 64 selected biosamples. Predicting both data modalities was found to slightly improve performance in preliminary testing. A detailed schematic of model architecture can be found in **Supplementary Figure 1A**. Chromosome-aware data splits (i.e. training, validation, and test sets) were designed using the prtpy python package (https://github.com/coin-or/prtpy) such that each split comprised a rough 80/10/10 (train/validation/test) ratio of DHSs, and DHSs from the same chromosome were contained in the same set. Three different splits – arbitrarily named #0, #1, and #3 – were used to train three independent models (**Supplementary Table 2**). The following data augmentation techniques were used: 1) both the DHS sequence and its reverse complement were fed into the model during each epoch, and 2) left-

or right-zero padding were randomly selected for each sequence. The overall training loss was comprised of a mean squared error term between the observed and predicted continuous $\log_{10}$ accessibility signals and a binary cross-entropy term between the observed peak calls and their predicted probabilities, weighted equally. We used the Adam optimizer, a learning rate of 2e-4, and a batch size of 256. Early stopping was implemented by monitoring the validation loss with a patience parameter of 2. All training and model analysis were performed in tensorflow 2 (specific versions between 2.4 and 2.10) in python 3 (specific versions between 3.7 and 3.10). Model training was performed on Amazon AWS EC2 g5.2xlarge instances, and other model-related analyses were performed on g5.xlarge instances.

Since our primary modeling goal was to capture determinants of cell type-specific activity, we evaluated models mainly on two data sets: 1) all held-out DHSs in the test set, and 2) a subset of these containing DHSs likely to be cell type-specific enhancers selected via the following criteria: i) annotated “mean_signal” greater than 0.5. This corresponds to the mean signal across all DHS Index biosamples with a positive DHS peak call, serving as a proxy of the “strength” of this DHS when active. ii) a distance greater than 2kb from the closest gencode- or refTSS-annotated TSS, to remove promoters which tend to have strong but non-specific signals. iii) Full-stack ChromHMM annotations corresponding to the state groups “enhancers”, “weak enhancers”, and “transcribed and enhancer”. iv) Active in 10 or fewer biosamples out of the 64 selected (i.e. at most 10 “ones” in the corresponding row of the binary matrix). While the latter set is dramatically smaller (32,151 DHSs compared to 186,404 in the full test set for chromosome split #3), it is a more interesting evaluation set for our purposes. A model naively trained on all available training DHSs (n = 1,636,099 for chromosome split #3) performed well on the larger test set of all DHSs, but significantly worse on the reduced cell type-specific enhancer-like set (**Supplementary Figure 1D**). We found that restricting the training dataset to those DHSs active in 10 or fewer biosamples (n = 1,401,497) improved performance on the cell type-specific enhancer-like test set while reducing it in the larger set of all DHSs (**Supplementary Figure 1E**). More detailed analysis revealed that the latter model performed better on more cell type-specific DHSs, independently of whether they had the mean_signal, TSS, and ChromHMM enhancer annotations (**Supplementary Figure 1F-G**). Further restricting the training set to those with mean_signal, TSS, and ChromHMM annotations did not improve performance, likely because the number of training samples decreased too much (n = 278,266). With exception of **Supplementary Figure 1**, all DHS64 models in this manuscript refer to models trained on biosample-specific DHSs only. Reported prediction performance on each biosample can be found in **Supplementary Table 3**.

##### DHS64-guided design of biosample-specific enhancers

Let $n = 64$ be the total number of biosamples modeled by DHS64 and $x_{i\in\{1\ldots n\}}$ be the predicted $\log_{10}$ accessibility of a sequence for the $i$-th modeled biosample. To optimize sequences specific to one target biosample with index $i = t$ (**Figure 1E-G**, **Figure 2**), we maximize the function $x_t - \frac{1}{n}\sum_i x_i$. This is equivalent to $\left(\frac{n-1}{n}\right)\left[x_t - \frac{1}{n-1}\sum_{i\neq t} x_i\right]$, i.e. the difference between the target accessibility and the average accessibilities across the non-targets shown in **Figure 1E**, up to a constant.

To avoid generating sequences that overfit a particular model, we used two strategies. First, we optimized sequences against predictions from a “pessimistic” ensemble of two independently trained DHS64 models (data splits #1 and #3), where predictions for target biosamples to maximize were taken as the minimum across models, whereas non-target predictions to minimize were taken as their maximum. Second, after sequence generation, accessibility predictions and performance metrics were recalculated using a third "validation" DHS64 model (split #0). All data shown from designed sequence predictions (e.g. **Figure 1F-G**, **Supplementary Figure 18A** and **E**) were generated with this model. Model-predicted peak call probabilities were not used for sequence design.

We reimplemented Fast SeqProp[4] (https://github.com/castillohair/corefsp) to simplify the API and enable compatibility with TensorFlow 2. The following design parameters were used: `target_weight`: 1, `pwm_weight`: 3, `entropy_weight`: 1e-3, `learning_rate`: 1e-3, number of iterations: 2500. Similarly, we updated the DEN[5] codebase (https://github.com/castillohair/genesis) to support TensorFlow 2. We trained individual DENs for each target biosample, each of which required different parameters to at least match the predicted performance of Fast SeqProp-designed sequences (**Supplementary Table 4**). For *in silico* analysis of synthetic enhancers across all 64 modeled biosamples (e.g. **Figure 1F-G**, **Figure 3A**), 250 Fast SeqProp-designed and 250 DEN-designed enhancers per target biosample were used (**Supplementary Table 5**). For MPRAs (**Figure 2**), sequences with KpnI (GGTACC and GTACC at the 5' end) and XbaI (TCTAGA) restriction sites were removed, and a smaller subset per target biosample and design method was selected (see library design section below).

#### DHS64-guided design of enhancers with complex design objectives

To optimize enhancers specific to two or three targets (**Figure 4A-I**), where $T = \{t, u\}$ or $T = \{t, u, v\}$ is the set of target biosample indices, we maximized the objective function $\min_{i \in T} x_i + \frac{a}{m} \sum_{i \in T} x_i - \frac{1}{n-m} \sum_{i \notin T} x_i$, where $m = 2$ or 3 is the number of target biosamples and $a = 0.2$ is a constant. Maximizing a combination of the minimum and average target signal gave the best results, as maximizing only the average sometimes resulted in high signal in only one target whereas maximizing only the minimum sometimes resulted in low overall target signal values. Target pairs and triplets were chosen by sampling uniformly from the list of target cell lines used in MPRAs (**Figure 2**). 117 sequences per target pair or triplet were obtained via Fast SeqProp using the objective function just described and a similar model ensemble strategy as with single-target enhancers, followed by removing sequences with KpnI (GGTACC and GTACC at the 5' end) and XbaI (TCTAGA) restriction sites. Since all these enhancers were assayed in MPRAs, their sequences have been included in the MPRA results table (**Supplementary Table 9**).

To design sequences with tunable activity (**Figure 4J-L**), we maximized the objective function $-|x_t - x_t^*| - \frac{1}{n} \sum_{i \neq t} x_i$, where $x_t^*$ is the accessibility setpoint value on target biosample $t$. We selected 120 $x_t^*$ values uniformly spaced between biosample-specific lower and upper bounds spanning the range achieved by Fast SeqProp in the "maximize specificity" design task described above. Specifically, the lower bound was set to the minimum average prediction value that a biosample reached when optimizing sequences for all 64 targets, and the upper bound was taken as 1.5 times the maximum average value. In addition, we explicitly penalized short sequences corresponding to the KpnI (GGTACC and GTACC at the 5' end) and XbaI (TCTAGA) restriction sites. For each target cell line used in MPRAs (**Figure 2**), we designed 120 sequences with Fast SeqProp, the objective function and setpoint values just described, and a model ensemble where non-target predictions were taken from the maximum across two models as described above, but target predictions were taken from their average. The sequences of these enhancers can be found in the MPRA results table (**Supplementary Table 9**)

#### Selection of DHS-sourced enhancer controls

To perform *in silico* analysis of synthetic enhancers across all 64 modeled biosamples (e.g. **Supplementary Figure 2**, **Figure 3A**), a set of matched "enhancer-like" DHSs were selected as follows: 1) DHSs in the Index dataset were filtered for i) annotated "mean_signal" greater than 0.5, ii) a distance greater than 2kb from the closest gencode- or refTSS-annotated TSS to remove promoters, iii) full-stack ChromHMM annotations corresponding to the state groups "enhancers", "weak enhancers", and "transcribed and enhancer", iv) peak calls in a number of modeled biosamples between 1 and 10. Note that these are similar criteria as those used to construct the DHS64 test set. 2) For a given target biosample, DHSs without positive peak calls in that biosample were discarded. 3). The remaining DHSs were sorted by the difference between target biosample accessibility and average accessibility across non-targets, and the top sequences were selected. For controls used for MPRAs (**Figure 2**), DHSs were truncated to their central 145bp, and an additional filtering criterium to exclude sequences with KpnI (GGTACC and GTACC at the 5' end) and XbaI (TCTAGA) restriction sites was used.

DHS controls targeting two and three cell types (**Figure 4A-I**) were chosen similarly with the following differences: In step 2, positive peak calls in all target cell types were required; In step 3, sorting was performed using the difference between the average target and average non-target accessibilities. Note that in some cases these stringent criteria resulted in insufficient or even zero sequences (**Supplementary Figure 18B** and **F**).

Negative control DHSs were chosen by randomly sampling from the set of enhancer DHSs ("mean_signal" > 0.5, >2kb from the closest TSS, enhancer-related full-stack ChromHMM annotations) with no peak calls in any of the modeled biosamples and no KpnI and XbaI restriction sites. An additional set of negative controls was generated via dinucleotide shuffling of the DHS-sourced negative controls.

##### Design of enhancers with manually embedded TF motifs

We designed enhancers with one, two, or three different TF motifs placed at multiple copy number and orientations within putative inert sequences (**Supplementary Figure 16**). Motifs were selected due to their enrichment in DHSs specifically accessible in MPRA cell lines or their biological components compared to all other DHS64-modeled biosamples. Specifically, we used FIMO[6] (q-value threshold of 0.05) to scan DHSs with motif PWMs from the JASPAR 2022 core vertebrate non-redundant database, and selected MA0142.1 (Pou5f1::Sox2, enriched in NT2-D1 DHSs), MA0050.3 (Irf1, enriched in GM12878/lymphoid DHSs), MA0476.1 (FOS, enriched in 786-O DHSs), MA0046.2 (HNF1A, enriched in 786-O/renal DHSs), MA1638.1 and MA0681.2 (HAND2 and PHOX2, enriched in SK-N-SH DHSs), MA0712.2 and MA0668.2 (OTX2 and Neurod2, enriched in WERI-Rb1 DHSs), MA0499.2 and MA1967.1 (MYOD1 and TFAP4::FL1, enriched in SJCRH30 DHSs), MA0114.4 (HNF4A, enriched in HepG2/digestive DHSs), MA0140.2 (GATA1::TAL1, enriched in K562/myeloid DHSs), MA1105.2 (GRHL2, enriched in MCF7 DHSs), MA0041.2 (FOXD3, enriched in HeLaS3/fetal lung DHSs), and MA1134.1 (FOS::JUNB, enriched in renal and cancer DHSs). As background sequences we used two of the dinucleotide-shuffled negative controls. PWM consensus sequences were used in all cases. Spacing between motifs was between 8 and 12 bases (preferably 10 unless it created KpnI/XbaI sites). Enhancers with two or three different motifs were designed such that the cell lines where these motifs were enriched corresponded to the double and triple targets in **Figure 4**. For example, a set of enhancers were embedded with GATA1::TAL1 and HNF4A to target the K562/HepG2 pair. Most enhancers with embedded motifs, however, failed to show the predicted specificity in MPRAs, but instead supported our later results on motif combinations being required for specificity (**Figure 3B**, **Supplementary Figure 15**). **Supplementary Figure 16** shows activities of a selected subset of these enhancers, and measurements of all can be found in the MPRA results table (**Supplementary Table 9**).

#### Enhancer library design

The enhancer library used for MPRA experiments contained a large number of sequences selected or designed for specificity towards the 10 cell lines where MPRAs were conducted (**Figure 2**), along with a minority of enhancers targeting the remaining 54 modeled biosamples, and additional controls. Specifically, our library included: 1) 1370 DHS-sourced enhancers selected to target one biosample (110 per MPRA cell line + 5 per remaining biosample). 2) 1770 synthetic enhancers designed with Fast SeqProp to target one biosample (150*10 + 5*54). 3) 1500 synthetic enhancers designed with DENs to target one MPRA cell line (150*10). 4) 1200 synthetic enhancers designed with Fast SeqProp for tunable specific activity towards each MPRA cell line (120*10). 5) 378 DHS-sourced enhancers selected to target 8 pairs of MPRA cell lines (50 for each of 7 pairs, 28 for SK-N-SH + GM12878, see **Supplementary Figure 18B**). 6) 936 synthetic enhancers designed with Fast SeqProp to target the same cell line pairs (117 per pair). 7) 192 DHS-sourced enhancers selected to target 8 MPRA cell line triplets (up to 50 per triplet but often fewer, see **Supplementary Figure 18F**). 8) 936 synthetic enhancers designed with Fast SeqProp to target the same cell line triplets (117 per triplet). 9) 20 DHS-sourced negative controls. 10) 20 dinucleotide-shuffled negative controls. 11) 40 highly specific HepG2- and K562-targeted enhancers from our previous publication[7], including 20 from the SHARPR-MPRA dataset[8] we used as training data as well as 20 *de novo* designed enhancers. 12) 740 enhancers with manually embedded TF motifs. This resulted in a total of 9102 enhancers. However, the DHS control selection process sometimes resulted in identical sequences for different targets, resulting in a final number of 8989 unique enhancers. Sequences of all tested enhancers can be found in the MPRA results table (**Supplementary Table 9**).

Two barcodes, to be cloned into the 3'UTR of the reporter gene to identify enhancers from their transcripts, were used per enhancer. Barcodes were generated with the `barcode_design` package (https://github.com/feldman4/dna-barcodes/blob/master/barcode_design.py) with options `--length 10 --distance 2 --limit 50000 --exclude "GGTACC|TCTAGA|ATC$|AATAAA|ATTAAA"`. The final oligo pool had the following structure: `ACTGGCCGCTTCACTG[145nt enhancer]GGTACCTCTAGA[10nt barcode]TGCCGGACCAGGTAGAT`.

#### Enhancer library cloning

The library described above was ordered as a ssDNA oligo pool (200nt, <=18k oligos) from Twist Biosciences. Cloning into a plasmid library was performed similarly to our previous work[7]. This process includes Gibson assembly of the oligo library into the backbone of the pMPRA1 plasmid (Addgene# 49349), restriction of the resulting plasmid to separate the enhancer from the barcode, and insertion of a pMPRAdonor2-derived (Addgene# 49353) fragment containing the minP promoter and the luciferase gene via T4 ligation.

To assemble the oligo pool into the pMPRA1 backbone, we first resuspended the pool in ddH2O to 10ng/uL. 10ng were then used as PCR template with primers containing Gibson overhangs, using 25uL Phusion polymerase master mix (NEB M0531), 2.5uL 10uM primer CY01, 2.5uL 10uM primer SC_01264 (**Supplementary Table 6**), and water to 50uL, with the following thermocycler program: denature at 98°C for 30s; denature/anneal/extend at 98°C for 10s, 62°C for 30s, and 72°C for 15s for 15 cycles; and finally extend for 10 minutes. The amplified library was purified using KAPA pure beads (Roche KK8002) per manufacturer's instructions using a beads-to-product ratio of 1.5x, and resuspended in 15uL ddH2O. In parallel, pMPRA1 was digested using SfiI (NEB R0123) and the 2.5kb-long backbone was gel-purified. 158ng amplified library and 325ng pMPRA1 backbone (1-to-5 molar ratio) were assembled with the NEBuilder HiFi DNA Assembly kit (NEB E2621) in a 40uL reaction at 50°C for 60 minutes. The assembly reaction was purified with KAPA pure beads (2x bead-to-product ratio) and resuspended in 12uL. This reaction was transformed into NEB 10-beta electrocompetent cells (NEB C3020K) via two electroporations (2.5uL assembly product + 40uL cells), recovered in 1mL SOC at 37°C for 1h, and incubated in 200mL LB + Ampicillin overnight. 40uL of SOC was extracted after recovery to plate serial dilutions and estimate CFU numbers. The next day, plasmid was purified from the LB culture using the Qiagen Plasmid Maxi Kit (Qiagen 12162). The estimated CFU number was 1.53e9, or ~3.9k per library member.

To insert the minP and luciferase reporter cassette, we first digested the purified pMPRA1/library plasmid with KpnI (NEB R3142) and XbaI (NEB R0145) by preparing two identical 50uL reactions with the rCutSmart buffer and 2ug plasmid each, adding 1uL KpnI per reaction and incubating at 37°C for 2h, then adding 1uL XbaI and incubating for 37°C for 6h, followed by heat-inactivating at 65°C for 20 min. We then dephosphorylated with Antarctic phosphatase (NEB M0289) by adding 6uL of phosphatase buffer, 1uL water, and 3uL phosphatase directly to the 50uL reaction (final volume = 60uL), incubating at 37°C for 1h and heat inactivating at 80°C for 2 min. Finally, the resulting fragment was gel purified. In parallel, the pMPRAdonor2 plasmid was digested with KpnI and XbaI using an identical protocol without the Antarctic Phosphatase step, and the 1.78kb-long fragment was gel purified. Ligation was performed using T4 ligase (NEB M0202) via two identical 50uL reactions each with 2.5uL T4 ligase, 5uL 10x ligase buffer, 150.96ng pMPRA1/library-derived fragment and 187.5ng pMPRAdonor2-derived fragment (molar ratio 1-to-2, maximum ligated product: 250ng per reaction). Reactions were incubated at 16°C for 16h, 65°C for 10 min, and 4°C forever, purified with KAPA pure beads and a 1.5X beads-to-product ratio, and resuspended in 12uL water as above. This was then transformed into NEB 10-beta cells via two electroporations (2.9uL assembly product + 40uL cells each) as above. The next day, before plasmid purification, 6 glycerol stocks were prepared (800uL overnight culture + 800uL 50% glycerol) and stored at -80°C. The estimated CFU number was 1.79e9, or ~7.5k per library member. We used the Qiagen Maxi Kit with the remaining overnight culture to obtain the final purified plasmid library, hereafter referred to as library preparation DNA_20230902. At later dates, glycerol stocks were revived by thawing and aliquoting their entire contents in 200mL LB + 200uL 1000X Carbenicillin, incubating overnight, and purifiying with the Qiagen Maxi Kit as above. This resulted in library preparations DNA_20240105 and DNA_20240612.

Successful cloning and diversity of each library preparation was verified mainly by amplifying and sequencing a fragment from the plasmid library containing the enhancer region, the promoter, reporter gene, and barcode. First, a 20uL qPCR with Phusion polymerase was prepared with 10ng plasmid, 1uL 10uM primer MPRA_seq_F, 1uL 10uM primer MPRA_seq_R (**Supplementary Table 6**), 10uL Phusion master mix, and 0.2x EvaGreen (Biotium 31000). The reaction was run for 30 cycles and a cycle number before the end of exponential amplification was selected (usually ~10). The reaction was then scaled up to 100-150uL total volume without EvaGreen and ran as a regular PCR with the previously determined number of cycles. The resulting product was verified via gel electrophoresis, purified with the DNA Clean & Concentrator - 5 (Zymo D4014), and submitted to Plasmidsaurus (previously Primordium sequencing) for their long-read Premium PCR sequencing service. Each run resulted in 5000 long reads, which we verified for the expected constant regions such as promoter and reporter gene, for enhancers with the appropriate length and expected sequence, and for enhancer/barcode matching, within sequencing quality limits. In addition, we submitted for Sanger sequencing individual colonies from the plates used for CFU quantification, for both the intermediate plasmid (pMPRA1/library, 10 colonies) and final plasmid preparation (DNA_20230902, 10 colonies). We used primers MPRA_Seq_F and MPRA_Seq_R, which covered the enhancer and barcode regions, respectively, and verified that enhancer sequences and enhancer/barcode matching was as expected.

#### Cell culture, library transfection, and RNA processing

**Supplementary Table 7** lists the cell lines used in MPRA experiments (**Figure 2**), their origin, media and their source, and transfection conditions. Cells were cultured at 37°C and 5% CO2 using standard cell culturing techniques. All media was supplemented with 10% FBS (Cytiva SH30396.03) and 100 U/ml Penicillin/Streptomycin (ThermoFisher 15140122). Cell lines listed as “adherent” were detached using Trypsin-EDTA 0.25% (ThermoFisher 25200056) except for NT2-D1, which was detached with TrypLE (Fisher 12605028). During MCF7 passaging, both adherent and floating cells were retained as suggested by ATCC.

Library transfection was performed via lipofection or electroporation as indicated in **Supplementary Table 7**. Lipofection was performed using Lipofectamine 3000 (ThermoFisher L3000001) as follows: On day 1, cells were seeded at the indicated number. On day 2, lipofectamine reagent and DNA solutions were prepared as indicated, mixed, incubated for 15 minutes, and added slowly to the culture. The culture was gently swirled to mix and returned to the incubator. 4-6 hours later, media was replaced. On day 4, cells were detached and RNA extraction was performed. Electroporation on suspension cultures was performed using the Neon Transfection System (Invitrogen MPK5000) as follows: On day 1 the indicated number of cells were aliquoted, centrifuged, and resuspended twice in PBS with no Ca2+ and Mg2+. Cells were then resuspended in 100uL Resuspension Buffer R with plasmid DNA and electroporated with a 100uL Neon Tip, a Neon tube with E2 Electrolytic Buffer, and the electrical parameters in **Supplementary Table 7**. Finally, cells were placed in an appropriate container with media and FBS but no Penicillin/Streptomycin, and returned to the incubator. On day 3, RNA extraction was performed. Electroporation on adherent cells was performed as follows: On day 1 cells were seeded at the indicated number. On day 2, cells were detached, then centrifuged and resuspended twice in PBS with no Ca2+ and Mg2+, electroporated, and replated as indicated above. On day 4, cells were detached and RNA extraction was performed. Two replicates per cell line were performed. Note that the parameters in **Supplementary Table 7** correspond to a single replicate. With each transfection, two parallel cultures in a 24-well format were maintained, one of which was transfected with a GFP plasmid under identical but downscaled conditions, in order to estimate transfection efficiency via flow cytometry on the last day.

Total RNA was purified from cells using the Monarch Total RNA Miniprep Kit (NEB, T2010S. mRNA was isolated from RNA extracts using the Monarch Magnetic mRNA Isolation Kit (NEB S1550S) following the manufacturer’s instructions. Multiple parallel mRNA isolation reactions were conducted per RNA extract to keep the input RNA amount within a range of 5-10ug, and were recombined at elution step by resuspending in the same 14uL Tris.

#### Mouse retina electroporation and RNA processing

We ordered timed pregnant CD-1 mice (Charles River Labs) and collected pups at P0 for electroporation. Eyes were removed and the retinas were isolated with the lens in. The retinas were electroporated in a solution of 70uL 1x PBS with plasmid library and electroporation control in a 3:1 ratio with a concentration of 1 ug/uL. With this method, we are able to electroporate a total of 6 eyes per library solution. Electroporation was performed in a series of 5 pulses at 30 mV, 950 ms off, 50 ms on. Afterwards, lenses were removed, and retinas were cultured *ex vivo* for 11 days in a media droplet on a Whatman filter floated on explant media (10%FBS in 1:1 DMEM:F12 with 1x Pen/Strep supplemented with (l)-Glutamine). After culturing, eyes were collected, RNA was extracted using Trizol reagent (Invitrogen 15596026), and mRNA was isolated with the Monarch Magnetic mRNA Isolation Kit as above.

##### Sequencing library preparation

Samples were sequenced in four separate runs, the first of which used custom sequencing primers and a different library preparation strategy. The plasmid aliquot DNA_20230902 was sequenced with both strategies, and the resulting enhancer barcode counts were found to match strongly, showing that these two strategies produced equivalent results. **Supplementary Table 6** contains primer sequences. **Supplementary Table 8** shows the correspondence between cell line samples, replicates, sequencing run and strategy, index primers, and the DNA library corresponding to each cell sample.

For RNA samples sequenced with custom primers (HEK293T, HeLaS3, HepG2, HMC3, K562, GM12878), reverse transcription was run using Maxima H Minus Reverse Transcriptase (Thermo EP0753) as follows: the mRNA isolate was mixed with 3uL 10uM of UMI-containing primer CY05_v2, 2uL 10mM dNTP mixture, and dH2O up to 29uL, and incubated at 65°C for 5 min, then 4°C for 1 min. Next, 8uL Maxima 5x RT buffer, 1uL SUPERase-In (Invitrogen AM2694), and 2uL Maxima H Minus RT Enzyme were added and incubated at 50°C for 15 min followed 85°C for 5 min. Next, 1uL RNase I (Thermo AM2294) and 1uL RNase H (NEB M0297) were added and the mixture was incubated at 37°C for 15 min. Finally, the reaction was purified with the Zymo DNA Clean & Concentrator-5 using 7x binding buffer and eluted in 12uL dH2O. To amplify cDNA, we first ran pilot qPCR reactions to determine, for each sample, the optimal number of cycles before the end of exponential amplification: 2uL cDNA was mixed with 0.5uL 10uM i5 and i7 primers, 5uL KAPA Hifi HotStart ReadyMix (Roche 07958935001), Evagreen Dye to 1x (Biotium 31000), and water to 10uL. The mixture was incubated with the following thermocycler program: denature at 95°C for 3 min; denature/anneal/extend at 98°C for 15s, 65°C for 30s, and 72°C for 30s for 30 cycles; and finally extend at 72°C for 1 minute. Scaled up reactions were then run in a regular thermocyler with the optimal number of cycles using 10uL cDNA, 2.5uL of each primer, 25uL Kapa Hifi HotStart ReadyMix, no Evagreen, and water to 50uL. Corresponding sequencing libraries of plasmid DNA were prepared by first running a two-cycle PCR to introduce UMIs as follows: 500ng plasmid were mixed with 2.5uL 10uM of UMI-containing primer CY05_v2, 2.5uL 10uM i5 primer, 25uL 2x KAPA Hifi HotStart ReadyMix, and dH2O to 50uL. Then, the mixture was incubated with the following thermocycler program: denature at 95°C for 3 min; denature/anneal/extend at 98°C for 15s, 65°C for 30s, and 72°C for 30s for 2 cycles; and extend at 72°C for 30 seconds. The reaction was then purified with the Zymo DNA Clean & Concentrator-5 using 5x binding buffer and eluted in 10uL dH2O. Using either 50ng or 250ng of this product as template, we performed pilot qPCRs and full PCRs as above with a P5 primer (not the i5-containing primer) and the corresponding i7 primer. Two separate replicates of this plasmid library preparation workflow, starting with the UMI insertion PCR, were performed per DNA library.

For the remaining samples sequenced with standard Illumina primers, reverse transcription was performed as follows: the mRNA isolate was mixed with 1.5uL 10uM of UMI-containing primer SC_01286_RT_MPRA_stdr2, 1uL 10mM dNTP mixture, and dH2O up to 14.5uL, and incubated at 65°C for 5 min, then 4°C for 1 min. Next, 4uL Maxima 5x RT buffer, 0.5uL SUPERase-In, and 1uL Maxima H Minus RT Enzyme were added and incubated at 50°C for 15 min followed 85°C for 5 min. Next, 0.5uL RNase I and 0.5uL RNase H were added and the mixture was incubated at 37°C for 15 min. Finally, the reaction was purified with the Zymo DNA Clean & Concentrator-5 using 7x binding buffer and eluted in 22.5uL dH2O. Next, a second strand synthesis reaction was run by mixing cDNA with 2.5uL 10uM primer SC_01287_RT2_MPRA_stdr1 and 25uL 2x KAPA Hifi HotStart ReadyMix, and incubating in the thermocycler at 95°C for 3 min, 98°C for 15 s, 65°C for 30 s, and 72°C for 3 min. The product was purified with Zymo DNA Clean & Concentrator-5 using 5x binding buffer and eluted in 13uL dH2O. Pilot qPCRs and full PCRs with corresponding i5 and i7 primers were performed as above. DNA libraries were prepared as above but using primers SC_01286_RT_MPRA_stdr2 and SC_01287_RT2_MPRA_stdr1 during the UMI insertion short PCR, and i5 and i7 primers for pilot qPCRs and full PCRs.

All reaction products were gel purified in 2% agarose using the Monarch DNA Gel Extraction Kit (NEB T1020). Final library concentrations were determined using the KAPA Library Quantification Kit (Roche 07960204001) before mixing. Sequencing with custom primers was performed using an Illumina NextSeq 550 with the following settings: Read 1 (enhancer barcode): 10 cycles, read 2 (UMI): 40 cycles, Index 1: 8 cycles, Index 2: 8 cycles, custom primers: Read 1: CY08_custom_read_1, read 2: Bri022_custom_read_1, Index 1: Bri021_ind1_custom_seq, Index 2: CY09_custom_index_2. All other sequencing runs were performed at the Altius Institute using an Illumina NovaSeq 6000 with the following settings: Read 1: 151 cycles, Read 2: 151 cycles, Index 1: 8 cycles, Index 2: 8 cycles, no custom primers.

#### MPRA data analysis

Sequencing data processing was performed using the following pipeline: First, cutadapt[9] was used to filter reads based on expected constant sequences in read 2 after the UMI (bases 11-40) for the custom sequencing run, or in read 1 before and after the enhancer barcode (bases 1-23 and 33-50) and read 2 after the UMI (bases 11-50). UMIs were then clustered using starcode-umi[10]. Next, a custom python script was used to count enhancer barcodes based on exact matching to the designed library. Counts from barcodes corresponding to the same enhancer were pooled together. Finally, DESeq2[11] was used to calculate $\log_2$(RNA/DNA) values, with RNA and DNA library pairings as indicated in **Supplementary Table 8**. **Supplementary Figure 4** shows the number of reads and UMIs at each step, as well as the resulting count correlations across replicates.

All downstream data analysis was performed in python 3 (specific versions between 3.7 and 3.10). $\log_2$(RNA/DNA) values were normalized to negative controls, so that they represented the increase or decrease in activity with respect to a putative inactive enhancer in each cell line. To this end, we calculated the average $\log_2$(RNA/DNA) across cell lines for each of the 40 negative controls, and removed those with outlier activity, i.e. outside the interval (first quartile – 1.5*interquartile range, third quartile + 1.5*interquartile range). We then calculated the median $\log_2$(RNA/DNA) of the remaining 35 controls in each cell line, and subtracted this from $\log_2$(RNA/DNA) values of every enhancer. Only enhancers with a UMI count greater than 20 summed across replicates in every DNA library were used for analysis. After these filtering steps, we obtained activities for 9061 out of the intended 9102 enhancers in every cell line (**Supplementary Table 9**).

#### Transcription factor (TF) motif preprocessing and alignment

We scanned enhancers with TF motifs from the SCENIC+ motif collection[12], which compiles motif data from several sources including CIS-BP, HOCOMOCO, and JASPAR. A motif annotations table, including mappings of motifs to TF genes, was downloaded from https://resources.aertslab.org/cistarget/motif2tf/motifs-v10nr_clust-nr.hgnc-m0.001-o0.0.tbl. Motif PWM files in .cb format were downloaded from https://resources.aertslab.org/cistarget/motif_collections/v10nr_clust_public/singletons/. The motif collection was filtered and preprocessed such that each motif corresponded to one and only one TF gene as follows: 1) Motifs in the annotation table where the "description" field was different than "gene is directly annotated" or that did not have a corresponding PWM file were discarded. 2) Annotation entries corresponding to motif "metaclusters", which pointed to .cb files with multiple PWMs, were reannotated to point to an individual motif PWM. If the PWM did not exist, the entry was discarded. 3). Motifs sourced from "taipale_tf_pairs" or "tfdimers" were discarded. 4) Annotation entries with duplicated motif IDs were discarded. 5) PWMs of all remaining motifs were saved in .meme format. The final set contained 6,092 motifs from 1,237 TFs.

Motif PWMs were further clustered to reduce redundancy as follows: 1) We computed length-normalized correlations between all motif pairs using the `compare-matrices-quick` module from the `matrix-clustering_stand-alone` package (https://github.com/jaimicore/matrix-clustering_stand-alone)[13]. 2) We computed motif pair distances as 1 minus their normalized correlation. 3) We performed hierarchical clustering using `scipy` with complete linkage and an euclidean distance, and generated clusters with the `fcluster` function, the `distance` criterion, and a threshold of 7.5. This process resulted in 440 motif clusters. For motif clusters in any figure in this manuscript, a human-readable name was assigned based on the TFs that corresponded to their component motifs, prioritizing those that occurred frequently in DHSs or DHS64-designed sequences. In some cases, name suffixes were used to clarify seemingly redundant clusters. For example, the suffix "_x2" was used to denote a seemingly dimeric version of another motif cluster for the same TFs. Suffixes "(1)", "(2)", etc. denoted different clusters for the same TFs with slightly different PWMs. Motif clusters, the IDs of their member motifs, and the assigned human-readable names can be found in **Supplementary Table 10**.

Enhancers were scanned for matches to the filtered set of 6,092 motifs using FIMO[6]. For each motif, a p value was chosen in advance for a false discovery rate of 0.1 on a representative set of sequences, and re-used across all DHS64-related alignments to maintain consistency. P values were determined as follows: 1) We compiled representative sequences with i) the top 1,000 DHSs with the highest biosample-specific accessibility for each of the 64 modeled biosamples, ii) the top 1,000 specific DHSs per biosample among those with enhancer-like annotations, iii) 250 synthetic enhancers designed with Fast SeqProp and DHS64 per biosample, iv) 250 enhancers per biosample designed with DENs. Duplicates, particularly identical DHSs from different sets, were removed. 2) We ran FIMO on these sequences and the relevant motif .meme file with options `--text --no-pgc --skip-matched-sequence --thresh 1` to store all matches and their p values. 3) using the function `false_discovery_control` from scipy, we computed q values via the Benjamini-Hochberg method. 4) The smallest p value corresponding to a q value larger than 0.1 was selected. Motif scanning for analysis was performed with options `--text --no-pgc --thresh {p_val}`, where `p_val` is the previously determined p value threshold. In post processing, we merged occurrences of motifs of the same cluster if they overlapped by more than 70% of their length within the same sequence, retaining only information from the occurrence with the lowest p value. These were used in all aggregate analysis including motif enrichment and density heatmaps (e.g. **Figure 3A, Figure 4H, I, L**). For most examples of individual sequences (e.g. **Figure 3D, G, H**), overlapping matches were merged independently of their motif identity – i.e. only the most significant motif at each position is shown.

To generate **Supplementary Figure 10**, overlapping matches from motifs annotated with the same TF were merged as described above. The Human Protein Atlas Cancer Cell line dataset[14] was downloaded from https://www.proteinatlas.org/humanproteome/cell+line/data#cell_lines. Expression values were taken from the nTPM field. GTEx tissue RNA-seq data[15] was downloaded from https://www.gtexportal.org/home/downloads/adult-gtex/bulk_tissue_expression as TPM, and converted to nTPM via NOISeq[16]. The median expression of each gene across samples labeled from the same tissue was used for analysis. Motifs were matched to TFs via their gene name annotation in the SCENIC+ table.

Enhancers designed for the cardiomyocyte differentiation timecourse (**Figure 6C-G**) were scanned with the same 6,092 motifs with FIMO, using a q value of 0.02 instead of the DHS64-determined p values. Overlapping motifs corresponding to the same motif cluster were merged as above.

##### Finetuning of DHS64-MPRA

DHS64-MPRA predicts enhancer activity in 12 cell lines, 10 corresponding to DHS64 design targets (**Figure 2**) and the last two to HEK293T and HMC3 , which are not included in the DHS Index and were not considered for analysis. Three MPRA data splits were prepared, each of which had ~1/7$^{th}$ of the dataset (~1270 sequences) separated as a test set and the remaining data split 9:1 for training/validation. Each split was used to finetune one of the three independently-trained DHS64 models. Finetuning was performed in two stages: First, the output layer of the pretrained model was replaced with a linear Dense layer with as many outputs as cell lines to predict. Training was performed on this layer with the rest of the model frozen, using a learning rate switching scheduler starting at 1e-3 and decreasing 2x stepwise to 3.9e-6 once the validation loss stalled for more than 5 epochs. Finally, the rest of the model was unfrozen and training was resumed using a similar learning rate progression starting from 1e-5 and finishing with 3.9e-8. The training loss comprised two equally weighted terms: 1) MSE between observed and predicted $\log_2$FC averaged across cell lines, and 2) a cosine similarity between vectors comprised of observed and predicted cell line $\log_2$FC. The last term incentivizes the model to focus on the differences between predictions in different cell lines for a given sequence[17,18]. As with DHS64, we used the Adam optimizer and a batch size of 256. All training and model analysis were performed in tensorflow 2 (specific versions between 2.4 and 2.10) and python 3 (specific versions between 3.7 and 3.10). While DHS64-MPRA performance is very good on the tested sequences (**Supplementary Figure 13**), its generalization abilities can be limited by the sequence space covered by the finetuning data. In the case of 786-O, designed enhancers were unsuccessful and thus only a reduced number of active sequences were part of the finetuning set. Therefore, outside of our model interpretation analysis, we would use DHS64-MPRA 786-O predictions with caution.

##### Model interpretation

Nucleotide contributions towards accessibility and enhancer activity predictions were computed using the DeepExplainer module from a custom version of SHAP (https://github.com/castillohair/shap) which incorporates modifications made by the Kundaje lab to work with genomics models with updates from the base package that enable compatibility with tensorflow 2. Accessibility contributions were calculated using an ensemble of all three DHS64 models where predictions were averaged across models. Enhancer activity contributions were similarly calculated using DHS64-MPRA. As background, 10 dinucleotide-shuffled variations of a given input sequence were used.

Contributions of each motif occurrence were calculated by summing nucleotide contributions across their matching regions. Motif contributions to mingap score were calculated from enhancer activity contributions using a similar definition of the mingap score in relation to $\log_2$FC (**Supplementary Figure 14**). Heatmaps illustrating motifs that contribute to more than one cell type depending on context (**Figure 3B**, **Supplementary Figure 14**) were calculated by considering natural or DHS64-designed enhancers targeted to one cell type and assayed via MPRAs (**Figure 2**), including those with tunable activity (**Figure 4J-L**). Heatmaps comparing motif contributions to accessibility and enhancer activity (**Figure 3I**, **Supplementary**

**Figure 17**) additionally used natural and DHS64-designed enhancers targeted to multiple cell types (**Figure 4A-I**).

##### DHS733 data preprocessing and model training

We performed quantile normalization and log transformation on the entire 3,591,898 x 733 DHS Index accessibility matrix as described above for DHS64. The following DHSs were removed from all analysis: 1) 126,840 DHSs not in the autosomes, and 2) 31 DHSs with sequences containing non-canonical bases. The remaining 3,465,027 DHSs were used to train and evaluate DHS733 models.

DHS733 is a deep residual network similar to DHS64 but with three main differences: 1) 480 convolutional filters were used per layer instead of 256, 2) the output regression head predicts all 733 $\log_{10}$-transformed continuous accessibility signals, and 3) there is no matching DHS peak calling classification head (**Supplementary Figure 24A**). We trained three DHS733 models with identical train/validation/test splits as with DHS64, using similar data augmentation techniques. We used a training loss comprised of an MSE and a cosine similarity term, similar to DHS64-MPRA, to incentivize the model to focus on learning accessibility differences across biosamples without filtering out non-specific DHSs from the training set[17,18]. We used a learning rate scheduler that switched from an initial value of 2e-4 to 2e-5, then to 2e-6, and stopped training after one, three, and three epochs of no decrease in validation loss, respectively. Weights on the epoch with the lowest validation loss were retained. As with DHS64, we used the Adam optimizer, a batch size of 256, tensorflow 2, python 3, and AWS EC2 g5.xlarge instances.

##### DHS733-guided generation of the Atlas of Synthetic Human Enhancers

As with DHS64, we generated enhancer 145 nt-long sequences using Fast SeqProp and a pessimistic ensemble of DHS733 models trained on data splits #1 and #3. We used an early stopping mechanism where the loss function needed to decrease by more than 0.001 times its current absolute value over the last 50 iterations to continue. Other parameters used with Fast SeqProp were: `target_weight`: 1, `pwm_weight`: 0.3, `entropy_weight`: 1e-3, `learning_rate`: 1e-3, `n_iter_max`: 10000. As our optimization goal, we maximized the function $x_t - P^k_{i \neq t}(x_i)$, where $P^k_{i \neq t}$ denotes the $k$-th percentile of all non-target predictions. We generated 50 sequences per target using each one of four values of $k$ (85, 90, 95, and 98). Depending on the target, sequences designed with a higher $k$ could result in activity in fewer non-target biosamples, but in many cases also led to lower on-target activity and/or a higher value of the non-target to minimize (**Supplementary Figure 25B**). Designed enhancers, along with predictions from a validation model not used for sequence design (split #0), can be found in **Supplementary Table 12**. We recommend testing enhancers designed with multiple values of $k$, but if experimental throughput is limited, we recommend selecting a value of $k$ that maximizes predicted specificity across the cell types of interest. To generate the heatmap in **Figure 6B**, we used enhancers designed with only one $k$ on each design target (i.e. row), selected as the highest value where the non-target percentile value to minimize remained low (within 0.75 $\log_{10}$ accessibility units of the median across biosamples). For the *in silico* MPRA validation (**Supplementary Figure 25C-E**), we pooled all 200 enhancers designed with all four $k$ values on each target. For our analysis of differentiation stage-specific enhancers (**Figure 6C-G**), we selected one value of $k$ per design target based on predictions across differentiation time only: $k = 90$ for $t = 0$ days, $k = 95$ for $t = 5$ days, $k = 98$ for $t = 9$ days, and $k = 95$ for $t = 14$ days.

#### Supplementary Figures

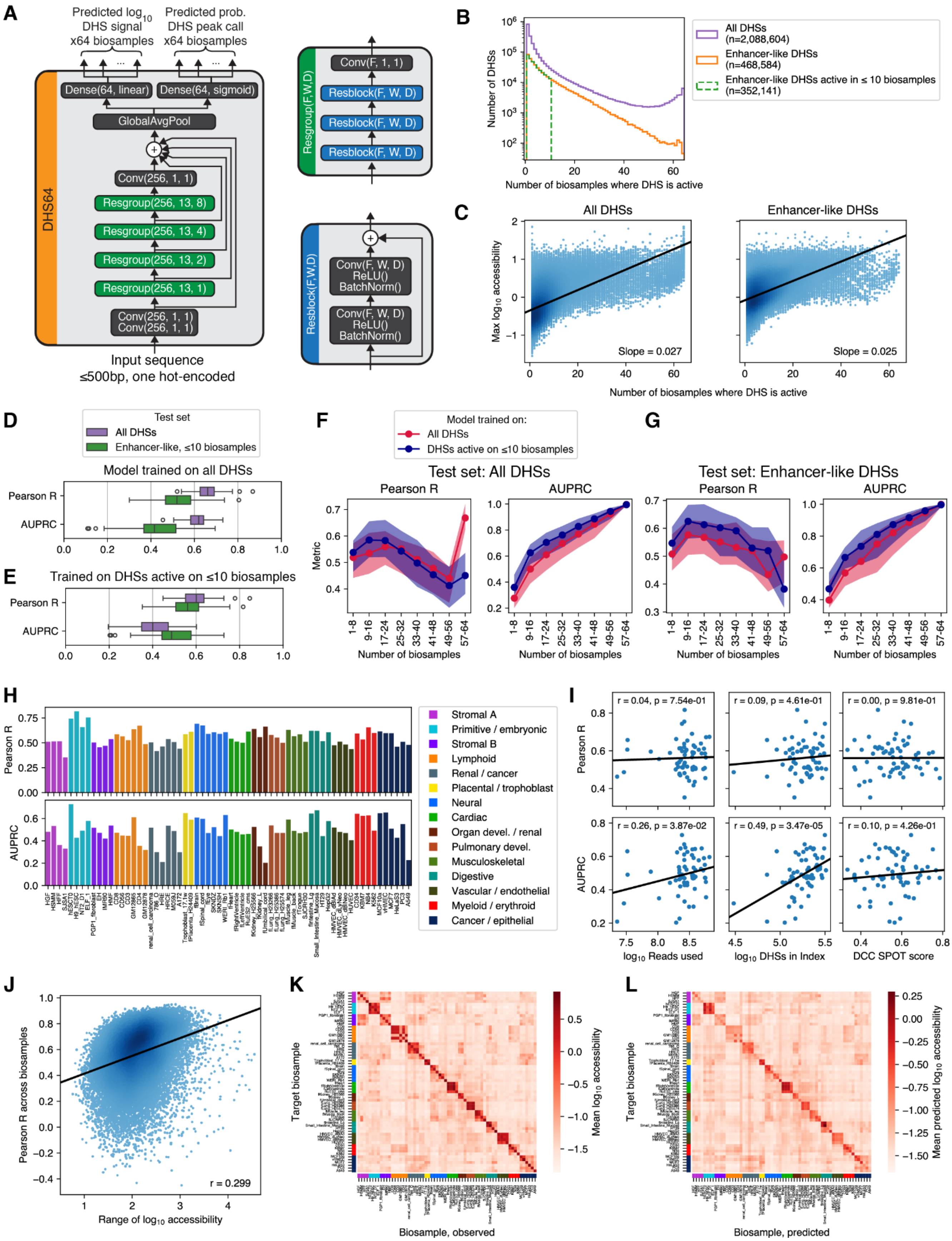

A
Predicted log10 DHS signal x64 biosamples
Predicted prob. DHS peak call x64 biosamples
Dense(64, linear)
Dense(64, sigmoid)
GlobalAvgPool
Conv(256, 1, 1)
Resgroup(256, 13, 8)
Resgroup(256, 13, 4)
Resgroup(256, 13, 2)
Resgroup(256, 13, 1)
Conv(256, 1, 1)
Conv(256, 1, 1)
DHS64
Input sequence
≤500bp, one hot-encoded
Resgroup(F,W,D)
Conv(F, 1, 1)
Resblock(F, W, D)
Resblock(F,W,D)
Conv(F, W, D)
ReLU()
BatchNorm()
B
All DHSs (n=2,088,604)
Enhancer-like DHSs (n=468,584)
Enhancer-like DHSs active in ≤ 10 biosamples (n=352,141)
Number of DHSs
Number of biosamples where DHS is active
C
All DHSs
Enhancer-like DHSs
Max log10 accessibility
Slope = 0.027
Slope = 0.025
Number of biosamples where DHS is active
D
Test set
All DHSs
Enhancer-like, ≤10 biosamples
Model trained on all DHSs
Pearson R
AUPRC
E
Trained on DHSs active on ≤10 biosamples
Pearson R
AUPRC
F
Model trained on:
All DHSs
DHSs active on ≤10 biosamples
Test set: All DHSs
Pearson R
AUPRC
Metric
Number of biosamples
G
Test set: Enhancer-like DHSs
Pearson R
AUPRC
Number of biosamples
H
Pearson R
AUPRC
Stromal A
Primitive / embryonic
Stromal B
Lymphoid
Renal / cancer
Placental / trophoblast
Neural
Cardiac
Organ devel. / renal
Pulmonary devel.
Musculoskeletal
Digestive
Vascular / endothelial
Myeloid / erythroid
Cancer / epithelial
I
r = 0.04, p = 7.54e-01
r = 0.09, p = 4.61e-01
r = 0.00, p = 9.81e-01
r = 0.26, p = 3.87e-02
r = 0.49, p = 3.47e-05
r = 0.10, p = 4.26e-01
Pearson R
AUPRC
log10 Reads used
log10 DHSs in Index
DCC SPOT score
J
Pearson R across biosamples
r = 0.299
Range of log10 accessibility
K
Target biosample
Mean log10 accessibility
Biosample, observed
L
Target biosample
Mean predicted log10 accessibility
Biosample, predicted

**Supplementary Figure 1. DHS64 model performance.** Plots were generated on models trained on chromosome split #3 (**Methods**, **Supplementary Table 2**). Results are similar for the other two DHS64 models used in this manuscript (splits 0 and 1), and their per-biosample performance metrics (equivalent to **H**) can be found in **Supplementary Table 3**. **(A)** Model architecture. Input is the sequence of a candidate DNase-hypersensitive site (DHS). Outputs are peak call probabilities (classification) and $\log_{10}$ accessibility signals (regression) for all 64 modeled cell types / biosamples. Conv(F, W, D): Convolutional layer with F filters of width W and dilation rate D. Dense(U, A): Dense layer with U units and activation function A. **(B)** Distribution of the number of biosamples in which DHSs are accessible (i.e. positive peak call) in any of the 64 modeled biosamples. Distributions are shown for all DHSs, "enhancer-like" DHSs (high "mean_signal" annotation, far from annotated transcription start sites, "enhancer" ChromHMM annotations, **Methods**), and enhancer-like DHSs active in ≤10 biosamples. Enhancer-like DHSs have a higher proportion of cell type-specific DHSs (low # biosamples) compared to broadly-accessible DHSs (high # biosamples). **(C)** Maximum accessibility signal across biosamples (y axis) for each DHSs, against the number of biosamples in which the DHS is active. Accessibility is generally higher in broadly accessible DHSs, even in the "enhancer-like" subset. Linear regression slopes shown are in units of $\log_{10}$(accessibility signal) per additional biosample. **(D)** Per-biosample performance of a model trained on all available DHSs, tested against all held-out DHSs or enhancer-like DHSs active in ≤10 biosamples. This model does worse on the latter set, possibly in part because broadly accessible DHSs have a higher signal, as shown in **(C)**, contributing disproportionally to error metrics. **(E)** Per-biosample performance of a model trained on a filtered set of DHSs active in ≤10 biosamples. There is a decrease in performance when evaluating against all held-out DHSs, but an improvement when testing on enhancer-like DHSs active in ≤10 biosamples. This is our DHS64 model, and the results reported on enhancer-like DHSs active in ≤10 biosamples correspond to **Figure 1C**. **(F)** Per-biosample performance of both models on DHSs binned by their number of active DHSs. As expected, the model trained on DHSs active in ≤10 biosamples has better performance on DHSs active in fewer biosamples (i.e. more cell type-specific). **(G)** Per-biosample performance on enhancer-like DHSs binned by their number of active DHSs. The difference in performance is more dramatic. **(H)** Regression (top) and classification (bottom) per-biosample performance for each modeled biosample. **(I)** DNase-seq quality metrics of each biosample, such as the number of sequencing reads, number of called peaks, and Signal Portion of Tags (SPOT) scores, generally do not correlate with performance metrics. The exception is the number of called DHSs with respect to AUPRC, as the floor of this metric corresponds to the fraction of positive samples. **(J)** Per-DHS performance, as measured by the Pearson R across biosamples, is higher for DHSs in which their accessibility values span a wider range (max - min across all biosamples). **(K)** Accessibility of the most specific DHSs. For each biosample, we selected 50 DHSs from the test set with the largest difference in $\log_{10}$ accessibility between the target biosample and the average across all other biosamples, and plotted their average observed accessibility as a row. **(L)** Predicted accessibility of DHSs in **(K)**.

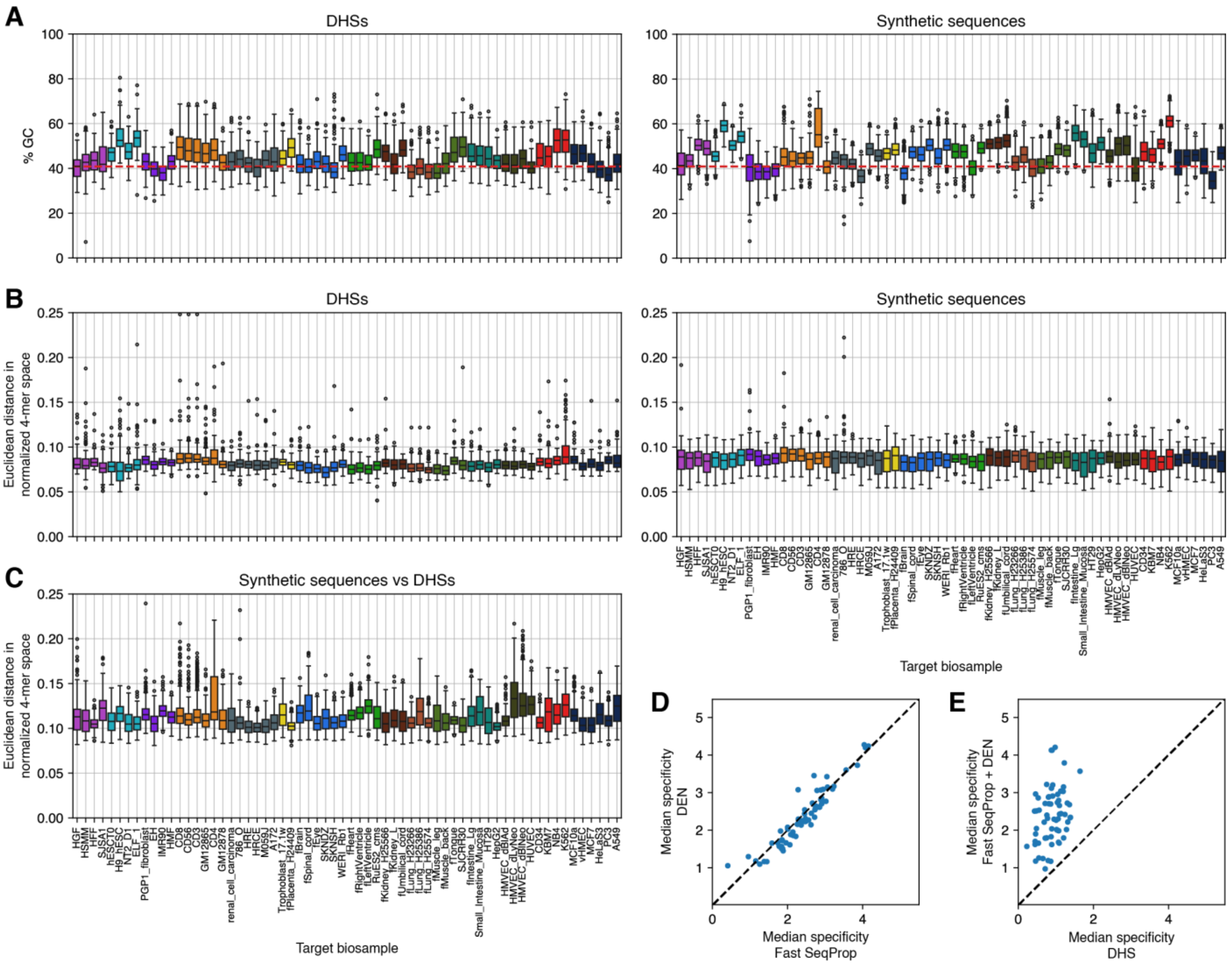


**Supplementary Figure 2. Sequence features of DHSs and synthetic enhancers.** Panels **(A)** to **(C)** compare, for each target biosample, the top 100 DHSs with the highest specificity (measured difference in $\log_{10}$ accessibility in the target biosample versus the average across non-targets) selected from DHSs with enhancer-like chromatin annotations and peak calls in ≤10 modeled biosamples, against 500 sequences (250 per method) designed via Fast SeqProp or DENs for maximal DHS64-predicted specificity (**Methods**). Boxes show the distribution of each metric across sequences selected or designed for a given target. **(A)** GC content. Red line shows the human genome GC content[19]. **(B)** Euclidean distance of 4-mer counts between sequences within their own target-specific set. **(C)** Euclidean distance of sequence 4-mer counts between DHSs and synthetic sequences targeting the same biosample. **(D)** Median predicted specificity of 250 Fast SeqProp- (x axis) and DEN- (y axis) designed sequences, for each of the 64 target biosamples. **(E)** Median predicted specificity of the top 100 enhancer-like DHSs per biosample (x axis) versus 500 NN-designed sequences (y axis).

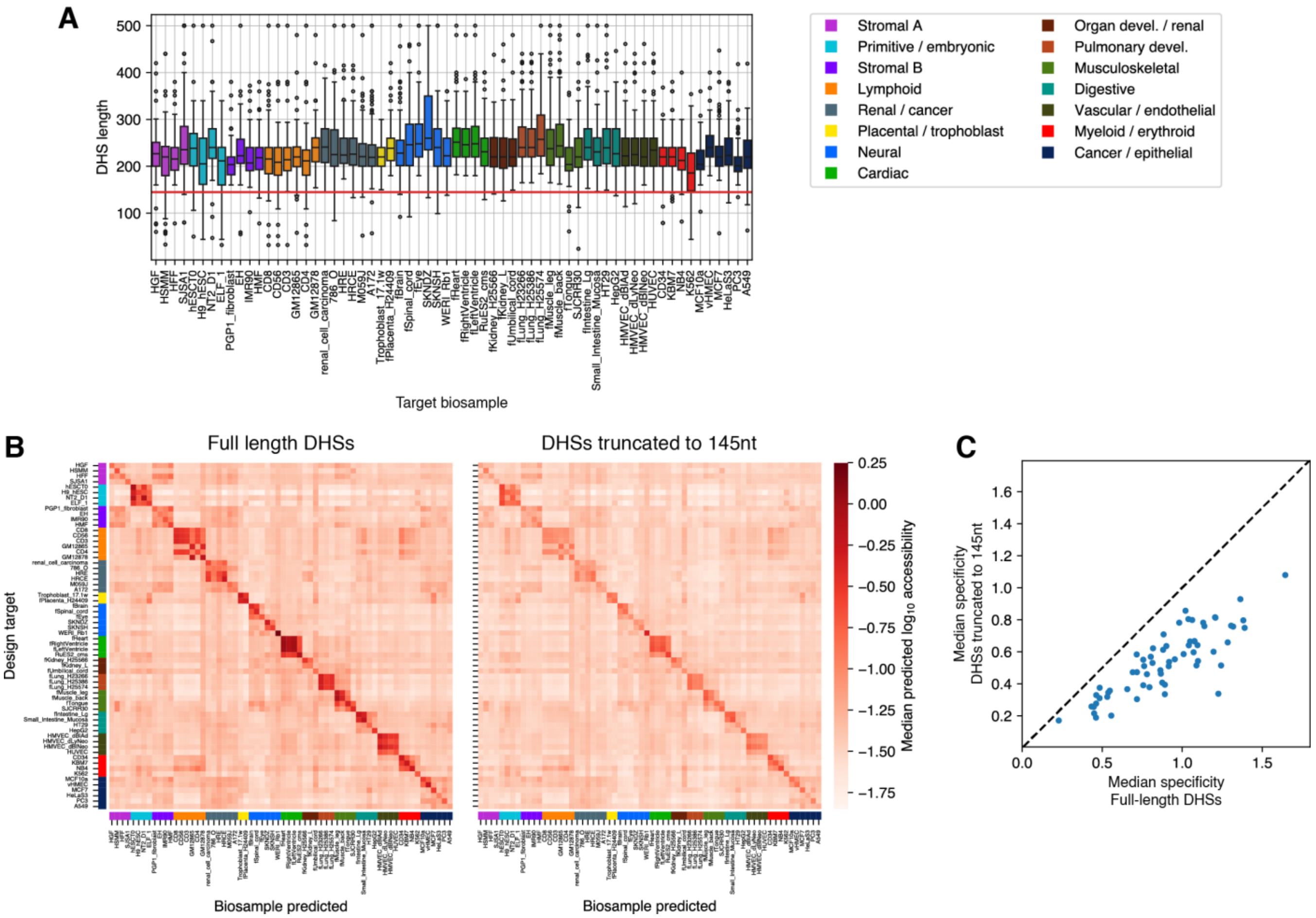


**Supplementary Figure 3. Predicted effects of truncating DHSs to 145 nt.** For each target biosample, we selected the top 100 DHSs by specificity (measured difference in $log_{10}$ accessibility in the target biosample versus the average across non-targets) from DHSs in the DNase I Index with enhancer-like chromatin annotations and peak calls in ≤10 modeled biosamples (**Methods**). **(A)** Annotated DHS lengths. The horizontal bar marks y=145 nt. **(B)** Predicted median $log_{10}$ accessibility for full-length DHSs (left) and the same DHSs truncated to 145 nt (right). Each row represents 100 DHSs for a given target. **(C)** Predicted specificity of full length (x axis) versus truncated (y axis) DHSs. Each dot represents the median predictions for DHSs targeting a biosample. The dashed line indicates y=x.

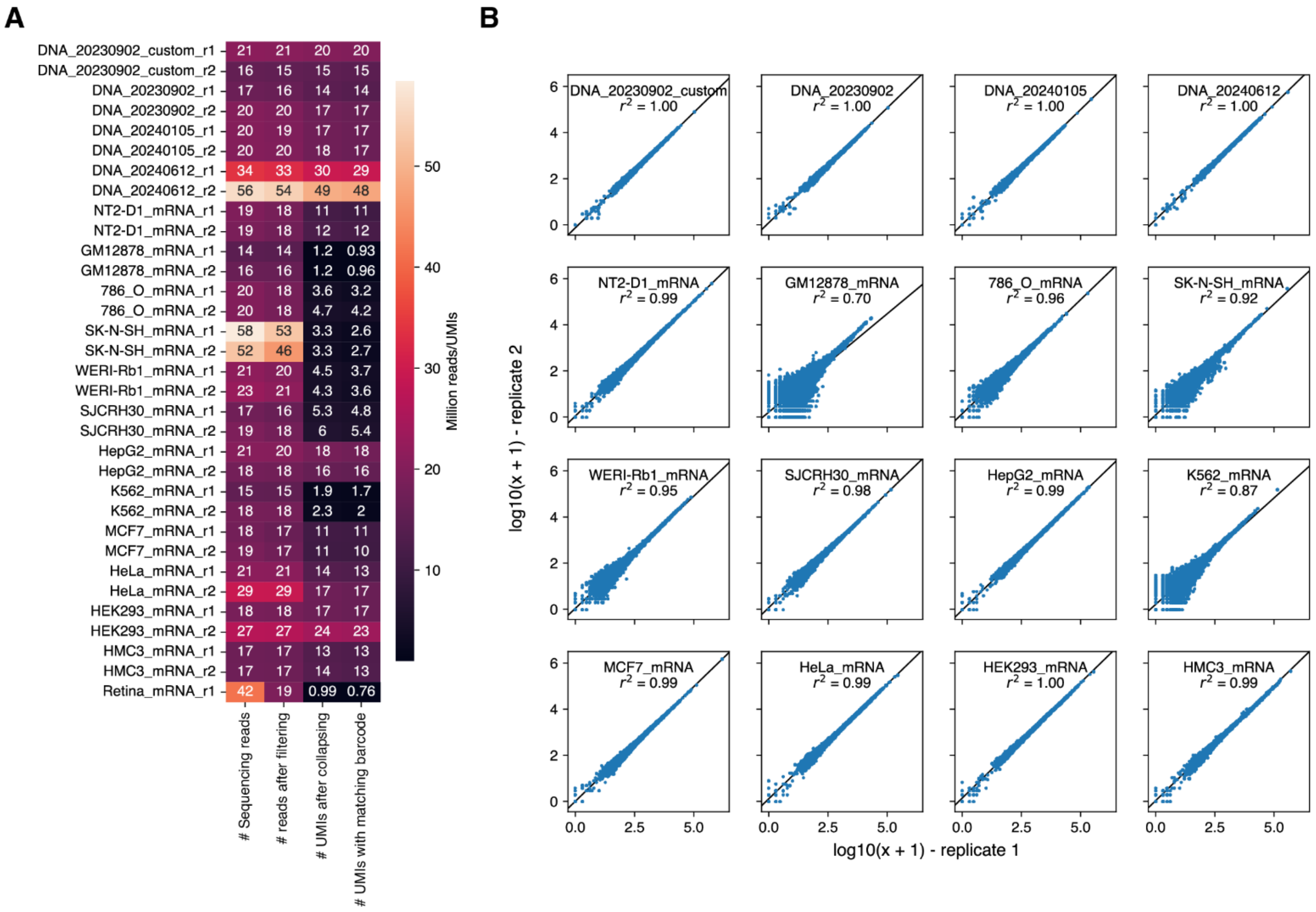


**Supplementary Figure 4. Sequencing quality metrics of all DNA and mRNA libraries sequenced for enhancer MPRAs.** Four DNA library sequencing runs were performed: three for distinct DNA library preparations (DNA_20230902, DNA_20240105, and DNA_20240612), and one where custom Illumina adaptors and sequencing primers were used (DNA_20230902_custom). Two replicates of each DNA library (_r1 and _r2) were sequenced. For each cell line, mRNA extracts from two biological replicates were sequenced. For mouse retina, only one replicate was sequenced (**Methods**). **(A)** Sequencing read counts, read counts after amplicon quality filtering, UMIs after collapse and clustering, and “useful” UMIs (barcodes matching expected sequences), in millions, for each replicate library. **(B)** Inter-replicate correlation of DNA and mRNA log-transformed counts. Least squares regression line and coefficient of determination are shown.

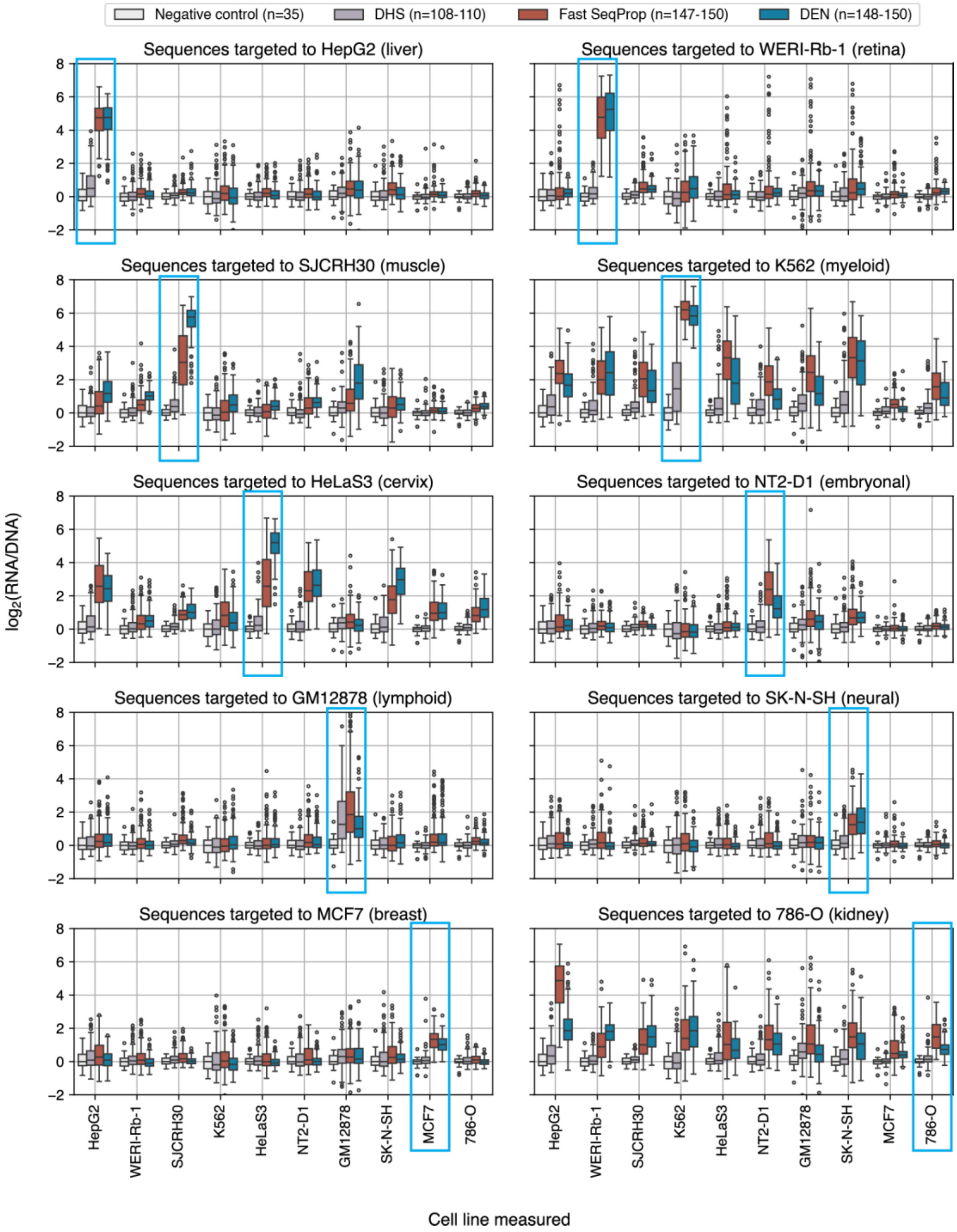


**Supplementary Figure 5. Enhancer activity measurements of sequences targeting cell lines where enhancer MPRAs were performed.** Each panel shows measurements across 10 cell lines (x axis) of sequences targeted towards one of the cell lines. Box colors indicate sequence source (DHS, Fast SeqProp, DEN, negative control). Due to sequencing depth filters (**Methods**), some panels contain slightly fewer sequences than shown in **Figure 2A**. Thus, the number of sequences per source are indicated as ranges in the legend. Negative controls are the same across all panels. Light blue rectangles highlight the target cell type in each panel.

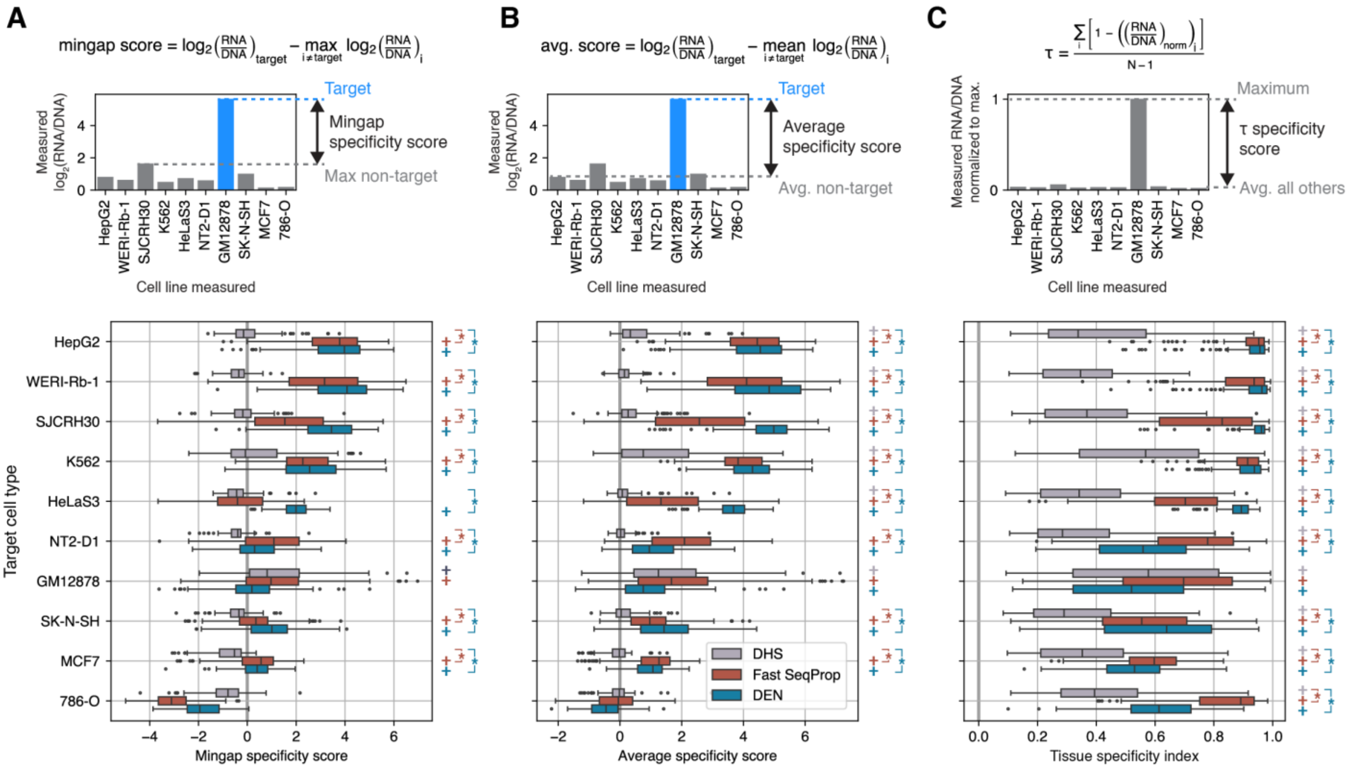


**Supplementary Figure 6. Specificity scores of deep learning-designed and DHS-sourced enhancers.** Top: mathematical definition and graphical representation of specificity score calculations given cell line enhancer measurements of a sequence. Bottom: score distribution of all sequences grouped by source and target cell type. Plus signs on the right indicate whether the median of each box is significantly positive (Wilcoxon test, one-sided, Bonferroni-corrected p-value < 0.05). Brackets with asterisks denote whether Fast SeqProp- (red) or DEN- (blue) generated sequences have significantly higher medians than DHS-derived sequences (Mann-Whitney U test, one sided, Bonferroni-corrected p-value < 0.05) **(A)** Mingap scores[20]. Compared to **Figure 2D**, here we show all synthetic and DHS-sourced sequences, with synthetic sequences further separated by design method. **(B)** Average scores, computed using the mean non-target $\log_2$FC instead of the maximum, aligning more closely with the design objective we used to optimize sequences for biosample-specific accessibility (**Figure 1E-G**) **(C)** Tissue specificity index[21], represented by τ, calculated using RNA/DNA count ratios rather than $\log_2$-transformed ratios. Note that τ is based on the maximum expression across cell types, regardless of whether it corresponds to the intended target, and can therefore be misleading when specificity is obtained towards the incorrect cell type. For example, sequences targeting 786-O show high τ values despite higher off-target expression in HepG2 and other cell lines (**Supplementary Figure 5**).

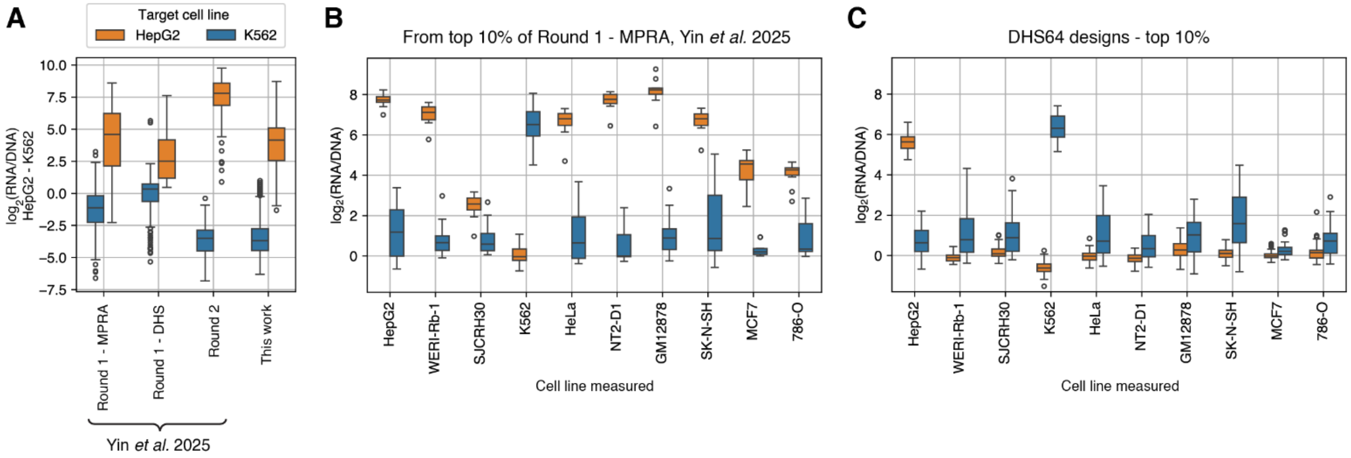


**Supplementary Figure 7. Comparison of enhancers designed with DHS64 against models of enhancer activity.** 40 sequences from our previous study[7] were re-synthesized and tested across 10 cell lines as part of the DHS64 MPRAs. These included 20 highly specific sequences from the SHARPR-MPRA dataset[8], containing measurements of ~400k genomic candidate enhancers in HepG2 and K562, and used for training CNN models in Yin *et al*. 2025, and 20 sequences from the top 10% by specificity designed with those models (Round 1-MPRA), evenly split between HepG2- and K562-targeted sequences. New measurements of these sequences in HepG2 and K562 were used to batch-correct the measurements from Yin *et al*. 2025 via simple linear regression. **(A)** Distribution of HepG2/K562 specificity (difference in $\log_2$(RNA/DNA)) for sequences designed using various methods. From Yin *et al*. 2025: “Round 1-MPRA”: sequences designed with CNNs trained on the SHARPR-MPRA dataset. “Round 1-DHS”: sequences designed with a generative adversarial network (GAN) trained to produce DHS-like sequences and a classifier of HepG2/K562 peak calls. “Round 2”: sequences designed via an iterative train/design/test approach, where CNNs trained on SHARPR-MPRA data were finetuned on both Round 1 measurements and used for a new round of designs. “This work”: sequences designed with DHS64, Fast SeqProp, and DENs to target HepG2 and K562. DHS64 HepG2 enhancers match those from the Round 1 MPRA set, whereas K562 enhancers match even those from the improved Round 2 set. **(B)** Enhancer activity of 20 re-synthesized sequences from Round 1-MPRA in Yin *et al*. 2025 measured across 10 cell lines. HepG2-targeted enhancers show high off-target activity across multiple cell lines. **(C)** Enhancer activity of a comparable set of DHS64-designed sequences (top 10% by HepG2/K562 specificity only). These exhibit reduced off-target activity, though HepG2-targeted designs show lower on-target activity than Round 1-MPRA.

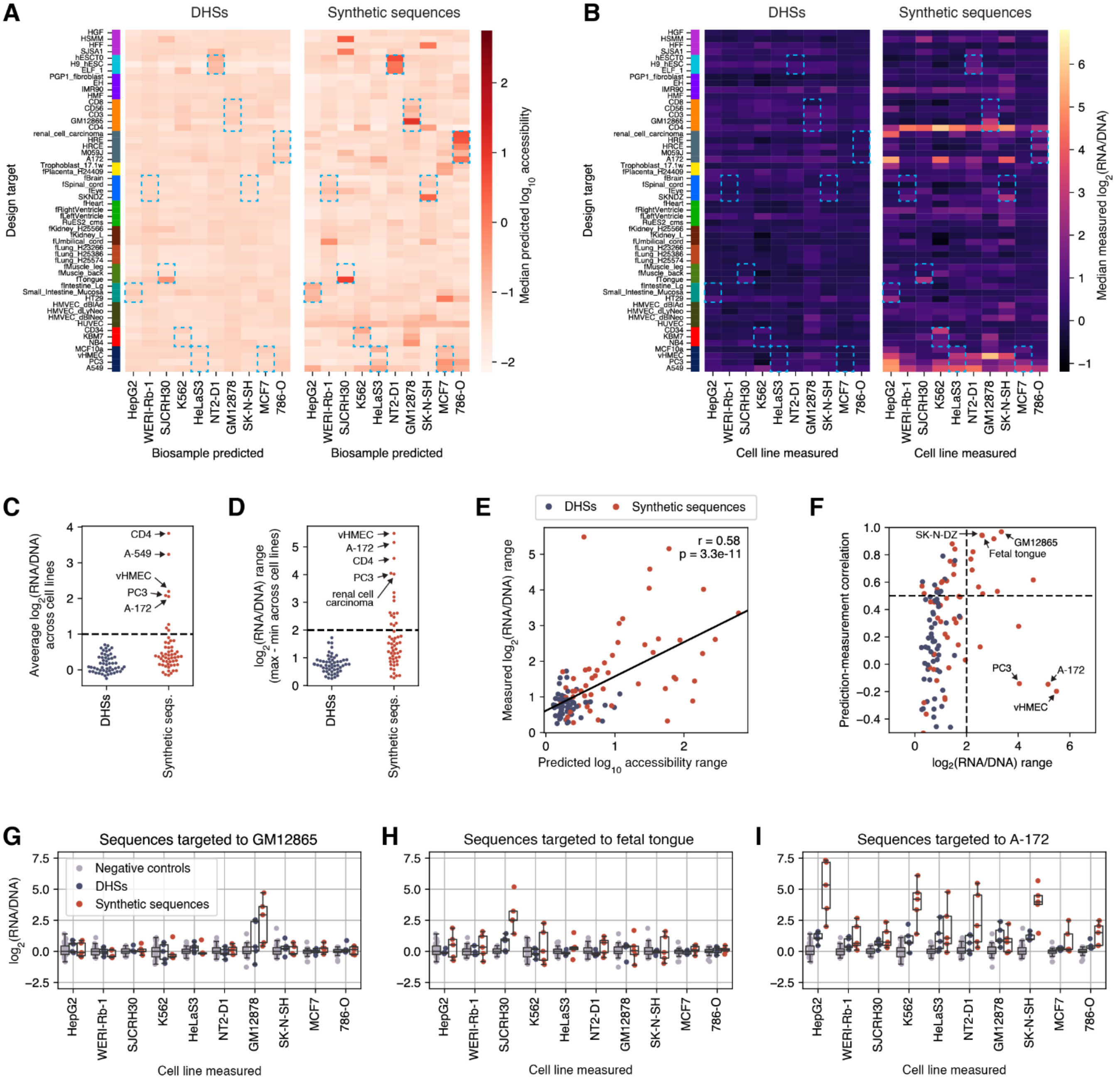


**Supplementary Figure 8. Sequences targeting DHS64 biosamples other than MPRA cell lines.** We included in our library 5 DHS-derived and 5 synthetic (Fast SeqProp-designed) sequences for each of the 54 targets. MPRA measurements were successfully obtained for all but one DHS-sourced sequence targeting HT29 cells. **(A)** DHS64-predicted $\log_{10}$ accessibility. **(B)** Measured $\log_2$(RNA/DNA). In **(A)** and **(B)**, each row corresponds to the median across sequences for a given target. Blue dashed rectangles show where cell lines (columns) correspond to the same biological component as the target biosample (rows). **(C)** Average $\log_2$(RNA/DNA) across cell lines, showing that most sequences exhibited low overall cell line activity. **(D)** $\log_2$(RNA/DNA) range (maximum - minimum) across cell lines, showing that most sequences exhibited low differential activity. In **(C)** and **(D)**, dots represent sequences targeted to each biosample, calculated from the average **(C)** or range **(D)** of each row in **(B)**. Dots marking the highest values for synthetic sequences are labeled with their biosample name. **(E)** Correlation between predicted $\log_{10}$ accessibility range and observed $\log_2$(RNA/DNA) range. x and y coordinates of each dot were calculated from the range within each row in **(A)** and **(B)**. Least squares linear regression fit, Pearson correlation, and p-value are indicated. **(F)** Prediction-measurement correlation versus differential activity ($\log_2$(RNA/DNA) range) across cell lines, showing that most cases where sequences showed differential activity were well predicted by DHS64 (upper right quadrant). Dots correspond to each target biosample. y values were obtained by correlating median predicted $\log_{10}$ accessibility with measured $\log_2$(RNA/DNA) for each target, i.e. rows in **(A)** and **(B)**. **(G-I)** Measured $\log_2$(RNA/DNA) of sequences targeting biosamples with high differential activity across cell lines. Dots correspond to individual sequences. **(G)** and **(H)** show cases where differential activity was well predicted, and high expression was present in cell lines corresponding to the same biological component as the target. **(I)** shows a case where broad high expression was not predicted.

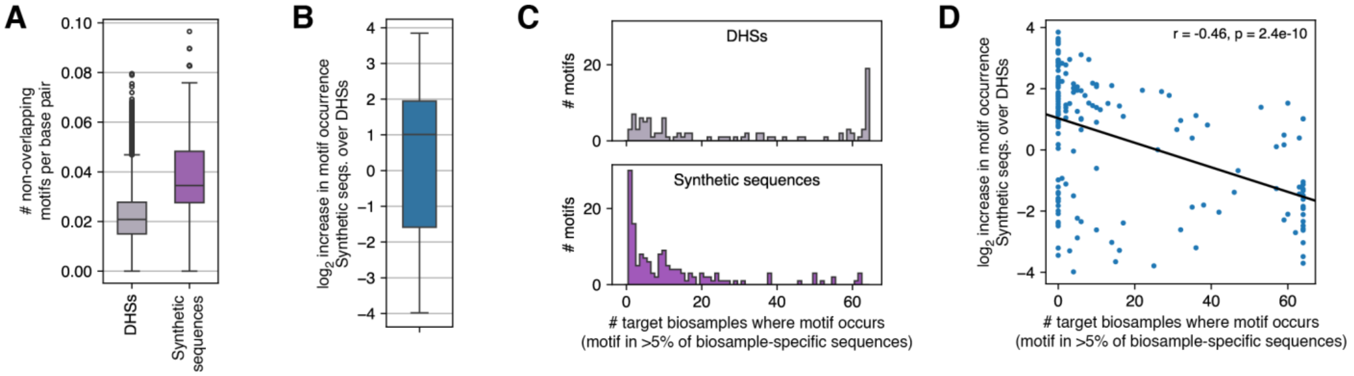


**Supplementary Figure 9. DHS64-designed sequences are enriched for biosample-specific motifs compared to DHSs.** We compare enhancer-like DHSs selected for high specific accessibility towards each DHS64-modeled biosample (1,000 per target, 64,000 total) against DHS64-designed sequences (500 per target, 32,000 total, **Methods**). **(A)** Distribution of the number of non-overlapping motifs per base pair across both sets. **(B)** Increase in motif occurrence in designed sequences relative to DHSs, measured as the $\log_2$ change in the average number of matches per sequence. Motif occurrence increases on average, although it decreases for some motifs. **(C)** Distribution of the number of target biosamples where a motif occurs (present in >5% of biosample-specific sequences), across biosample-specific DHSs (top) or DHS64-designed sequences (bottom). While the DHS distribution skews left, indicating that many motifs are used broadly, the distribution for synthetic sequences skews right, reflecting that these favor biosample-specific motifs. **(D)** Correlation between the number of biosamples where a motif is originally present in DHSs and its increase in occurrence in DHS64-designed sequences compared to DHSs.

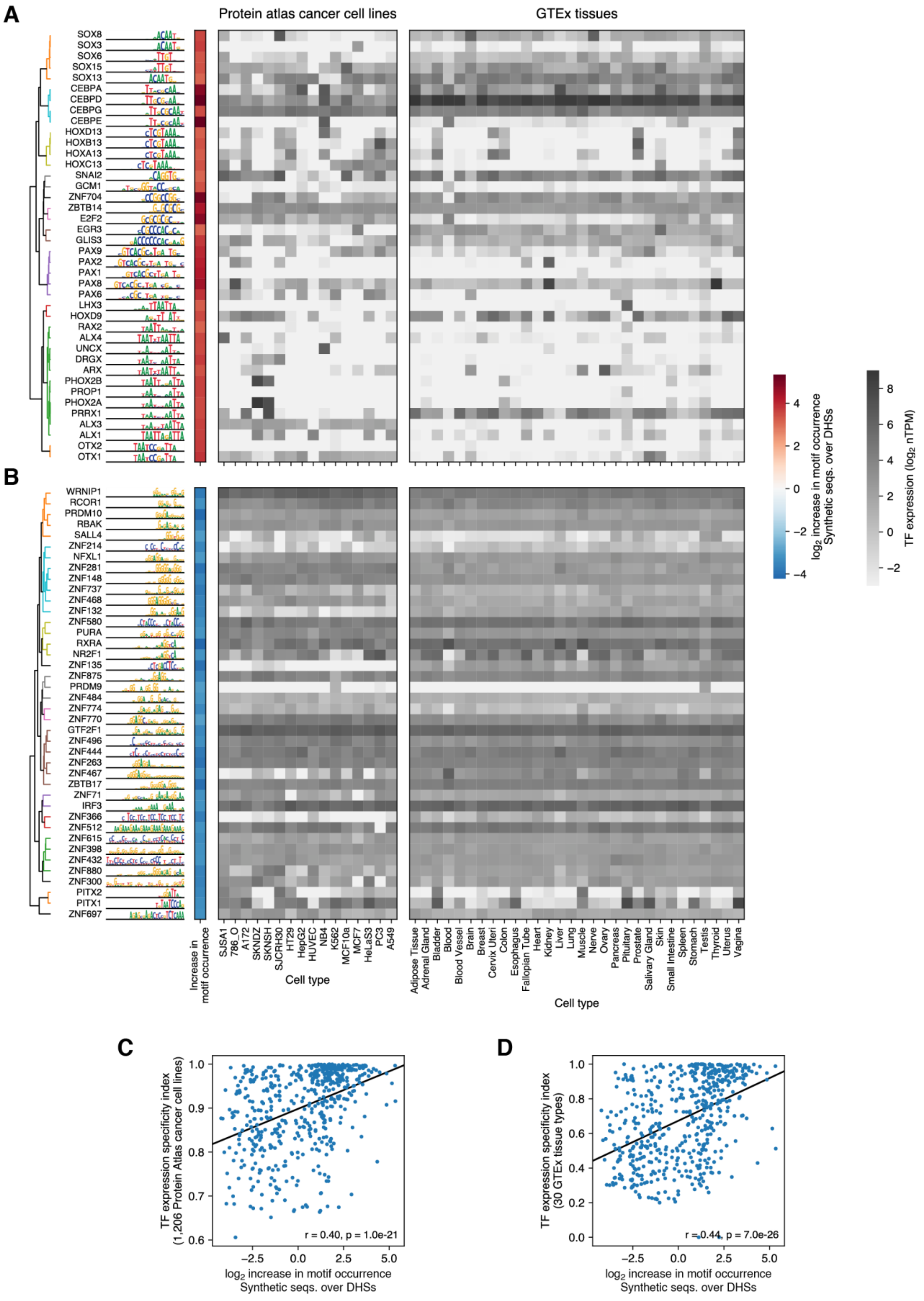
A
B
C
D
Protein atlas cancer cell lines
GTEx tissues
SOX8
SOX3
SOX6
SOX15
SOX13
CEBPA
CEBPD
CEBPG
CEBPE
HOXD13
HOXB13
HOXA13
HOXC13
SNAI2
GCM1
ZNF704
ZBTB14
E2F2
EGR3
GLIS3
PAX9
PAX2
PAX1
PAX8
PAX6
LHX3
HOXD9
RAX2
ALX4
UNCX
DRGX
ARX
PHOX2B
PROP1
PHOX2A
PRRX1
ALX3
ALX1
OTX2
OTX1
WRNIP1
RCOR1
PRDM10
RBAK
SALL4
ZNF214
NFXL1
ZNF281
ZNF148
ZNF737
ZNF468
ZNF132
ZNF580
PURA
RXRA
NR2F1
ZNF135
ZNF875
PRDM9
ZNF484
ZNF774
ZNF770
GTF2F1
ZNF496
ZNF444
ZNF263
ZNF467
ZBTB17
ZNF71
IRF3
ZNF366
ZNF512
ZNF615
ZNF398
ZNF432
ZNF880
ZNF300
PITX2
PITX1
ZNF697
Increase in motif occurrence
SJSA1
786_O
A172
SKNDZ
SKNSH
SJCRH30
HT29
HepG2
HUVEC
NB4
K562
MCF10a
MCF7
HeLaS3
PC3
A549
Cell type
Adipose Tissue
Adrenal Gland
Bladder
Blood
Blood Vessel
Brain
Breast
Cervix Uteri
Colon
Esophagus
Fallopian Tube
Heart
Kidney
Liver
Lung
Muscle
Nerve
Ovary
Pancreas
Pituitary
Prostate
Salivary Gland
Skin
Small Intestine
Spleen
Stomach
Testis
Thyroid
Uterus
Vagina
Cell type
log2 increase in motif occurrence Synthetic seqs. over DHSs
TF expression (log2 nTPM)
TF expression specificity index (1,206 Protein Atlas cancer cell lines)
r = 0.40, p = 1.0e-21
log2 increase in motif occurrence Synthetic seqs. over DHSs
TF expression specificity index (30 GTEx tissue types)
r = 0.44, p = 7.0e-26
log2 increase in motif occurrence Synthetic seqs. over DHSs

**Supplementary Figure 10. Motifs enriched in DHS64-designed sequences correspond to transcription factors (TFs) with cell type-specific expression. (A and B)** The 40 TF motifs with the greatest **(A)** or lowest **(B)** increase in occurrence in synthetic sequences compared to DHSs. For each motif, we show its position weight matrix (PWM), $\log_2$ increase in occurrence, TF expression in selected cancer cell lines (Human Protein Atlas[14]), and TF expression across tissues (GTEx[15]). For cancer cell line TF expression, only cell lines corresponding to those modeled by DHS64 are shown. **(C and D)** Correlation between motif enrichment in designed sequences and the tissue specificity index[21] of expression of the corresponding TF across 1,206 cell lines in the Human Protein Atlas dataset **(C)** or across 30 GTEx tissue types **(D)**. See **Methods** for details on analysis.

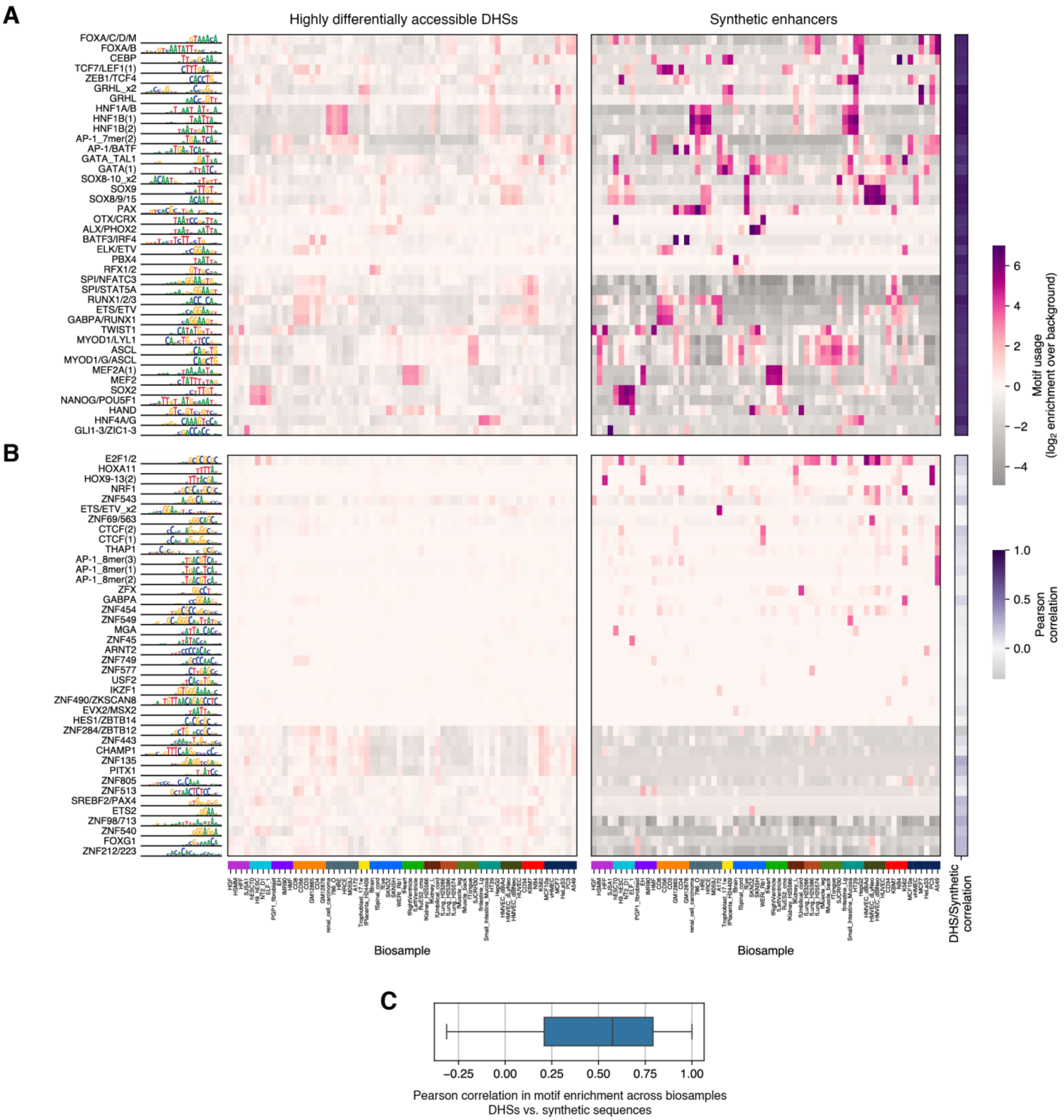


**Supplementary Figure 11. Changes in motif utilization across cell type-specific sequences. (A and B)** motif usage in highly specific enhancer-like DHSs (top 1,000 per target biosample by specificity) and in designed sequences (500 per biosample). As a measure of motif usage, for each biosample we calculate the $\log_2$ enrichment in the number of motifs per sequence compared to a background set of all DHSs specific to all other modeled biosamples, similarly to **Figure 3A**. Additionally, Pearson correlation between motif enrichment in DHSs and synthetic sequences (i.e. each row in both enrichment heatmaps) is reported, as a measure of whether a motif is used similarly across biosamples in these two sets. **(A)** The 40 motifs with the highest Pearson correlation, after filtering to remove motifs with very low usage (i.e. $\log_2$ enrichment lower than 1 in all biosamples). **(B)** The 40 motifs with the lowest correlation, after similar filtering. Motifs with low correlation can be generally classified in those that are highly depleted in synthetic sequences (bottom) and those not depleted but with true differences across biosample-specific usage (top) **(C)** Distribution of Pearson correlations, showing that usage of most motifs is similar across DHSs and synthetic sequences.

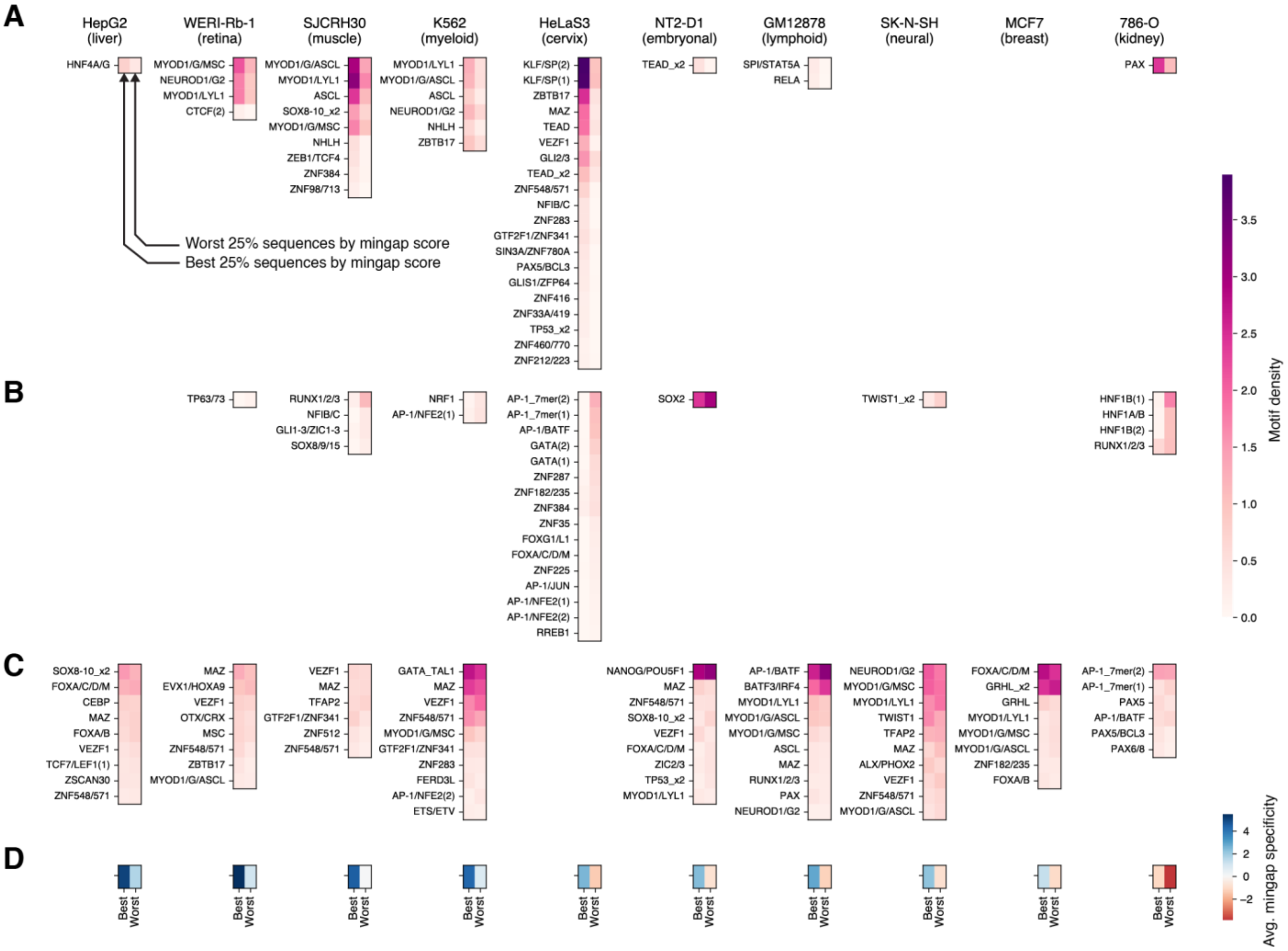


**Supplementary Figure 12. Motifs associated with the best- and worst-performing enhancers.** Each column shows data corresponding to synthetic enhancers targeted to the cell line indicated at the top. Columns within each heatmap show motif densities (average counts per sequence) in the best (left) and worst (right) 25% of enhancers for each target, as given by mingap scores calculated from MPRA measurements (**Figure 2**, **Supplementary Figure 5**, **Supplementary Figure 6**). **(A)** Motifs significantly enriched in the top 25% enhancers by mingap score. Significance was assessed via a t-test on motif counts per sequence with a Benjamini-Hochberg correction and an adjusted p value threshold of 0.05. **(B)** Motifs significantly enriched in the bottom 25% enhancers by mingap score. **(C)** Top 10 motifs by abundance across all sequences for each target, only considering those not significantly enriched in the best or worst performing sequences. **(D)** Average mingap specificity of the best and worst performing sets.

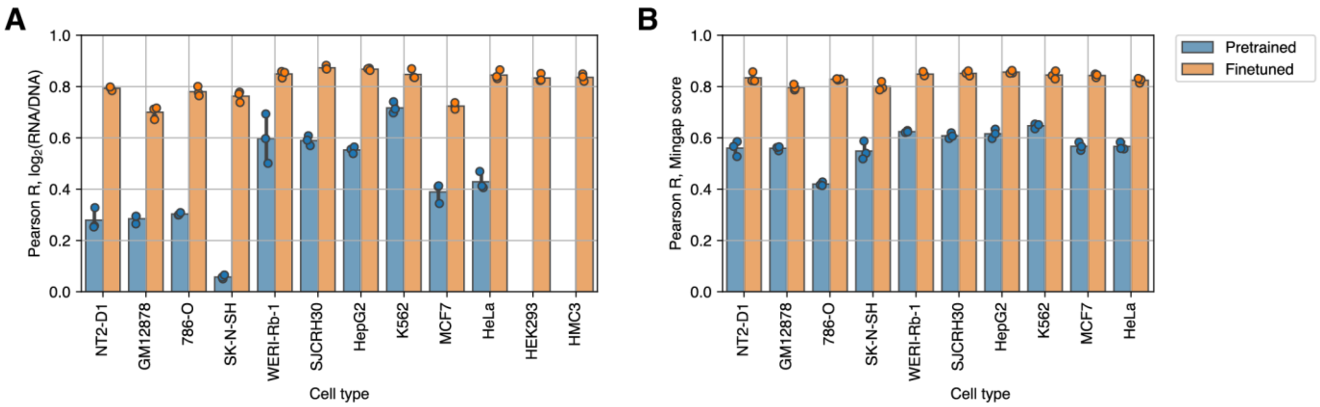


**Supplementary Figure 13. Prediction performance of DHS64-MPRA.** DHS64-MPRA was obtained by finetuning DHS64 on MPRA measurements from our entire enhancer library across 12 cell lines. These included the 10 cell lines used as enhancer design targets (**Figure 2**) as well as two additional lines, HEK293 and HMC3, which are not part of the DNase I Index dataset. Each marker shows the performance of a separate model trained on a distinct MPRA data split and evaluated on held-out data (**Methods**). Bars and error bars represent the average and standard deviation across three models. Performance is compared to the original (pretrained) DHS64 model when evaluated against MPRA data. **(A)** Correlation between model predictions and measured $\log_2$(RNA/DNA) for each cell line. Note that the pretrained model lacks outputs for HEK293 and HMC3. **(B)** Correlation between predicted and observed mingap scores, calculated on every sequence by treating every cell type as the target independently.

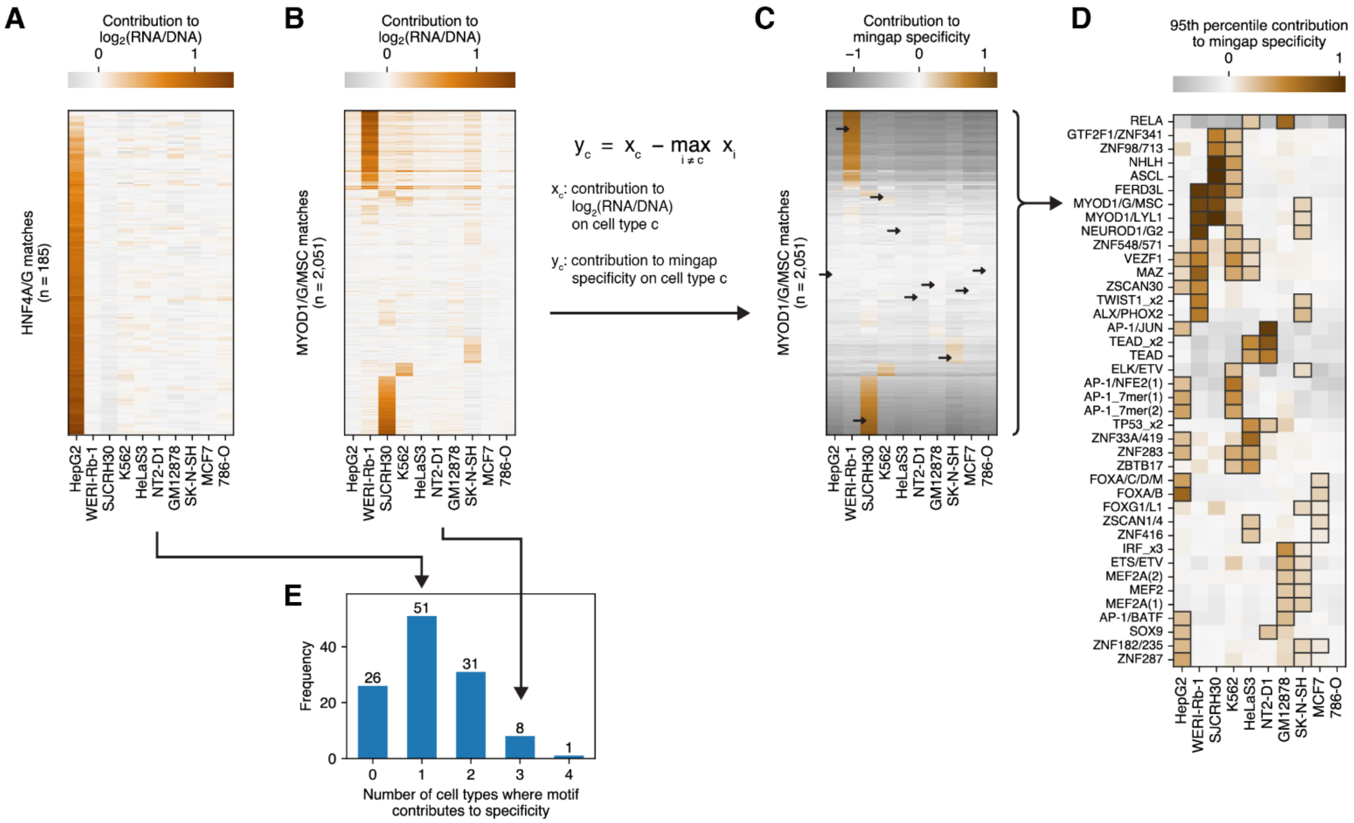


**Supplementary Figure 14. Finding motifs that contribute to specificity in multiple cell types through explainable AI.** Previously, contributions to cell line enhancer activity of every motif occurrence in every assayed sequence were calculated using our enhancer activity predictor DHS64-MPRA and SHAP[22] (**Methods**). **(A)** Enhancer activity contributions of all occurrences of the HNF4A/G motif. Most motif occurrences contribute strongly to HepG2 (liver) activity. **(B)** Contributions of all occurrences of the MYOD1/G/MSC motif, which can switch specificities between the retinal WERI-Rb-1, the muscle-derived SJCRH30, and less frequently to the myeloid K562 and the neural SK-N-SH depending on context. **(C)** Contributions of MYOD1/G/MSC occurrences to mingap specificity, calculated for each occurrence as indicated. Arrows show the 95th percentile on each cell line. **(D)** 95th percentile contributions of indicated motifs to cell line specificity. For example, values depicted for MYOD1/G/MSC correspond to where the arrows point in **(C)**. Motifs shown are those that substantially contribute to more than one cell line. A motif is considered to substantially contribute to a cell line if its 95th percentile mingap contribution is greater than 20% of the maximum across all motifs on that cell line. Squares are placed where a motif satisfies this requirement in a cell line. **(E)** Distribution of the number of cell lines that each motif contributes to specifically.

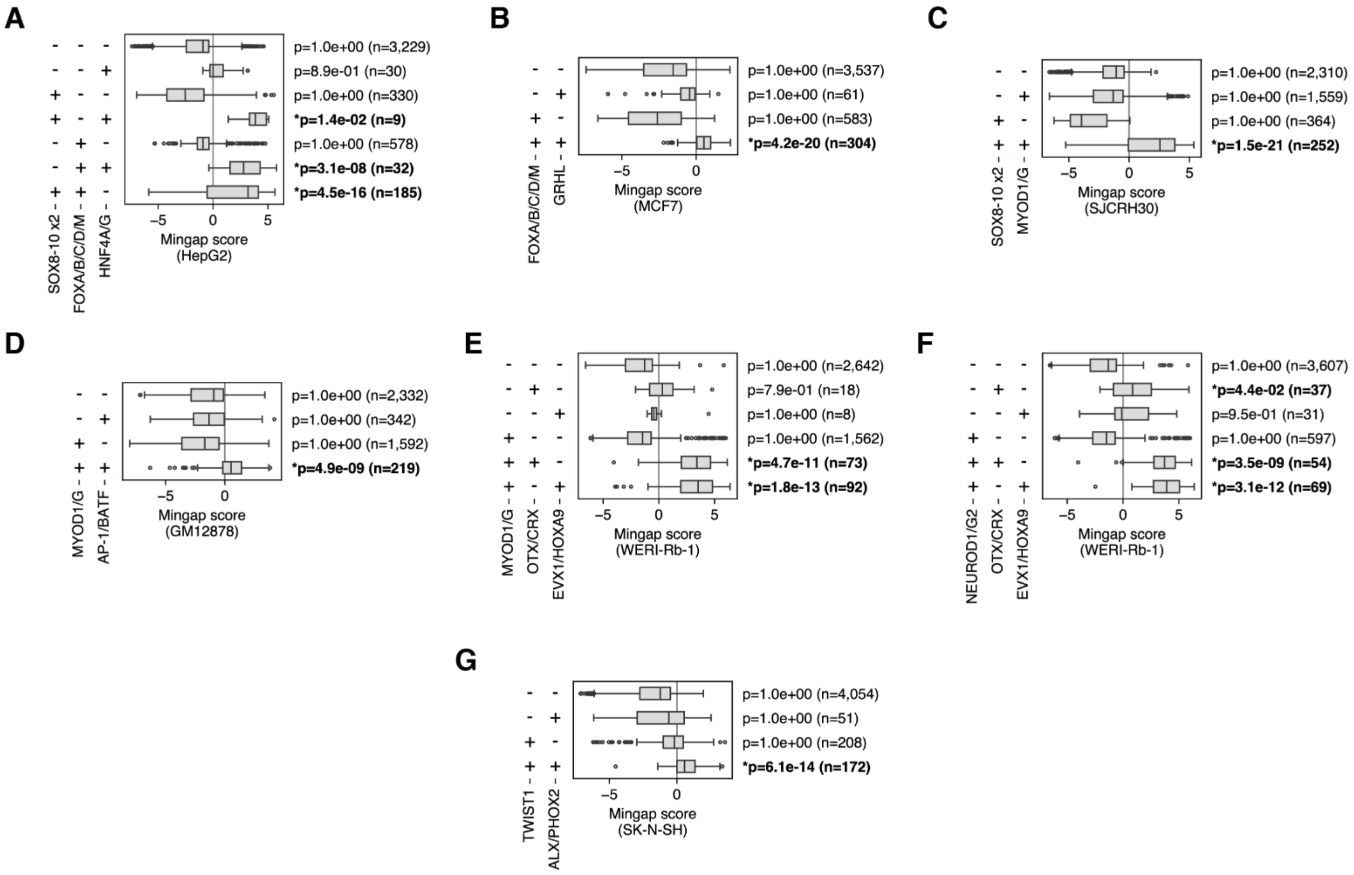


**Supplementary Figure 15. Combinations of motifs determine cell type-specific activity.** Analysis was performed on all sequences targeting each of the 10 MPRA cell lines (**Figure 2**). For each box, a Wilcoxon one-sided test was performed to determine whether the median enhancer specificity is significantly greater than zero. Asterisks and bold letters denote a p value lower than 0.05 after Bonferroni corrections. See also **Figure 3C**. **(A)** Specificity towards the liver-derived HepG2 can be achieved with pairwise combinations of FOXA/B/C/D/M, SOX8-10, and HNF4A/G. Surprisingly, HNF4A/G is not associated with well-performing enhancers in the absence of these partners. **(B)** FOXA/B/C/D/M is also associated with specificity towards the breast-derived MCF7 when present along with GRHL. **(C)** The combination of MYOD1/G and SOX8-10 motifs is associated with specificity towards the muscle-derived SJCRH30, but not either motif on their own. **(D)** MYOD1/G and AP-1/BATF are associated with specificity towards the lymphoid-derived GM12878 **(E)** MYOD1/G can be combined with OTX/CRX and EVX1/HOXA9 for specificity towards the retina-derived WERI-Rb-1. **(F)** Similarly with NEUROD1/G2 and OTX/CRX or EVX1/HOXA9. **(G)** TWIST1 and ALX/PHOX2 combined are associated with specificity towards the neural SK-N-SH.

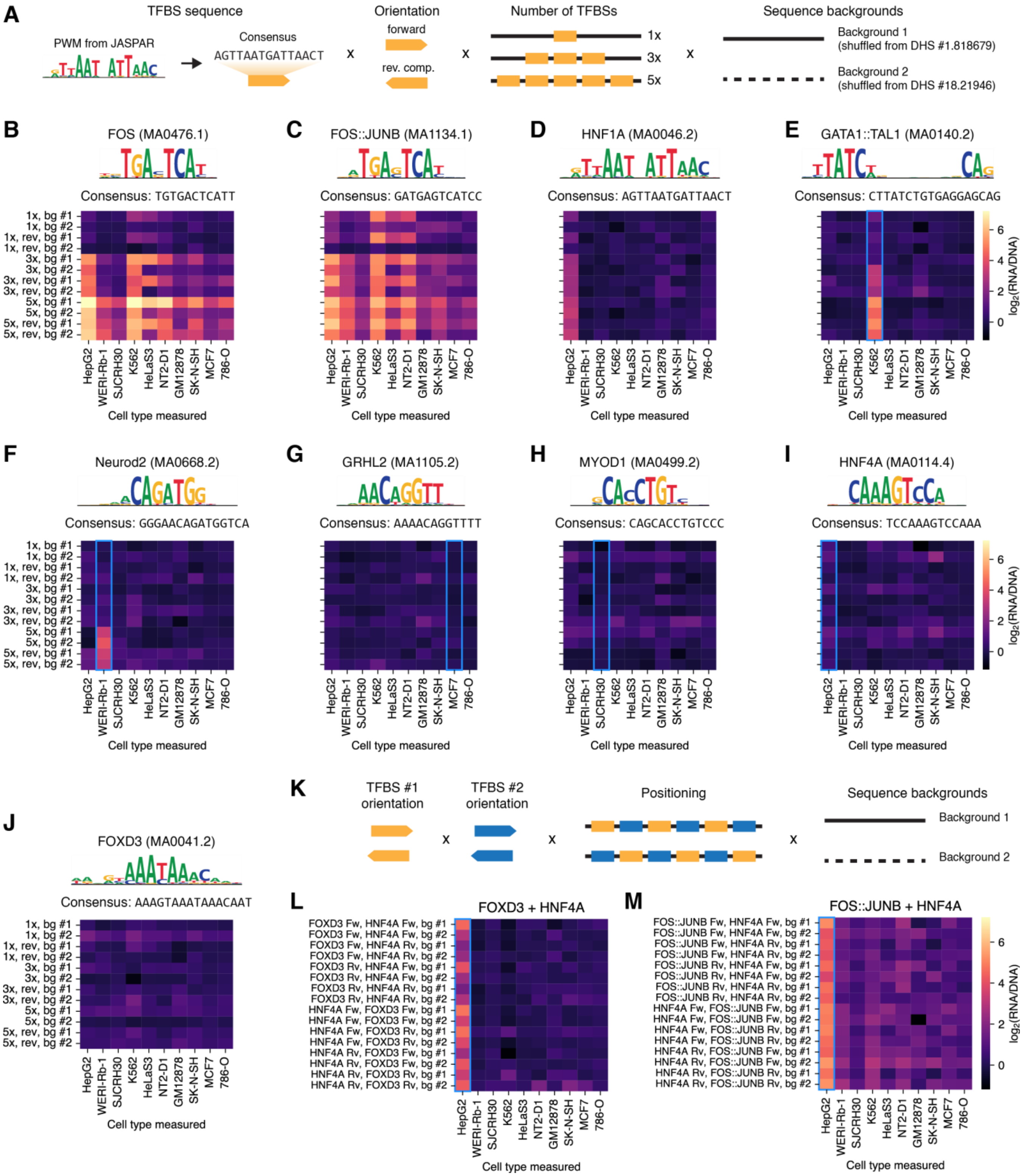


**Supplementary Figure 16. Handcrafted sequences with embedded TF motifs reveal determinants of cell type-specificity. (A)** Design of sequences with one or multiple copies of a single TF motif. Starting from PWMs from the JASPAR 2022 vertebrate database, we embedded 1, 3, or 5 copies of their consensus sequences into two dinucleotide-shuffled negative controls (**Methods**). **(B and C)** Enhancers with FOS and FOS::JUNB TFBSs drive gene expression in multiple cell types, starting from 3 TFBS copies. **(D)** Enhancers with up to 5 copies of HNF1A drive gene expression in HepG2 only. **(E)** Enhancers with the composite GATA1::TAL1 drive K562-

specific expression. **(F)** Enhancers with 5 copies of Neurod2 drive WERI-Rb1-specific expression. **(G-I)** Enhancers with GRHL2, MYOD1, and HNF4A alone do not drive gene expression in any cell line. Blue rectangles indicate cell lines where expression was expected. **(J)** FOXD3 does not drive gene expression in isolation. **(K)** Design of sequences with multiple copies of two different TF motifs (**Methods**). **(L)** Combinations of FOXD3 and HNF4A drive HepG2-specific expression. **(M)** Combinations of FOS::JUNB and HNF4A drive HepG2-specific expression, although with a higher off-target background.

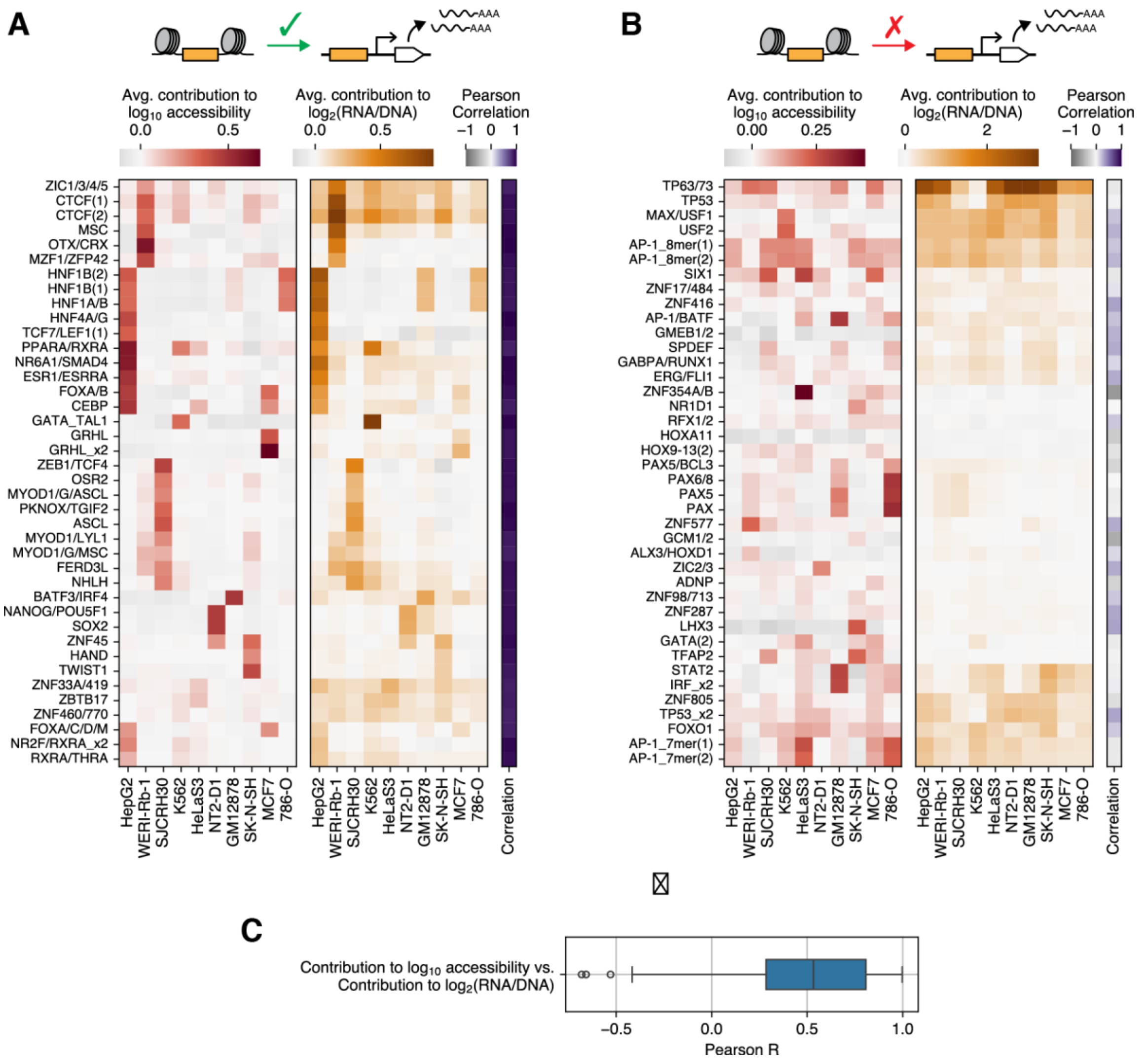


**Supplementary Figure 17. Relationship between motif contributions towards accessibility and enhancer activity. (A and B)** For the motifs shown, each panel displays: (1) the average motif contribution to predicted $\log_{10}$ accessibility on each assayed cell line (via DHS64), (2) the average contribution to predicted enhancer activity ($\log_2$(RNA/DNA), via DHS64-MPRA), and (3) Pearson correlation between cell line contributions (i.e. rows in the accessibility vs. enhancer activity heatmaps), as a measure of how well contributions to accessibility transferred to enhancer activity. **(A)** The 40 motifs with the highest Pearson correlation, after filtering to remove motifs with very low contribution (absolute contribution lower than 0.1 in all cell lines for both accessibility and enhancer activity). **(B)** The 40 motifs with the lowest Pearson correlation, after similar filtering. **(C)** Distribution of Pearson correlations for all motifs identified in any sequence.

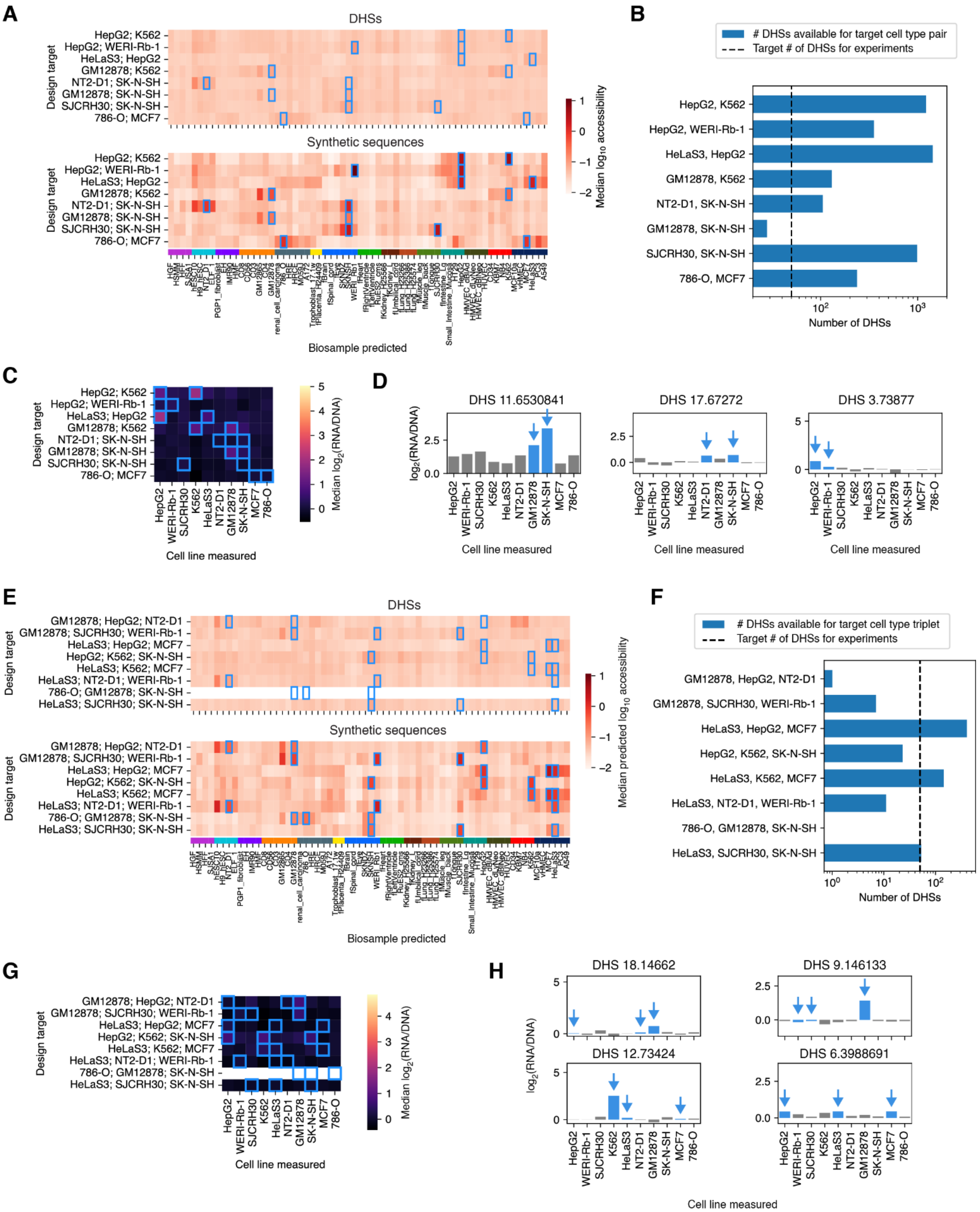
A
DHSs
Synthetic sequences
Design target
HepG2; K562
HepG2; WERI-Rb-1
HeLaS3; HepG2
GM12878; K562
NT2-D1; SK-N-SH
GM12878; SK-N-SH
SJCRH30; SK-N-SH
786-O; MCF7
Median log10 accessibility
Biosample predicted
B
### DHSs available for target cell type pair
Target # of DHSs for experiments
HepG2, K562
HepG2, WERI-Rb-1
HeLaS3, HepG2
GM12878, K562
NT2-D1, SK-N-SH
GM12878, SK-N-SH
SJCRH30, SK-N-SH
786-O, MCF7
Number of DHSs
C
Median log2(RNA/DNA)
Cell line measured
HepG2
WERI-Rb-1
SJCRH30
K562
HeLaS3
NT2-D1
GM12878
SK-N-SH
MCF7
786-O
D
DHS 11.6530841
DHS 17.67272
DHS 3.73877
log2(RNA/DNA)
Cell line measured
E
DHSs
Synthetic sequences
Design target
GM12878; HepG2; NT2-D1
GM12878; SJCRH30; WERI-Rb-1
HeLaS3; HepG2; MCF7
HepG2; K562; SK-N-SH
HeLaS3; K562; MCF7
HeLaS3; NT2-D1; WERI-Rb-1
786-O; GM12878; SK-N-SH
HeLaS3; SJCRH30; SK-N-SH
Median predicted log10 accessibility
Biosample predicted
F
### DHSs available for target cell type triplet
Target # of DHSs for experiments
GM12878, HepG2, NT2-D1
GM12878, SJCRH30, WERI-Rb-1
HeLaS3, HepG2, MCF7
HepG2, K562, SK-N-SH
HeLaS3, K562, MCF7
HeLaS3, NT2-D1, WERI-Rb-1
786-O, GM12878, SK-N-SH
HeLaS3, SJCRH30, SK-N-SH
Number of DHSs
G
Design target
Median log2(RNA/DNA)
Cell line measured
H
DHS 18.14662
DHS 9.146133
DHS 12.73424
DHS 6.3988691
log2(RNA/DNA)
Cell line measured

**Supplementary Figure 18. Properties of control DHSs selected to target multiple cell types. (A-D)** show data from control DHSs selected to target two cell types, whereas **(E-H)** show data on DHSs targeting three. **(A and E)** DHS64-predicted $\log_{10}$ accessibility for control DHSs (top), with rows representing the median of up to 50 sequences targeting each cell line pair or triplet. Predictions for designed sequences (bottom) are included for comparison. **(B and F)** Number of DHSs in the DNase I Index meeting our filtering criteria (enhancer-like chromatin annotations, peak calls in the target cell lines, peak calls in ≤10 of the 64 modeled biosamples) for each target. **(C and G)** Median MPRA-assayed expression of control DHSs. For each target pair and triplet, only sequences in the top 20% by mingap score were considered. Colorbar ranges are matched to **Figure 4A** and **D**. **(D and H)** DHSs with the highest mingap score for the indicated targets pairs or triplets. Target and non-target cell lines are indicated with blue and gray bars respectively, with blue arrows highlighting targets when they are too short to see. Targets and y axis ranges are matched to **Figure 4C** and **F**.

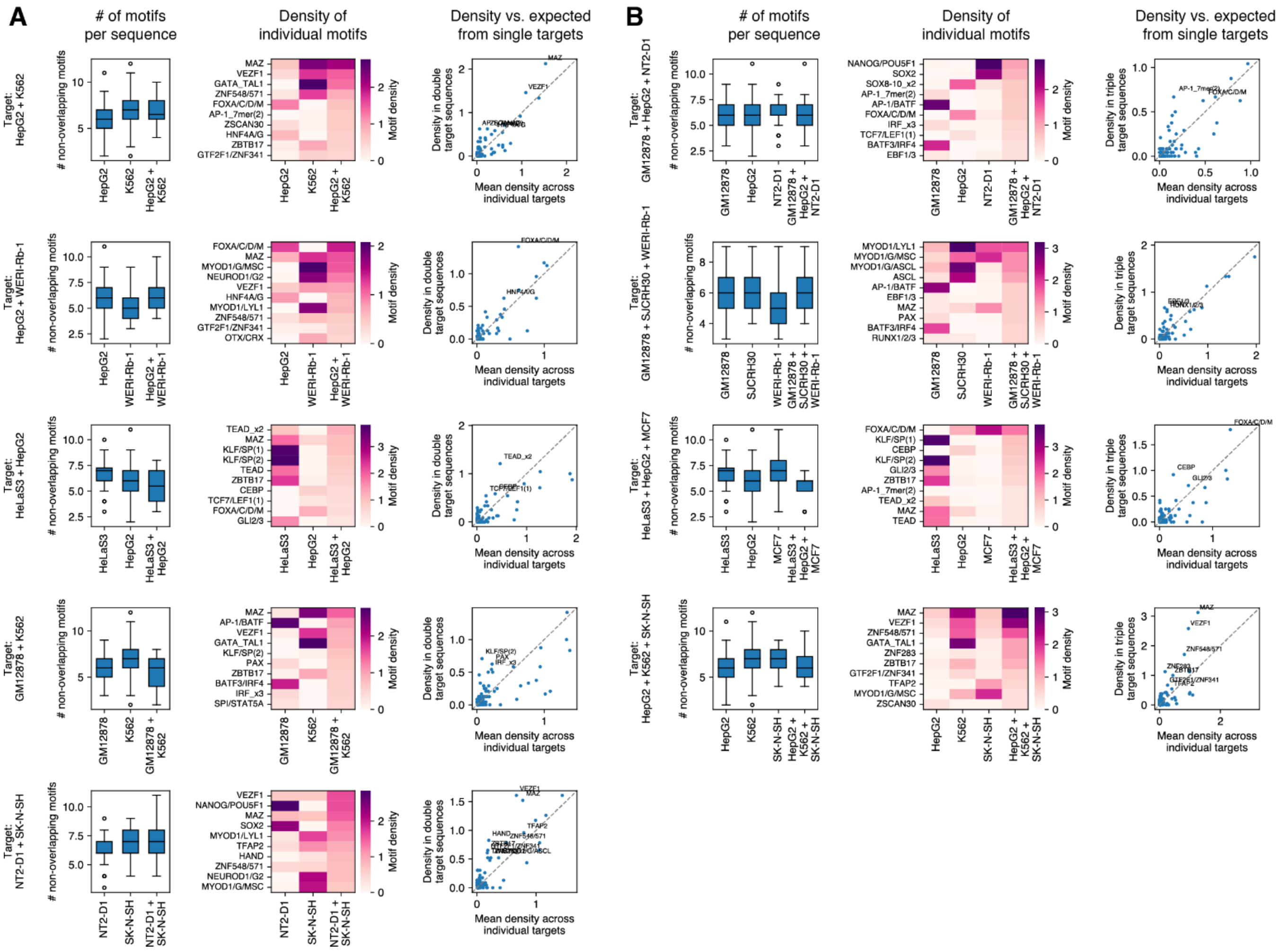


**Supplementary Figure 19. Synthetic enhancers targeting multiple cell types mostly borrow their regulatory grammar from their single-target counterparts.** Each row compares synthetic enhancers targeting two **(A)** or three **(B)** cell types with those designed for each of their corresponding individual targets. Only the top 20% of enhancers by mingap specificity are considered in each target, and only for target pairs or triplets with significantly positive specificity (**Figure 4B** and **E**). For each row, we include 1) the distribution of the number of non-overlapping motifs per sequence (left panel), showing that enhancers targeting multiple cell types used, on average, a similar total number of regulatory elements compared to their single-target counterparts; 2) Motif density (average number of motif occurrences per sequence, middle panel), for the 10 motifs with the highest density in the composite target, showing that most of these were present when targeting the corresponding individual cell types; and 3) motif density in the composite target compared to the average density across individual cell types (right panel). Markers above the diagonal represent motifs that are present at a greater frequency than expected in composite compared to single target enhancers. Motifs with a density in composite enhancers 20% higher than expected (i.e. y > 1.2x) are labeled by name.

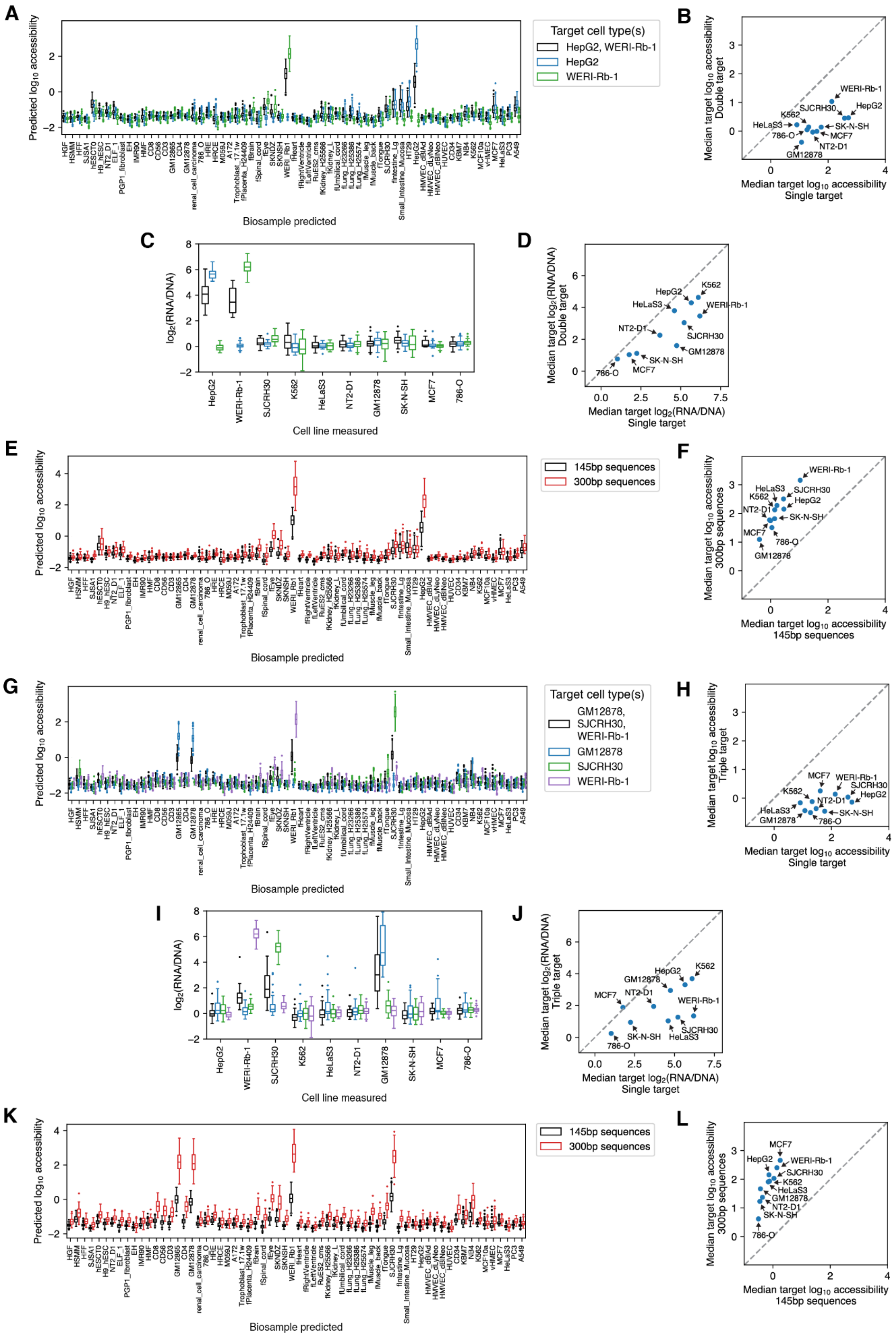

A
Predicted log10 accessibility
Biosample predicted
Target cell type(s)
HepG2, WERI-Rb-1
HepG2
WERI-Rb-1
B
Median target log10 accessibility Double target
Median target log10 accessibility Single target
WERI-Rb-1
HepG2
K562
SJCRH30
HeLaS3
SK-N-SH
786-O
MCF7
NT2-D1
GM12878
C
log2(RNA/DNA)
Cell line measured
HepG2
WERI-Rb-1
SJCRH30
K562
HeLaS3
NT2-D1
GM12878
SK-N-SH
MCF7
786-O
D
Median target log2(RNA/DNA) Double target
Median target log2(RNA/DNA) Single target
E
Predicted log10 accessibility
Biosample predicted
145bp sequences
300bp sequences
F
Median target log10 accessibility 300bp sequences
Median target log10 accessibility 145bp sequences
G
Predicted log10 accessibility
Biosample predicted
Target cell type(s)
GM12878, SJCRH30, WERI-Rb-1
GM12878
SJCRH30
WERI-Rb-1
H
Median target log10 accessibility Triple target
Median target log10 accessibility Single target
I
log2(RNA/DNA)
Cell line measured
J
Median target log2(RNA/DNA) Triple target
Median target log2(RNA/DNA) Single target
K
Predicted log10 accessibility
Biosample predicted
145bp sequences
300bp sequences
L
Median target log10 accessibility 300bp sequences
Median target log10 accessibility 145bp sequences

**Supplementary Figure 20. Enhancers designed to target multiple cell types show lower activity than single-target sequences, likely due to length limitations. (A-F)** focus on synthetic enhancers designed to target two cell types, whereas **(G-L)** focus on enhancers targeting three. **(A and G)** Predicted accessibility for synthetic enhancers targeting the HepG2;WERI-Rb-1 pair **(A)** or the GM12878;SJCRH30;WERI-Rb-1 triplet **(G)** as well as each corresponding individual cell line. Enhancers targeting individual cell types can achieve higher predicted activity than those with composite targets. **(B and H)** Median predicted accessibility on each cell line, for synthetic sequences targeting cell lines individually (x axis) or as part of a double- or triple-target design (y axis). Markers below the diagonal indicate weaker predicted accessibility in sequences designed for a composite target. **(C and I)** Measured enhancer activities of synthetic sequences targeted to the HepG2;WERI-Rb-1 pair **(C)** or the GM12878;SJCRH30;WERI-Rb-1 triplet **(I)** as well as to each corresponding individual cell line. Enhancers targeting individual cell types also drive stronger gene expression experimentally. **(D and J)** Median enhancer activity of sequences designed to target each cell line, either individually (x axis) or as part of a double- or triple-target design (y axis). In **(C)**, **(D)**, **(I)** and **(J)**, only sequences within the top 20% by mingap score per target were considered. **(E and K)** Predicted accessibility for 145 nt and 300 nt-long synthetic sequences targeting the HepG2;WERI-Rb-1 pair **(E)** or the GM12878;SJCRH30;WERI-Rb-1 triplet **(K)**. Longer sequences can achieve higher predicted activities that may restore the performance drop associated with targeting multiple cell types. **(F and L)** Median predicted accessibility on each cell line for double- and triple-target synthetic sequences with 145 nt (x axis) and 300 nt (y axis). Markers above the diagonal indicate higher predicted accessibility in longer sequences.

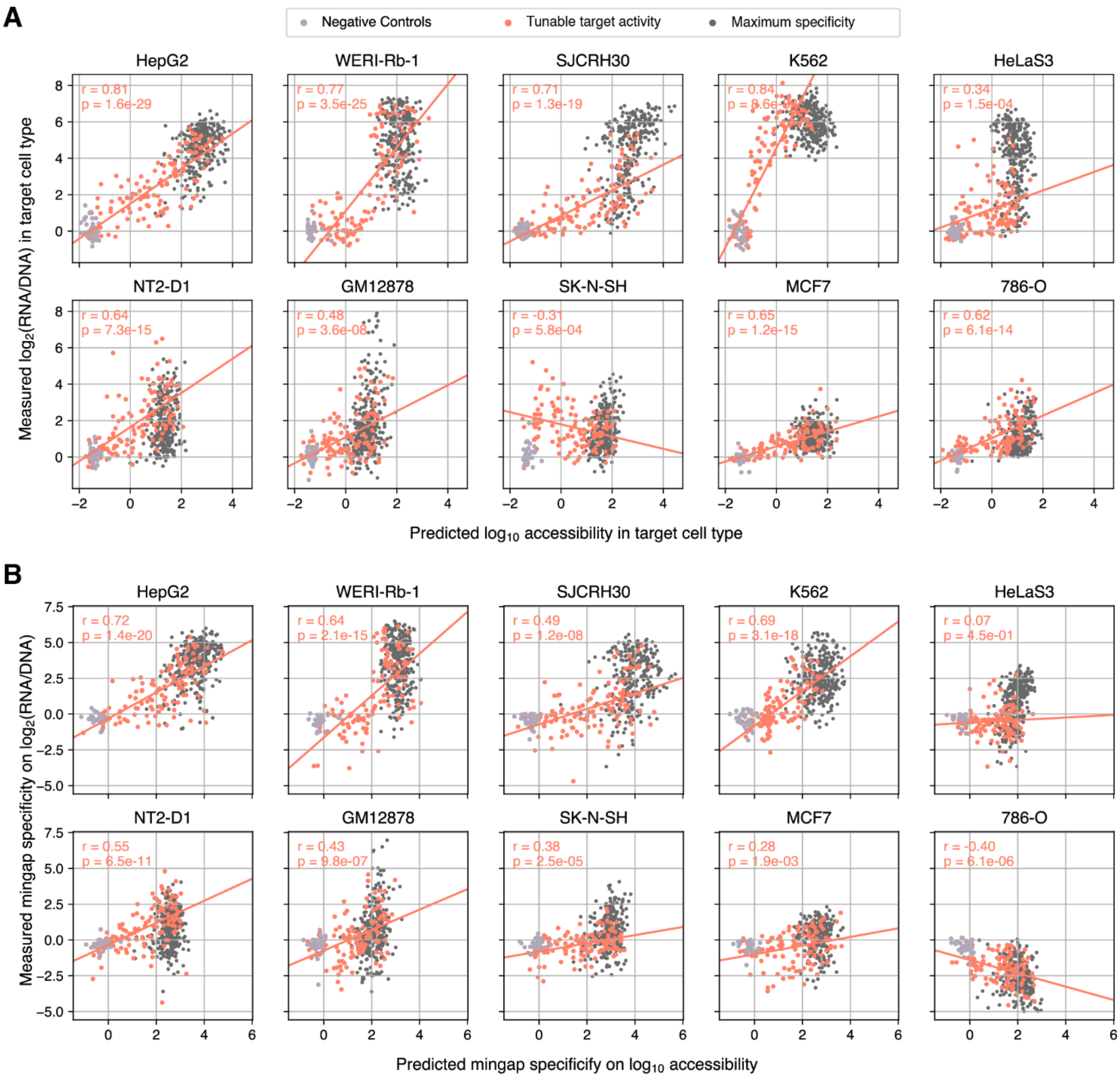


**Supplementary Figure 21. Synthetic enhancers designed for tunable target expression.** Relationship between predicted target accessibility and measured target $log_2$(RNA/DNA) **(A)**, and between DHS64-predicted and observed mingap scores **(B)** for sequences designed for maximal (i.e. **Figure 2**) and tunable target activity, alongside negative controls, for all 10 target cell lines. Mingap scores predictions were calculated from accessibility predictions across the 10 measured cell lines. Observed scores were calculated from $log_2$(RNA/DNA) values as in **Figure 2D**. Linear regression fits, Pearson r coefficients, and p-values were calculated using tunable enhancers only (salmon markers).

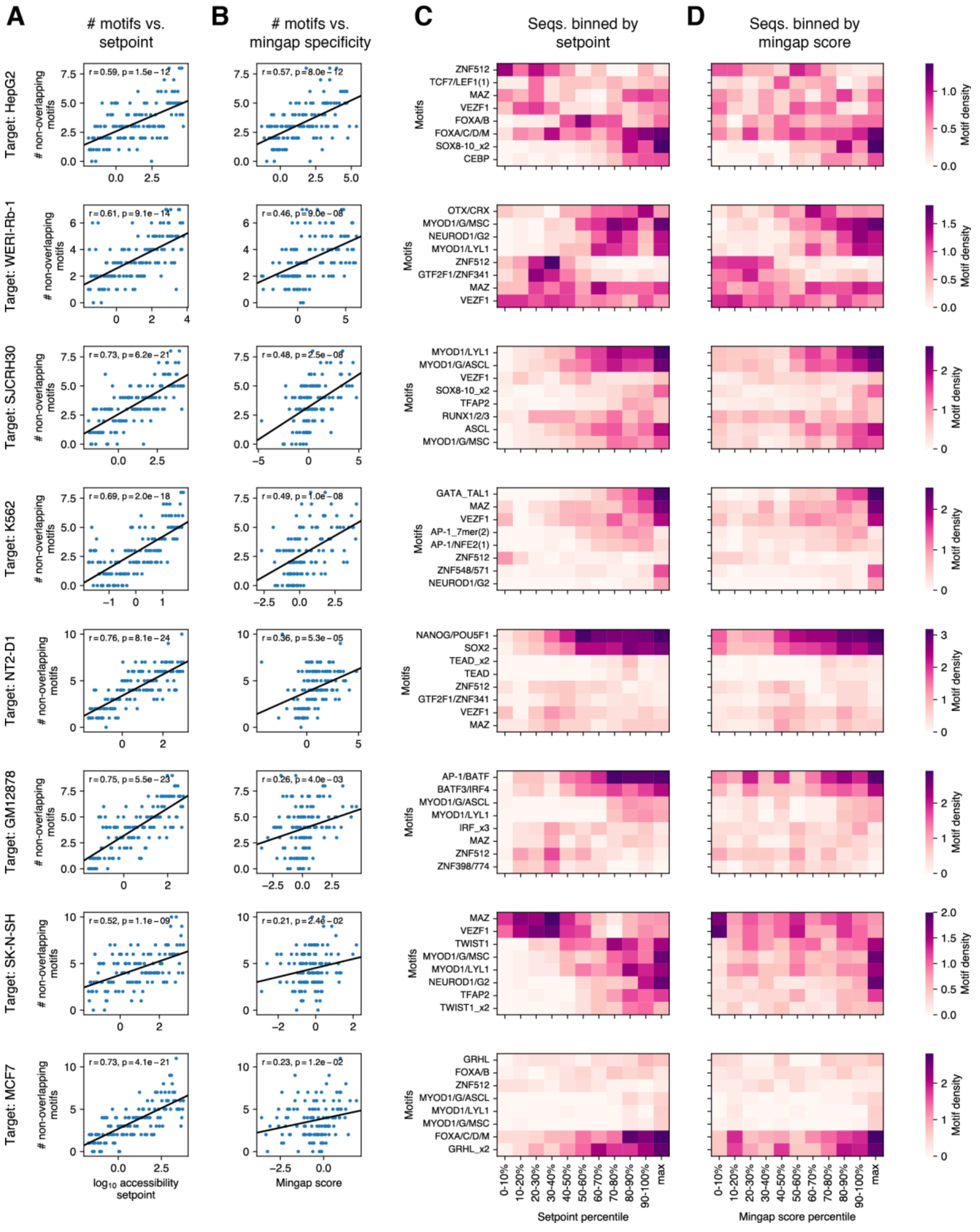


**Supplementary Figure 22. Tunable enhancer activity is achieved via changes in motif density and identity.** Each row shows the motif contents of enhancers designed to target individual cell lines with tunable target activity (**Figure 4I-J**). Only 8 target cell lines where tunability over the mingap specificity score was successful (positive correlation and $p < 0.05$ in **Supplementary Figure 21**) are shown. **(A)** Total number of (non-overlapping) motifs per sequence versus their intended target setpoint, showing that our design algorithm generally achieved intermediate activities via a variable number of regulatory elements. **(B)** Number of motifs versus experimental mingap score, showing that this strategy was generally successful. **(C)** Motif density for enhancers binned by their target setpoint value. Motif density of enhancers designed to maximize specificity (**Figure 2**) is shown in the rightmost column for comparison. While the density of many motifs monotonically increases with setpoint, some are only used at low or medium setpoint values, such as GTF2F1/ZNF341 which is present in low/medium strength SJCRH30-specific enhancers. **(D)** Motif density for enhancers binned by their experimentally measured mingap score, showing that most of the motif usage patterns intended to achieve intermediate setpoints translated to intermediate enhancer activities.

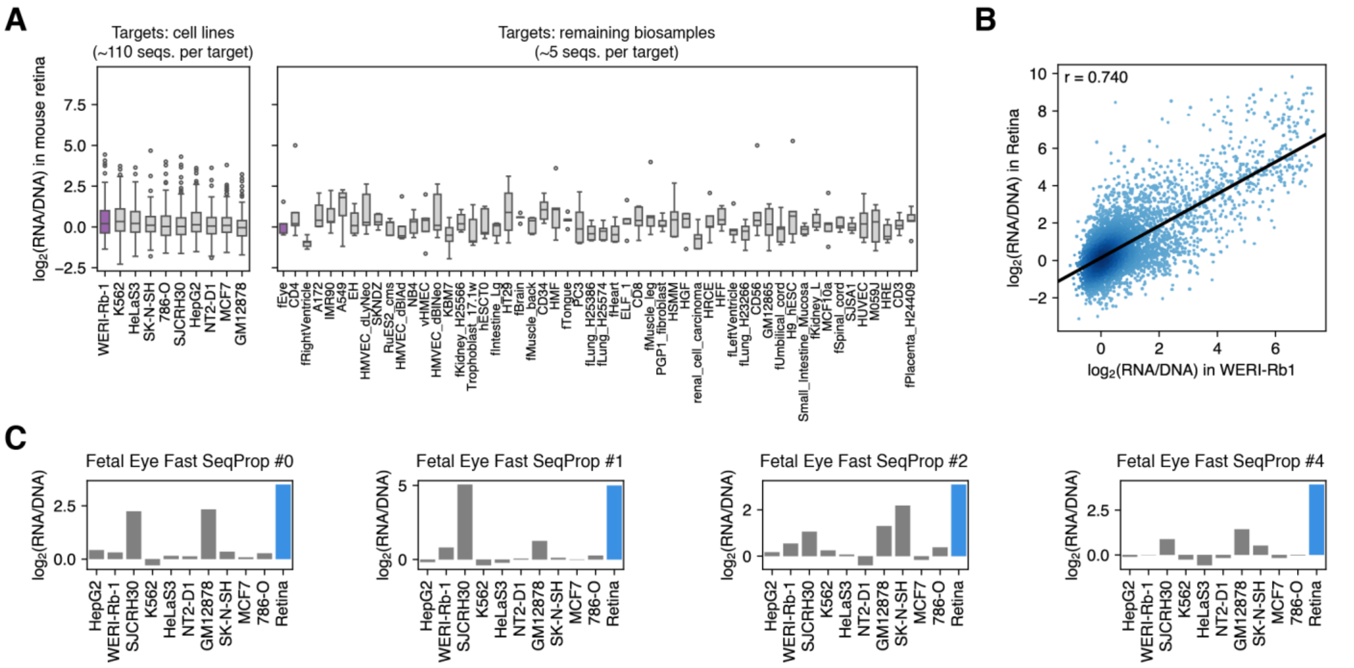


**Supplementary Figure 23. Additional results of mouse retina MPRA experiments. (A)** Measured retinal enhancer activity of DHSs selected for specificity towards all DHS64-modeled biosamples. The range of the y axis and the target biosample order are matched to **Figure 5B**. **(B)** Activities of all single-target enhancers and controls in the retinoblastoma cell line WERI-Rb1 compared to mouse retina. **(C)** Additional enhancers designed to target the fetal eye biosample and tested in cell line and mouse retina MPRAs.

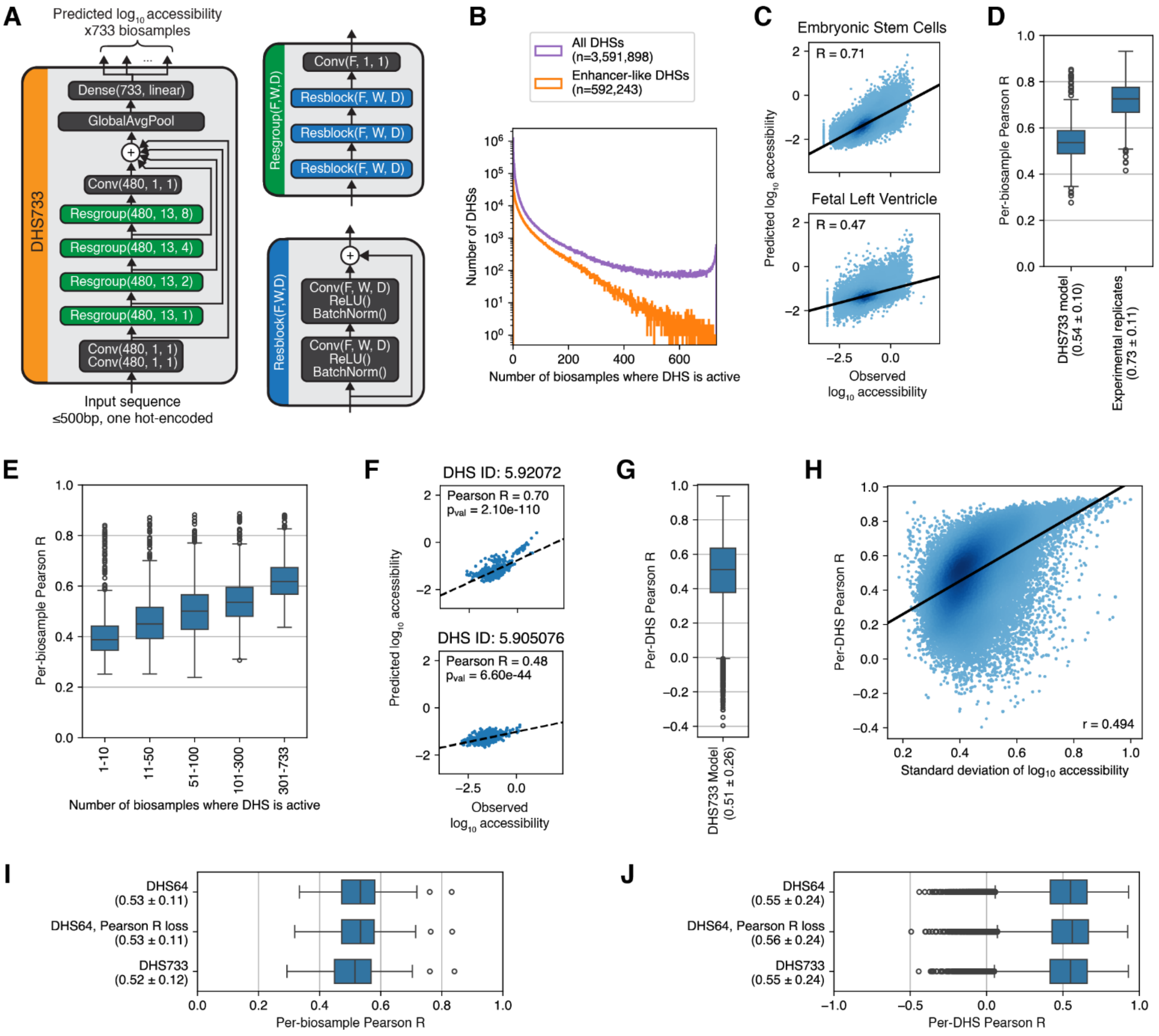


**Supplementary Figure 24. DHS733 model performance.** Results are shown for models trained on chromosome split #3 (**Methods**, **Supplementary Table 2**). Similar performance was observed for two additional models trained on splits 0 and 1. Per-biosample metrics of these models can be found in **Supplementary Table 11**. Numbers in **(D)**, **(G)**, **(I)**, and **(J)** indicate the median ± interquartile range. **(A)** DHS733 model architecture. **(B)** Distribution of the number of biosamples in which each DHS is accessible (i.e. has a positive peak call). Distributions are shown for all DHSs and for "enhancer-like" DHSs (high “mean_signal” annotation, far from annotated transcription start sites, "enhancer" ChromHMM annotations, **Methods**). **(C-H)** DHS733 performance evaluated on DHSs held out from training within the “enhancer-like” set: **(C)** Predicted vs. observed accessibility in two biosamples. Pearson correlation is shown at the top of each panel. **(D)** Distribution of prediction/measurement correlations across all biosamples (left), compared to estimated experimental replicate correlations (right). The latter were calculated by considering biosamples with identical names in the DHS Index as “replicates”. 68 biosamples with unique names (i.e. no “replicates”) were not considered. **(E)** Per-biosample performance on DHSs stratified by the number of biosamples they are active in, from most (left) to least (right) cell type-specific. An equivalent analysis on Enformer[23] reported median correlations of 0.1, 0.27, 0.35, 0.43, and 0.76 for accessible regions active in 0-10, 10-50, 50-100, 100-300, and 300-685 accessibility tracks[24]. **(F)** Predicted vs. observed accessibility across biosamples for two held-out DHSs. **(G)** Distribution of per-DHS Pearson correlations. **(H)** Per-DHS performance versus variability in measured accessibility across biosamples. Performance is better in DHSs that show greater variation. **(I and J)** Direct comparison of DHS733 vs. DHS64 performance on DHSs from the DHS64 test set, which consist of cell type-specific enhancers (**Supplementary Figure 1**).

Because of quantile normalization performed during preprocessing, where accessibility values of each biosample are "normalized" using values of all others, values used for DHS733 differ slightly to those used with DHS64 even for the initial 64 biosamples. Here, we compare predictions to data preprocessed for DHS733, therefore DHS64 performance values differ to those in **Figure 1** and **Supplementary Figure 1**. We additionally include a "DHS64, Pearson R loss" model, which was trained on DHS64 biosamples but using the same per-DHS Pearson R loss and no data filtering as in DHS733, as opposed to the standard MSE loss and extensive training data filtering used with DHS64 (**Methods**). This alternative strategy enabled the use of the full dataset while maintaining performance on cell type-specific DHSs. **(I)** Per-biosample performance. **(J)** Per-DHS performance.

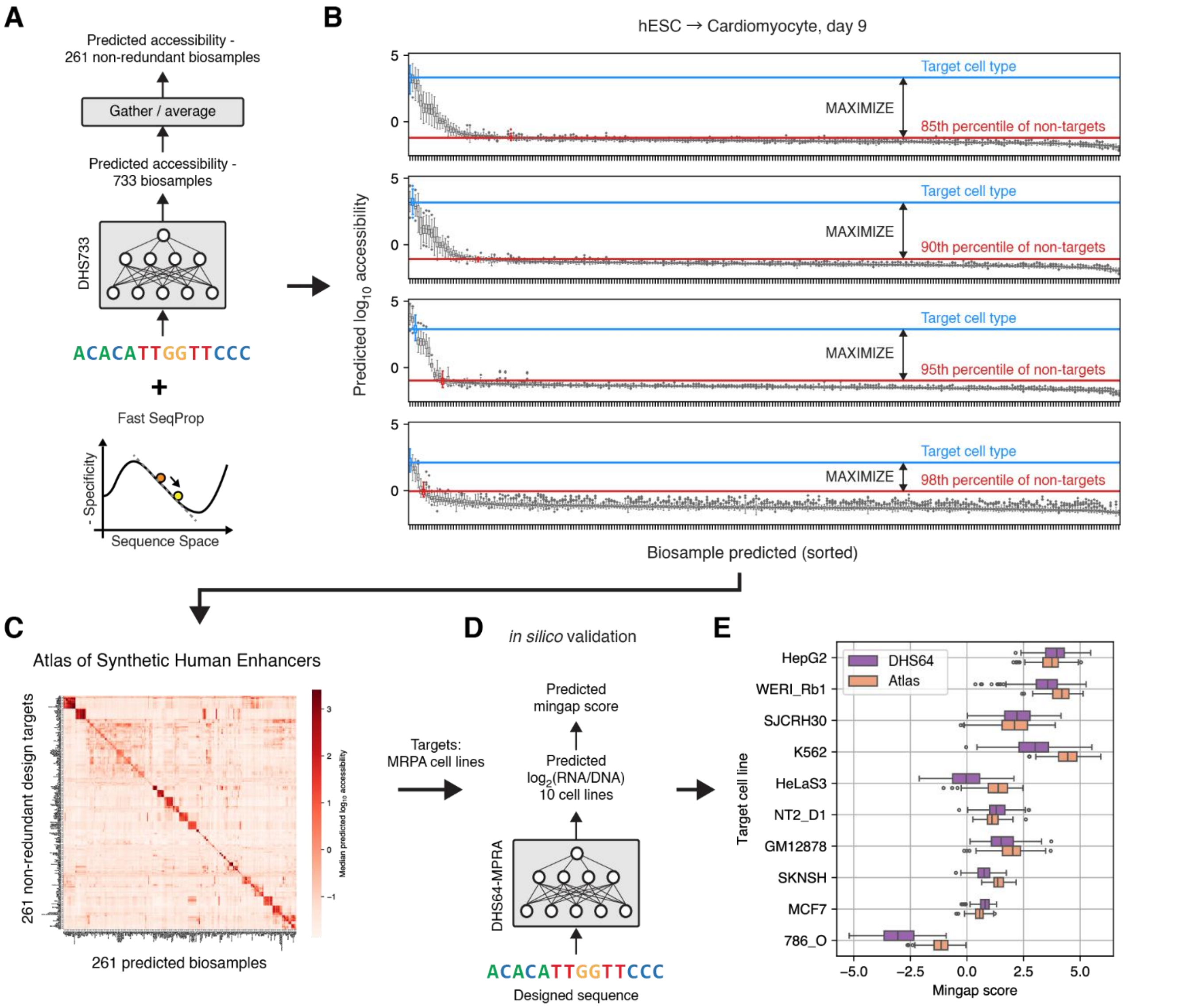


**Supplementary Figure 25. Design workflow and *in silico* validation of sequences in the Atlas of Synthetic Human Enhancers.** **(A)** DHS733 models were modified to aggregate and average predictions across redundant (i.e. identically named) biosamples, resulting in 261 non-redundant outputs. We used Fast SeqProp to generate synthetic sequences targeting every non-redundant biosample. **(B)** Because related biosamples often exhibit correlated accessibility, our previous loss based on penalizing the average of all non-target predictions (**Figure 1E**) can inadvertently penalize the target by suppressing biologically related non-target outputs. To address this, we instead minimized a selected percentile of non-target predictions, and performed four runs with different percentile selections. This resulted in a variety of sequences that trade off high target activity that may co-occur with activity in related biosamples (top) against a more stringent design but with lower on-target activity (bottom). 50 sequences per percentile value per target (total: 200 per target) are included in the Atlas. **(C)** The resulting Atlas of Synthetic Human Enhancers. **(D)** We took sequences from the Atlas targeted to MPRA cell lines (**Figure 2**) and simulated MPRAs by using DHS64-MPRA to estimate enhancer activities and mingap scores. **(E)** Predicted performance of Atlas enhancers against DHS64 Fast SeqProp-designed enhancers. Atlas enhancers have comparable activity and even overcome an issue where DHS64 HeLaS3 enhancers were unsuccessful on average when designed via Fast SeqProp, necessitating training DENs (**Figure 6**).

#### Supplementary Tables

**Supplementary Table 1. DNase I Index biosamples selected for DHS64.**

**Supplementary Table 2. Chromosome-based data splits for DHS64 and DHS733 training.**

**Supplementary Table 3. Prediction performance of DHS64 models on each modeled biosample.**

**Supplementary Table 4. DHS64 DEN training parameters.**

**Supplementary Table 5. DHS64-designed enhancers for all 64 modeled biosamples**

**Supplementary Table 6. Cloning and sequencing oligos.**

**Supplementary Table 7. Cell line culture and transfection conditions.**

**Supplementary Table 8. Sequencing libraries.**

**Supplementary Table 9. MPRA measurements.**

**Supplementary Table 10. Motif clusters.**

**Supplementary Table 11. Prediction performance of DHS733 models on each biosample.**

**Supplementary Table 12. DHS733-designed Atlas of Synthetic Human Enhancers.**

#### References


1. Meuleman, W. *et al.* Index and biological spectrum of human DNase I hypersensitive sites. *Nature* **584**, 244–251 (2020).
2. Abugessaisa, I. *et al.* refTSS: A Reference Data Set for Human and Mouse Transcription Start Sites. *J. Mol. Biol.* **431**, 2407–2422 (2019).
3. Vu, H. & Ernst, J. Universal annotation of the human genome through integration of over a thousand epigenomic datasets. *Genome Biol.* **23**, 1–37 (2022).
4. Linder, J. & Seelig, G. Fast activation maximization for molecular sequence design. *BMC Bioinformatics* **22**, 510 (2021).
5. Linder, J., Bogard, N., Rosenberg, A. B. & Seelig, G. A Generative Neural Network for Maximizing Fitness and Diversity of Synthetic DNA and Protein Sequences. *Cell Syst.* **11**, 49-62.e16 (2020).
6. Grant, C. E., Bailey, T. L. & Noble, W. S. FIMO: scanning for occurrences of a given motif. *Bioinformatics* **27**, 1017–1018 (2011).
7. Yin, C. *et al.* Iterative deep learning design of human enhancers exploits condensed sequence grammar to achieve cell-type specificity. *Cell Syst.* **16**, (2025).
8. Ernst, J. *et al.* Genome-scale high-resolution mapping of activating and repressive nucleotides in regulatory regions. *Nat. Biotechnol.* **34**, 1180–1190 (2016).
9. Martin, M. Cutadapt removes adapter sequences from high-throughput sequencing reads. *EMBnet.journal* **17**, 10–12 (2011).
10. Zorita, E., Cuscó, P. & Filion, G. J. Starcode: sequence clustering based on all-pairs search. *Bioinformatics* **31**, 1913–1919 (2015).
11. Love, M. I., Huber, W. & Anders, S. Moderated estimation of fold change and dispersion for RNA-seq data with DESeq2. *Genome Biol.* **15**, 1–21 (2014).

12. Bravo González-Blas, C. *et al.* SCENIC+: single-cell multiomic inference of enhancers and gene regulatory networks. *Nat. Methods* **20**, 1355–1367 (2023).

13. Castro-Mondragon, J. A., Jaeger, S., Thieffry, D., Thomas-Chollier, M. & van Helden, J. RSAT matrix-clustering: dynamic exploration and redundancy reduction of transcription factor binding motif collections. *Nucleic Acids Res.* **45**, e119 (2017).

14. Jin, H. *et al.* Systematic transcriptional analysis of human cell lines for gene expression landscape and tumor representation. *Nat. Commun.* **14**, 5417 (2023).

15. THE GTEX CONSORTIUM. The GTEx Consortium atlas of genetic regulatory effects across human tissues. *Science* **369**, 1318–1330 (2020).

16. Tarazona, S. *et al.* Data quality aware analysis of differential expression in RNA-seq with NOISeq R/Bioc package. *Nucleic Acids Res.* **43**, e140 (2015).

17. Maslova, A. *et al.* Deep learning of immune cell differentiation. *Proc. Natl. Acad. Sci.* **117**, 25655–25666 (2020).

18. Liu, J. *et al.* Dissecting the regulatory logic of specification and differentiation during vertebrate embryogenesis. 2024.08.27.609971 Preprint at https://doi.org/10.1101/2024.08.27.609971 (2024).

19. Piovesan, A. *et al.* On the length, weight and GC content of the human genome. *BMC Res. Notes* **12**, 1–7 (2019).

20. Gosai, S. J. *et al.* Machine-guided design of cell-type-targeting cis-regulatory elements. *Nature* **634**, 1211–1220 (2024).

21. Yanai, I. *et al.* Genome-wide midrange transcription profiles reveal expression level relationships in human tissue specification. *Bioinformatics* **21**, 650–659 (2005).

22. Lundberg, S. M. & Lee, S.-I. A Unified Approach to Interpreting Model Predictions. in *Advances in Neural Information Processing Systems* vol. 30 (Curran Associates, Inc., 2017).

23. Avsec, Ž. *et al*. Effective gene expression prediction from sequence by integrating long-range interactions. *Nat. Methods* **18**, 1196–1203 (2021).

24. Kathail, P. *et al*. Current genomic deep learning models display decreased performance in cell type-specific accessible regions. *Genome Biol.* **25**, 1–22 (2024).
